# High-yield bioproduction of virus-free virus-like P4-EKORhE multi-lysin transducing particles as an antimicrobial gene therapeutic

**DOI:** 10.1101/2025.01.15.633220

**Authors:** Robert Ramirez-Garcia, Antonia P. Sagona, Jeremy J. Barr, Alfonso Jaramillo

**Affiliations:** Monash-Warwick Alliance Joint Doctoral Program; School of Life Sciences, Faculty of Science, Engineering and Medicine, The University of Warwick, Coventry, United Kingdom; School of Biological Sciences, Faculty of Sciences, Monash University, Melbourne, Australia; Department of Computing, Faculty of Engineering, Imperial College London, London, United Kingdom; Innovation and Translation Hub, Imperial College London, London, United Kingdom; I2SysBio, CSIC-Universitat de València, Paterna, Spain; ACGTx, London, United Kingdom

**Keywords:** high-yield bioproduction, bioprocess engineering, virus-like particles, virus-free particles, transducing particles, antimicrobials, gene therapeutics, A549-model of infection, P4-EKORhE

## Abstract

A description of the construction of the bioengineered P4-EKORhE and a comprehensive method for producing very high yields (up to 10^12^ particles per millilitre) enable the use of virus-like particles to transduce genetically encoded antimicrobials through a combination of synthetic biology and optimised upstream and downstream processing. The final product, a gene-delivered antimicrobial in the form of the multi-lysins cassette, is fully functional before and after packaging within P4-EKORhE particles. The antimicrobial activity of the multi-lysins cassette, characterized by its lysis proteins, was tested *in vivo* in both pure bacterial *Escherichia coli* (*E. coli*) cultures and in a model of infection using A549 immortalised human epithelial tissue cell cultures. This work exemplifies several bioproduction methods and demonstrates how the virology of the P4 and P2 phages can be harnessed to establish a bioprocess for producing transducing particles at very high yields, avoiding contamination by the natural virus while maintaining the antimicrobial effectiveness of the final product.

## Introduction

Harnessing the natural mechanisms that phages use to repurpose their host bacteria’s metabolism to produce viral structural material and package viral DNA for self-replication and propagation is how transducing particles are produced. The most characteristic protein-made parts of phages are the capsid, which packages and protects the genetic material, a tail, which allows the genetic material to be ejected upon infection, and the tail-fibres, which enable specific interaction with a particular bacterial host[1]. For a general schematic of these basic structures, see Figure 1.

**Fig. 1.**
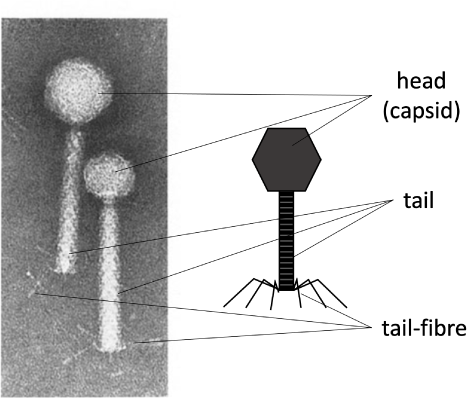
The characteristic protein structures in phages. *Left:* An electron microscope image of phage P2 (with a larger head) and P4 (with a smaller head). *Right:* A schematic labelling of the main protein structures in tailed phages. The capsid (head), the tail, and the tail-fibres. The right side of the image is made by the author. The left side of the image is adapted from the literature[2].

These protein structures allow the phage to store, protect and, upon infection, release the genetic material within the target bacteria. If these protein components can be produced while blocking the virus’s ability to package its own viral DNA, it is possible to produce a transducing particle that contains and delivers a heterologous genetic payload to a target bacterial host. This is particularly straightforward when using phages whose “cos-site” has been identified *e.g*., such as *λ* or P2/P4 phages.

### Virus-free VLP production based on conditional-ly-propagating P4 phage

As described extensively in the literature[2][3][4][5], the phage P4 is a helper-dependent phage that uses the late gene products of temperate phage P2 to encapsulate its own P4 double-stranded DNA. The P4 capsid is about one-third the volume of the P2 capsid[6] and harbours a genome also one-third its size (*i.e*. 11.624 kb versus 33.574 kb, respectively). As aforementioned, the P4 phage overexpresses of the P2 prophage late genes[4] without the P2 prophage being excised or replicated [7]. Another characteristic of P4 is that, when a P2 prophage is not present in the host bacterial cell, P4 can still be maintained as a plasmid[8], which allows P4 to hijack a subsequent infection by a virulent P2 phage.

Therefore, the viral cycle of phage P4 is a unique phenomenon in which one phage parasitizes another to perpetuate itself, repurposing the gene products of the helper phage. While the phage P2 constructs an icosahedral capsid (head) with T = 7 symmetry using the *gpN* capsid protein, the *gpO* scaffolding protein, and the *gpQ* portal protein, the presence of P4 drives these structural proteins into a smaller capsid with T = 4 symmetry. The regulation of this size transformation is facilitated by the P4-encoded protein *Sid* which creates an external scaffold around the compact P4 procapsids[9]. This capsid reduction restricts P2’s larger genome from propagating while favoring packaging of the smaller P4 genome.

The P4 phage viral cycle is described in the literature[10]. P4, a satellite virus, relies on the helper phage P2 for propagation. The cycle begins when P4 infects a susceptible bacterial host, such as *E. coli*, alongside P2. During infection, P4 utilises the P2 machinery and structures to replicate its genome and produce viral proteins. Notably, P4 can also exist in a plasmid form within the host cell, replicating independently of P2. This plasmid state allows P4 to remain within a bacterial population even without P2. When conditions become favorable, such as when P2 infects the same host, P4 integrates into the P2 life cycle, co-packaging its genome with P2 during lytic growth and releasing new virions containing both P2 and P4 genetic material. This interplay lets P4 exploit P2 resources while preserving its unique life cycle traits.

Since P4 can be maintained as a plasmid and depends on P2 for conditional propagation (*i.e*. using P2’s late structural proteins for packaging and storage), it is feasible to use this phenomenon to produce transducing particles via the P4’s conditionally propagating capability. Moreover, a minimal essential region of P4 has been described[11][12]. One can thus engineer a “conditionally-propagating transducing particle” with a P4-minimal construct, here referred to as “P4-min”, measuring 7.190 kb and allowing expression of the necessary gene products for fully functional P4 plus a foreign genetic circuit of up to 4.249 kb. This capacity parallels the full-size 11.439 kb of P4. A P4-min-based system can host a genetic device encoding antimicrobial-like molecules. This work illustrates a system that includes *the multi-lysin cassette*, a kanamycin-resistance marker, and a co-located P4-cos site spanning 4.151 kb. In this context, P4-min functions as a transducing particle capable of propagation only in the presence of P2. It thus reaches yields typical of natural phages (≥ 10^8^ cfu/mL).

Here, a transducing particle is defined as a viral particle that carries a plasmid encoding heterologous or synthetic genetic material, along with a “cos-site” acting as a “packaging signal”, instead of the phage’s own genome. Figure 2 summarizes how P2/P4 phages can generate transducing particles.

**Fig. 2.**
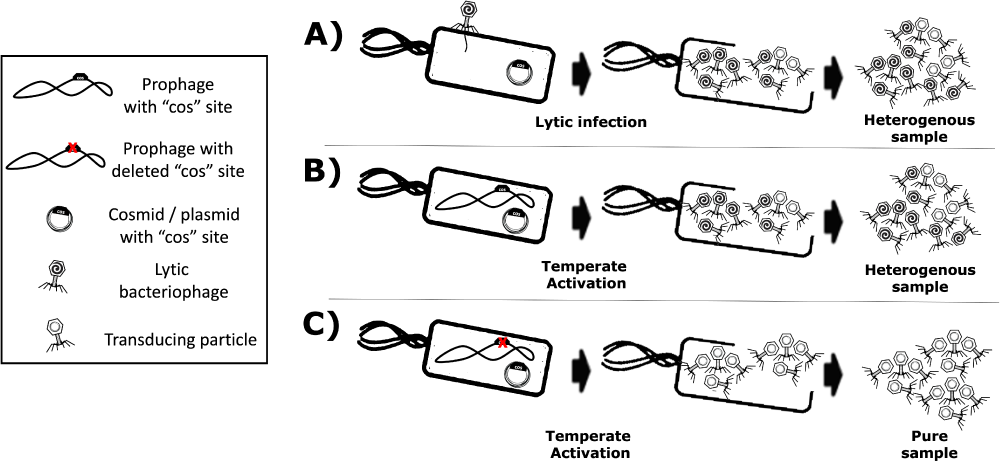
Production of transducing particles using lytic or temperate phages. Transducing particles are phage-based particles harbouring cosmids instead of phage genomes, which have been selected and maintained within host-producing strains. **A)** Lytic phage production yields a mixture of phages and transducing particles. **B)** Temperate phages with functional cos-site also yield a mixture of phages and transducing particles. **C)** A temperate phage with a *knocked-out* cos-site yields a “virus-free” pure sample of transducing particles.

The main disadvantage of using helper P2 phages for transducing-particle production is contamination of the lysate with wild-type phages, which are self-replicating and highly propagative. Such contamination can arise from non-adsorbed helper phage and from packaging of competing viral genomes. Because the virus’s DNA and the proteins needed for virion assembly are produced in the host cell, both the cosmid and the viral genome compete for packaging, yielding a mixture of transducing particles (plasmid/cosmid) and viruses (viral genome). Figure 2.A and Figure 2.B, show these scenarios.

Using helper P2 prophages (*i.e*. lysogens) to produce transducing particles presents another drawback: the spontaneous induction of lysogenic cells into a lytic cycle occurs at a low frequency[13]. Temperate strains can be activated chemically[14], or by UV-light[15][16], but each activator is phage-specific. For example, UV-light can induce *λ* lysogens, but not P2 lysogens.

In this regard, the epsilon (*ε*) gene product from natural P4 phage activates temperate P2 prophages[17]. P4 is a satellite phage that predates on P2 phage late gene products as a “helper” for its construction and lytic growth[10]. Such “late genes” contribute to the production of the aforementioned phage structural proteins such as the capsid, the tail, and tail-fibres (Figure 1) for its complete constitution as a virus[18]. In this light, *transactivation of P2 late gene expression by P4 requires the P4 delta (δ) gene product and works even without P2 DNA replication*[19] benefiting P4 viral genome packaging over competing P2.

Hence, P2-lysogenic strains can help produce phage-based transducing particles because inducers have been identified, and cosmid packaging can outcompete native viral genomes. Furthermore, temperate lysates become virus-free (Figure 2.C) by *knocking out* the cos-site of the temperate strain’s genome, ensuring only foreign cosmids (with the intact cos-site) get packaged.

In this work, the P4-EKORhE, a virus-free virus-like transducing particle production system based on P4’s ability to hijack the P2 prophage’s late structural proteins, is shown to deliver a multi-lysins cassette as a gene-delivered antimicrobial.

### Phage lysins as a gene-delivered antimicrobial

The multi-lysins cassette is a combination of genetically encoded lysins designed for delivery via transducing particles. Phage lysins and lysis-accessory proteins (*e.g*. holins, spanins) are natural effectors that disrupt bacterial membranes or cell walls to release phage progeny. Because these proteins typically target bacterial membranes and cell walls, they resemble membrane-disrupting antibiotics by compromising bacterial viability[20].

In this sense, *E. coli* K12 is used as a target because of its low biosecurity risk and its role as a model for more pathogenic strains, including *E. coli* K1, which can cross the blood-brain-barrier in neonatal meningitis and is a known uropathogen[21]. Combining multiple lysins and lysis-accessories can strengthen antimicrobial effects, as seen previously with extracellular lysins[22]. Here, however, the lysins are genetically encoded and expressed intracellularly in the target bacterium. Both, MS2 gpL and PhiX174 gpE single-gene lysis proteins, and the *λ* lysis system (LysS, LysR, Rz) are used within a single cassette (Supplementary Section SS4.4). Figure 3 and Figure 4 illustrate how these components function synergistically.

**Fig. 3.**
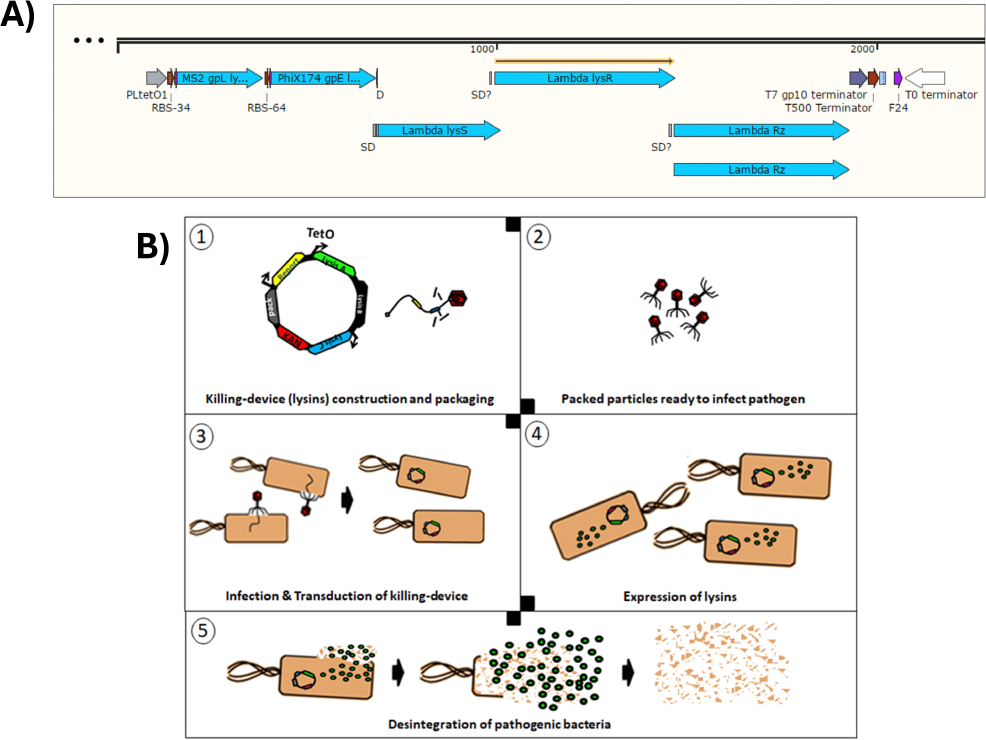
Conceptual scheme of the P4-EKORhE multi-lysin antimicrobial. **A)** The multi-lysins cassette harbouring all lysins and lysis-accessory proteins. The different encoded lysis and lysis accessory proteins used to construct the multi-lysins cassette. A study of the functionality of these circuits is presented in Figure 4. The “SD” and “SD?” labels contained within the construct are predicted “Shine-Delgarno” sequences that correspond to the multi-cistronic nature of the *λ* phage lysis operon n (*lysS, lysR* and *Rz*) and the likely presence of unidentified *cryptic promoters*. **B)** Conceptual scheme of the P4-EKORhE multi-lysins as a genetically-encoded antimicrobial therapeutic.

**Fig. 4.**
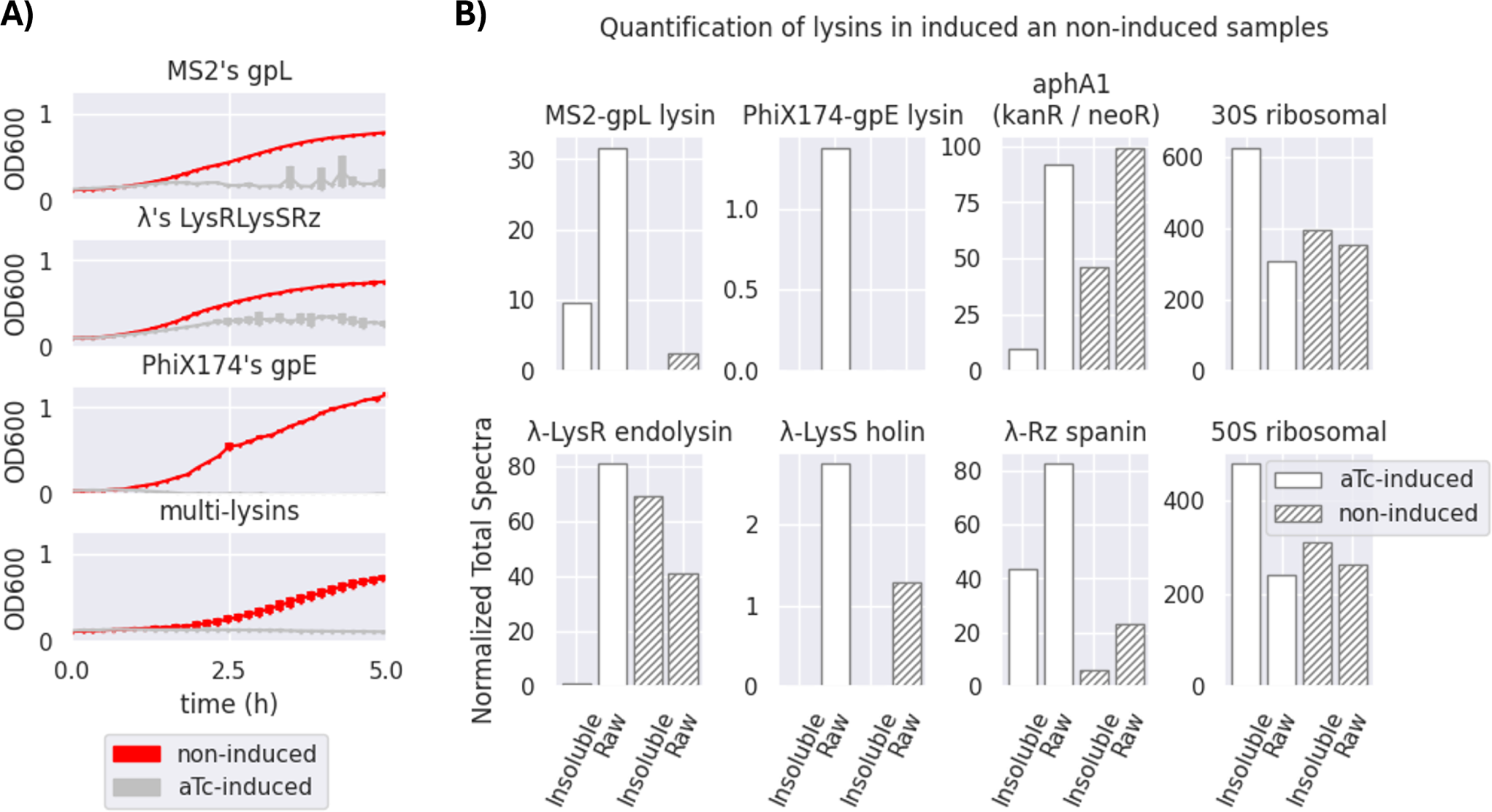
Effectiveness of a transformed multi-lysins cassette. **A)** Optical density (OD600) measurements of bacterial survival profiles in pure bacterial cell cultures of *Escherichia coli* DH5*α* Z1 showing the antimicrobial effect of the multi-lysins cassette (the bottom-most plot) and each of its contributors (MS2’s gpL, *λ*’s lysis cassette, and PhiX174’s gpE, the top-three-to-bottom plots). The “red lines” represent the bacterial cells harbouring the multi-lysins cassette that have not been induced with anhydrous tetracycline (aTc). The “silver lines” represent bacterial cells harbouring the lysins cassette that have been induced with anhydrous tetracycline at the start of the culture. *\* A second replica is available in Figure S1 of the Supplementary Section SS2 for Replicas. The thickness of each line represents variability among the “n” samples using the standard deviation, where n = 3*. **B)** Normalised Total Spectra (NTS) obtained from a Mass-Spectrometry (MS) analysis of 4 different samples (2 induced and 2 non-induced, raw- and insoluble-phase) of bacteria containing the multi-lysins cassette. The Normalised Total Spectra includes data that computes the peptides found with over 95% probability that belong to each of the lysin sequences studied. **aTc-induced:** The protein extract comes from bacteria for which cosmids containing the multi-lysins cassette have been activated by adding aTc (anhydrous tetracycline) to the culture media. **Non-induced:** The protein extract comes from bacteria for which cosmids containing the multi-lysins cassette have not been externally induced with aTc. **Insoluble fraction:** Protein fraction extracted from the pellet of the lysed cells. **Raw fraction:** Protein fraction extracted from samples containing both, the supernatant and the pellet, of lysed cells. For more details see the Supplementary Section SS3.

In this work, a rhamnose-inducible, controllable, and conditionally propagating phage, the P4-EKORhE (Figure 5), is utilised to encode, within its genetic cargo capacity, which is only limited by the *headful* size of its original genome, the multi-lysins cassette, as a means to test the use of a gene-delivered antimicrobial. The multi-lysins cassette is then characterized for its effectiveness as an antimicrobial (Figure 4), and a model of infection both using pure bacterial cell cultures (Figure 7.C) and a human model of bacterial infection using A459 immortalised human epithelial cells (Figure 7.D and 7.E) is used to further ascertain the effectiveness of gene-delivered antimicrobials using P4-EKORhE.

**Fig. 5.**
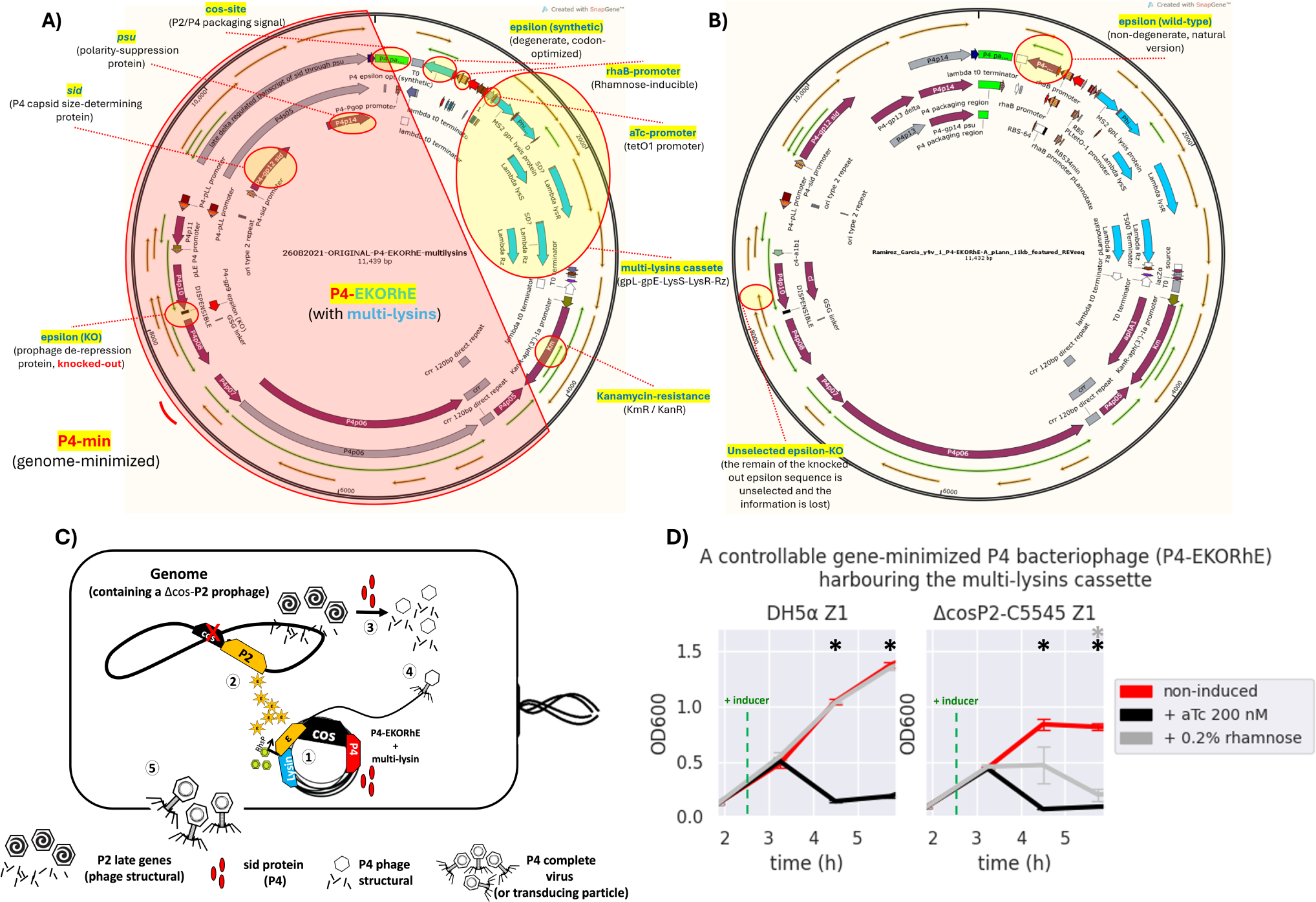
The bioengineered P4-EKORhE harbouring the multi-lysins cassette. **A)** A kanamycin-selectable cosmid of 11,439 bp containing a genome-minimised P4 phage (P4-min, see highlighted in red) that has a knocked-out-version of the epsilon (*ε*) gene, an alternative functional synthetic, codon-optimised, and degenerate version of the *ε* gene, and the multi-lysins cassette harbouring the 3 systems of lysis form MS2, PhiX174 and *λ* phages. **B)** The same cosmid after a few passages loses the dysfunctional *knocked-out wild-type* version of the *ε* gene, while the synthetic degenerate version gets restored to the sequence observed in the wild-type *ε* protein commonly seen in the natural P4 phage (see red circles for “unselected epsilon-KO” and “epsilon (wild-type)”). The stable version of the P4-EKORhE is now 11,432 bp. Fully featured sequences of **A)** and **B)** can be found in the Supplementary Sections SS8.5 and SS8.6, respectively. **C)** A conditionally-propagating, and rhamnose-inducible, P4 phage as a transducing particle. **(1)** The P4-EKORhE is replicated as a cosmid harbouring the multi-lysin cassette, and expresses the *Sid* protein upon activation. **(2)** a Δcos version of the P2 prophage is inhibited by the activity of P4 but activates its late genes, with the delta (*δ*) gene product, which are the necessary structural proteins for P2. Additionally, the rhamnose-inducible *ε* gene product activates the self-replication of the Δcos version of the P2 prophage. **(3)** The *Sid* protein from P4 converts P2 heads into smaller P4 heads, thus further compromising P2 propagation. **(4)** P4 structural proteins package the P4 cosmid harbouring the multi-lysins cassette (*i.e*. or any other engineered circuit), or any other accessory cosmid. **D)** Dynamics of of the activation of the P4-EKORhE-multi-lysins cosmids with inducer, aTc or rhamnose, in strains of *Escherichia coli*, transformed on *Escherichia coli* DH5*α* Z1, or the engineered lysogen Δcos:TriR-P2-c5545 Z1, respectively.The sample size (n) used is n = 3, * = *p-value* < 0.05, measured at the data points used after the addition of the inducer, using a standard t-test using SciPy libraries (see General Methods SS6.1).

**Fig. 6.**
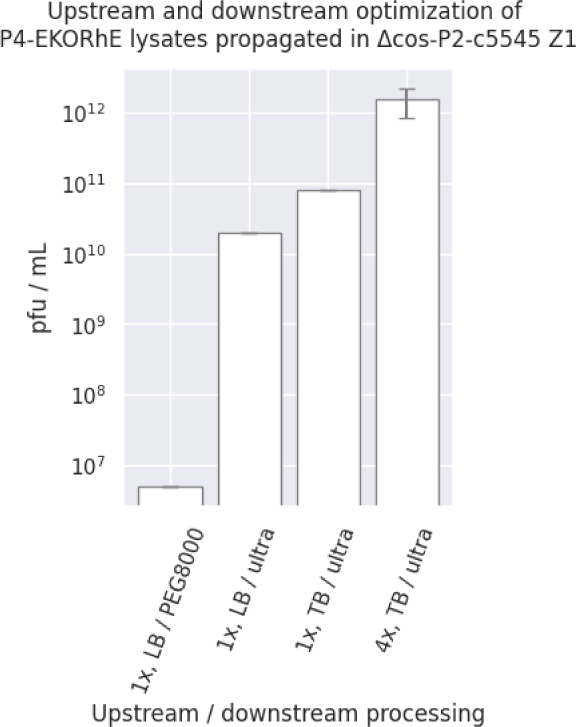
Upstream and downstream optimization of P4-EKORhE. Different yields are obtained when using P4-EKORhE according to the upstream and downstream optimization. In regards to upstream optimization, lysates are produced using either one or four enrichment cycles (*i.e*. 1x or 4x), using LB or TB broth. In regards to downstream optimization, lysates are concentrated using either 10% PEG8000 or molecular weight cut-off ultrafiltration. The plaque-forming units are taken from the plates shown in Figure S4. The sample size (n) used is n = 1, except for “4x, TB / ultra” which has n = 3.

**Fig. 7.**
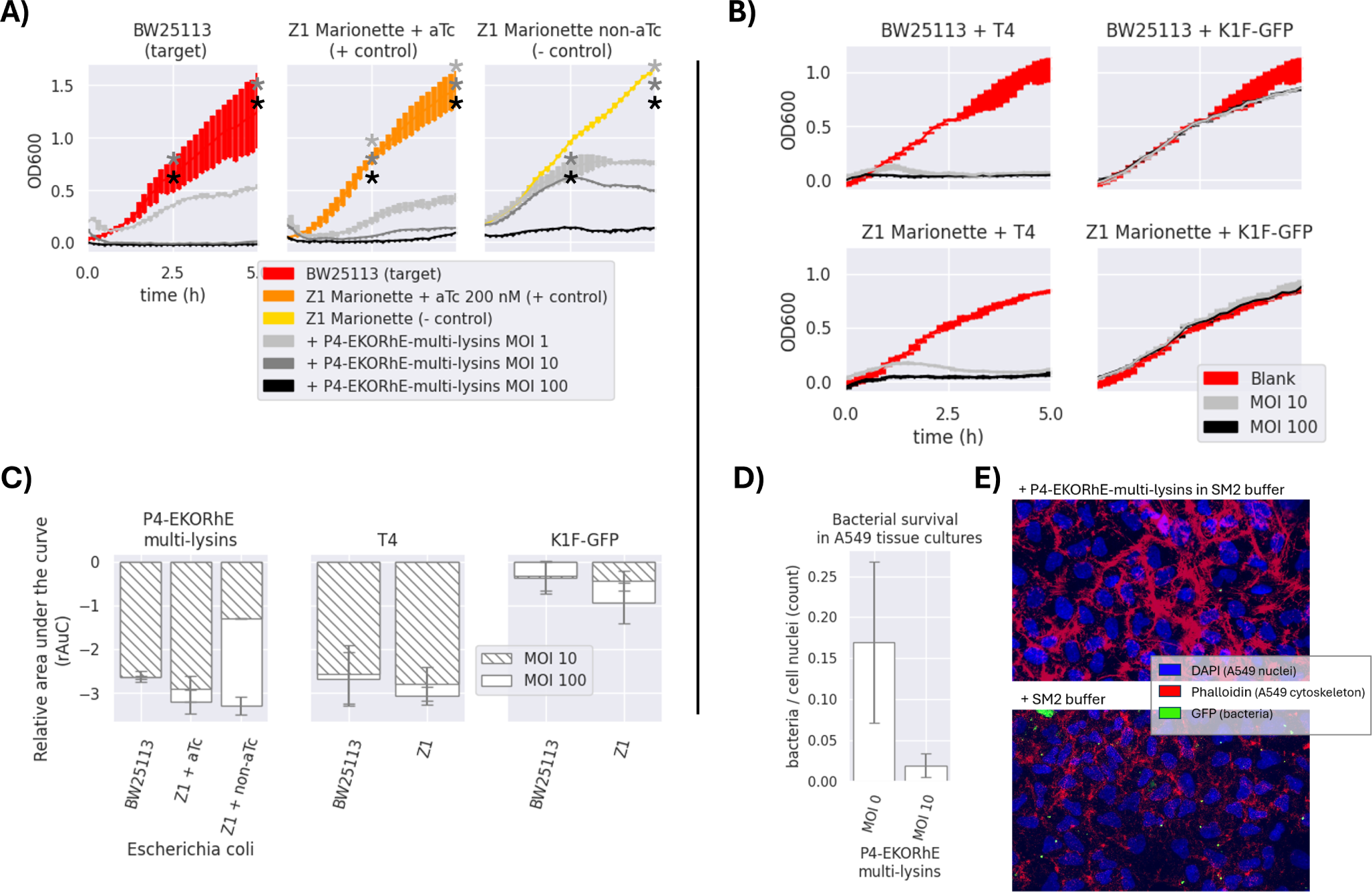
A) Antimicrobial activity of P4-EKORhE-multi-lysins. Optical density measurements of the target and control *Escherichia coli* strains *i.e*., BW25113 and Z1 Marionette, respectively, when exposed to P4-EKORhE-multi-lysins particles at different MOIs. No selection antibiotics were used for the realization of these experiments. **B) Antimicrobial activity of natural enterobacteriophages to its respective hosts**. Antimicrobial activity of enterobacteriophages to its respective target and control E coli strains *i.e*., BW25113 and Z1 Marionette when exposed to natural enterobacteriophages containing a replicative virus at different MOIs. For a replica of the same set of experiments see Figure S2 in the Supplementary Section SS2 for Replicas. The thickness of each line represents variability among the “n” samples using the standard deviation, where n = 3.**C)** Relative area under the curve of P4-EKORhE-multi-lysins, T4 and K1F-GFP at MOIs of 10 and 100 on *E. coli* strains BW25113 (target) and Z1 Marionette (+/− control). rAuC(MOI = 0) = 0 *±*1. The sample size (n) is n = 2 from the results obtained in the two different replicas in Figure 7 and Figure S2 which contains n = 3 for each measure of time (total n=6). The rAuC has been integrated from 0 to 5 hours in all cases.. **D)** The differences in bacterial count following treatment with P4-EKORhE harbouring the multi-lysins cassette on A549 cell tissue at an MOI of 10 versus with a blank solution (MOI = 0) of SM2 buffer, following an immunohistochemistry fluorescence image analysis. **E)** Immunohistochemistry fluorescence image analysis on a sample infected with GFP-fluorescent bacteria treated with P4-EKORhE harbouring the multi-lysins cassette versus the blank solution of SM2 buffer. The sample size (n) used is n = 7 for the (- control), while n = 6 for the target was used. The *p-value* for the difference between the two samples is p = 0.31. The images used to count for the results shown in **D)** can be found in the Supplementary Section SS5 for Immunohistochemistry imaging.

In this light, one of the novelties of the proposed approach is applying the concept of searching synergies among different lysins and lysis-accessory proteins that act as they are meant to act: intracellularly, taking advantage of transduction as a gene-delivery vehicle for the intracellular expression of effector proteins (*i.e*. the lysins, and lysis-accessory proteins, showcased in this work).

In Figure S11 is shown the first version of a functional cosmid, encoding the multi-lysins cassette, with a strong anhydrous-tetracycline (aTc)-inducible promoter. In Figure 3 a conceptual scheme of how the P4-EKORhE multi-lysins is employed as an antimicrobial therapeutic.

## Results and discussion

### R1. The multi-lysins cassette

There is an ample variety of promoters available of diverse strengths in *E. coli*[23]. In this work, a strong aTc-repressible promoter[24], for tight and strong induction of the multi-lysins cassette, and a weaker, rhamnose-inducible promoter[25][26], to control P4-EKORhE transducing particle production, are used.

Given that the multi-lysins cassette is delivered via P4-EKORhE transducing particles, a sufficiently strong promoter is essential during production and packaging of this genetically-encoded antimicrobial (*e.g*. the tetO1/aTc-inducible promoter used in Z1 cells). The tetO1/aTc-inducible promoter (PLtetO1 in Figure 3.A) represses the encoded genetic circuit in “production cells” harbouring the tetR repressor. These cells can safely produce high titers of transducing particles ready to package P4-EKORhE cosmids encoding the multi-lysins cassette without risk of premature activation. Figure S11 shows the first functional version of the multi-lysins cassette under the tetO1 promoter, also used in the high-yield P4-EKORhE that appears in Figure 5.

The multi-lysins cassette in this work combines three lysis systems from PhiX174, MS2 and *λ* phages[27][28][29][30][31] (Supplementary Section SS4.4), expressed under a single promoter and sized appropriately for packaging in P4-EKORhE. Figure 4 demonstrates the effectiveness of each lysin and their combined bactericidal activity. PhiX174 gpE is the strongest contributor to multi-lysins synergy. Figure 4 shows that the activation of lysis results in an antimicrobial effect, as observed over a 5-hour period. This suggests that the multi-lysins cassette is a strong alternative to natural phages.

All three systems of lysins (MS2, *λ*, and PhiX174) seem to contribute noticeably to the multi-lysins cassette, with PhiX174 gpE being the most outstanding contributor. Proteomics analysis of lysins/lysis-accessory proteins in induced and uninduced samples of the multi-lysins cassette is shown in Figure 4.B. Four samples were analysed: two induced with aTc (both soluble and insoluble fractions, as well as insoluble-only fractions), and two uninduced samples (labelled as non-induced in the figure) also for both raw and insoluble fractions.

Figure 4, shows the Normalised Total Spectra (NTS) of peptides corresponding to the lysins and lysis-accessory proteins targeted in this work, with kanamycin-resistance aphA1, 30S and 50S ribosomal proteins acting as positive controls for cosmid-borne and genome-borne expression. Supplementary Section SS3 further detail peptide coverage for these MS samples (Figure S5 and Figure S6).

Figure 4 confirms the presence of cryptic promoters and Shine-Delgarno (SD) sequences in the *λ*-lysis cassette (Figure 3.A and Figure S7). Peptides from *λ*-lysis cassette were detected in both induced and non-induced samples, with a important increase observed upon activation with aTc. This indicates that the *λ*-lysis cassette and MS2 gpL have moderated but detectable activity even before induction, likely due to the accumulation of gene products (LysS holin/antiholin, Rz and Rz1 spanins, and the LysR endolysin) over time. Both MS2 gpL and PhiX174 gpE appear only in aTc-induced samples, consistent with the absence of cryptic promoters or SD sequences upstream of their respective coding sequences (Figure S7).

The constitutive expression of LysS and LysR proteins, as well as Rz, can be explained by the presence of a cryptic promoter upstream of the potential SD-like sequence “GGAAGGAG” at the −3 position of PhiX174 gpE, or from upstream cryptic promoters in the sequences *lysS* and *Rz* (found exclusively in uninduced samples) (Supplementary Section SS8). Figure 4.B confirms cryptic promoters and SD sequences, evidencing their presence when the cosmid is not activated with aTc. However, due to the minimal contribution of these cryptic elements (especially for PhiX174 gpE), they do not impede the system’s use for transducing-particle production, as shown in Figure 5.C. In contrast, MS2 gpL and PhiX174 gpE are only present in aTc-induced samples, supporting their reliance on induction.

The hypothesized mechanism for the synergistic lysis dynamics of the multi-lysins cassette is as follows (refer to Supplementary Section SS4.4 for a comprehensive discussion): (1) *λ*’s LysS[32] accumulates in the cytoplasm and, upon reaching critical concentration, forms pores in the cytoplasmic side of the bacterial membrane[33]. (2) This pore-formation facilitates the intrusion of *λ* LysR endolysin[34], along with single-gene lysins MS2’s gpL[35] and PhiX174’s gpE[36], which disrupt the membrane[37][38]. (3) *λ*’s Rz[39] and Rz1[40] then deliver the final blow after the membrane is disrupted[41][42]. This lysis is observed in non-aTc-induced (uninduced) samples (from cryptic promoter/SD sequences allowing *λ*-lysis cassette expression), suggesting an automatic activation of lysis when accumulation of gene products occur.

The results confirm that the activation of the multi-lysins cassette with aTc corresponds to the presence of lysins, lysis-accessory proteins, and antimicrobial activity as shown in Figure 4.

### R2. The conditionally-propagating P4-EKORhE

The virus-free virus-like, conditionally propagating, P4-EKORhE (a *latinised* transcription for P4-ε KO-Rh ε) which literally stands for “*P4 epsilon (ε) knock-out rhamnose-inducible epsilon (ε)*”, is a bioengineered rhamnose-inducible P4-based system for the high-yield bioproduction of transducing particles presenting itself as a genome-minimised P4 (P4-min) phage with its *ε* gene *knocked-out*, and a degenerate, codon-optimised *ε* expressed elsewhere under the control of a rhamnose-inducible promoter (Figure 5), which is in this work used to harbour the multi-lysins cassette (Figure S11).

During construction, an essential part of the P4 phage[11] (from P4p05 to P4s04, Figure 5.A or Supplementary Section Section SS8) was PCR-amplified and cloned together with the multi-lysins cassette to confer a selection marker (kanamycin resistance). Following successful selection with kanamycin, the genome-minimised P4 (P4-min) was further amplified by PCR to produce a *knock-out* in the native *ε* gene coding sequence and to insert a codon-optimised *ε* that cannot recombine with the native gene.

The whole *ε* gene, the target for the *knock-out*, was not fully removed; instead, a minimal deletion was introduced to avoid potential disruption of P4 functionality. Ultimately, a final mutant was obtained with a stably *knocked-out ε* gene and inserted a construct containing a degenerate version of the *ε* gene downstream the cos-site—located away from the original position of the *ε* gene—under a rhamnose-inducible promoter. Therefore, P4-EKORhE differs from a natural P4 phage because it is a cosmid that (1) remains genome-minimised, (2) carries a selection marker, and (3) incorporates two *ε* gene sequences (with the native epsilon being nonfunctional and the inserted version fully functional), thereby preventing recombination.

Figure 5, displays the final construct of P4-EKORhE harbouring the multi-lysins cassette. The P4-EKORhE genetic sequence was confirmed by analysing trace sequences from the assembled P4-EKORhE (Figure 5.A and 5.B), and the full-sequence of the “stabilised form” of P4-EKORhE was obtained via whole-plasmid sequencing (Figure 5.B); both sequences appear in Supplementary Sections SS8.5 and SS8.6, respectively.

Figure 5.C. illustrates a scheme that shows how the controllable, gene-minimised P4-EKORhE produces transducing particles. The figure demonstrates that the system activates upon induction because P4-EKORhE encodes a synthetic *ε* coding sequence downstream a rhamnose-inducible promoter.

Figure 5.D depicts the dynamics of P4-EKORhE carrying the multi-lysins cassette. The data show that this production system represses phage production when rhamnose is absent (while maintaining 50 *μ*g/mL kanamycin in the culture) and activates fully when 0.2% (w/v) rhamnose is added. As expected, the P4-EKORhE propagates only in Δcos:TriR-P2-c5545 Z1 cells, since no *ε* gene activation is observed in DH5*α* Z1 cells upon rhamnose treatment. Instead, P4-EKORhE remains repressed in the absence of rhamnose and becomes fully inducible when 0.2% (w/v) rhamnose is added. Moreover, a notable drop in OD_600_ is observed upon addition of aTc to the production culture due to the controlled activation of the multi-lysins cassette.

Figure 5.D demonstrates that although repression of P4-EKORhE is not absolute, it sustains a stable culture suitable for high-yield bioproduction. This stability allows to maintain P4-EKORhE, alongside other cosmids, within a production strain, such as Δcos:TriR-P2-c5545 Z1, using a transformation protocol; and to store the strain in glycerol stocks at −80ºC, for later expansion. The frozen strain can thereon be used as a master seed for future culture expansion and further induce it with 0.2% (w/v) rhamnose to produce a new batch of P4-EKORhE transducing particles. In this way, the repressible capability of P4-EKORhE allows for an easier and more reproducible production of P4-based transducing particles.

### R3. High-yield bioproduction of transducing particles with P4-EKORhE

Use of the P4-EKORhE system for transducing particle production, benefits from the controllable feature of its rhamnose-inducible synthetic *ε* cassette. However, optimising the bacterial cell culture conditions upstream, together with downstream processing that preserves particle integrity and maximises yield, remains essential. Figure 6 highlights the improvement obtained when the literature standard PEG8000 phage purification is replaced with an ultrafiltration protocol[43] and switch the culture medium from LB to TB broth.

This controllability of P4-EKORhE enables the implementation of a specialised bioprocess in which continuously enriched phage lysates in a fed-batch fashion can be supplemented with growing cultures in parallel successive batches of fresh Δcos-P2-c5545 Z1 cells harbouring P4-EKORhE (as shown in the steps 5 and 6 of Figure 8). The process shows in Figure 6 yields of at least 10^9^ infectious particles per each drop of 10 *μ*L used for the spot assay (Figure S4), which translates to a lysate titer of approximately 10^12^ pfu/mL (*i.e*. a transduction capacity of 10^12^ cfu/mL), according to the formula below:

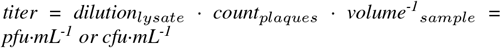

**Fig. 8.**
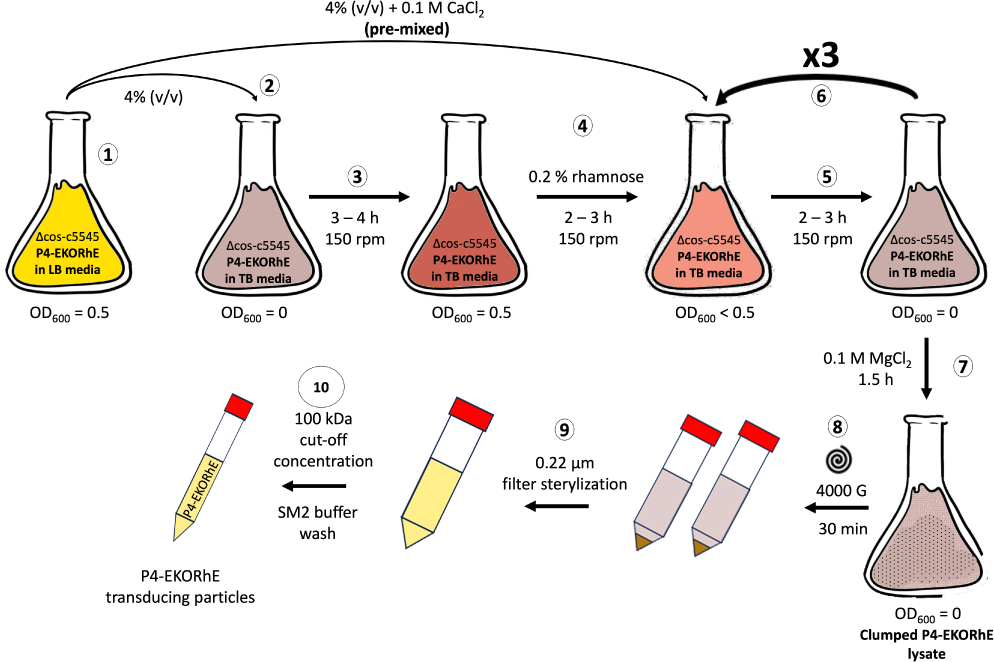
High yield P4-EKORhE based transducing particle production protocol. Following steps from **(1)** to **(10)**: **(1)** A flask containing a Δcos:TriR-P2-c5545 Z1 culture maintaining the P4-EKORhE cosmid is grown in LB (Luria-Bertrani) media un-til reaching exponential phase (*i.e*. OD600 = 0.5). **(2)** A 4% (v/v) of the exponentially growing LB culture is transferred into a flask containing TB (Terrific Broth) media. **(3)** The TB culture is grown to an exponential phase. **(4)** The TB culture is supplemented with 0.2% (w/v) rhamnose for the activation of the P2 prophage to enable the production of P4 particles, and a pre-mixed solution of exponentially growing cells with CaCl2 is prepared to reach a concentration of 0.1 M. **(5)** The TB culture is grown for 2 - 3 hours to accumulate lysed cells containing a proportion transduc-ing particles. **(6)** The TB culture is re-supplemented at a 4% (v/v) with the initial exponentially growing LB culture of Δcos:TriR-P2-c5545 Z1 and re-balanced to maintain 0.1 M CaCl2 and 0.2% (w/v) rhamnose for as many cycles as necessary (*i.e*. from **5** to **6**). **(7)** The TB culture is supplemented with 0.1 M MgCl2 and further grown for 1.5 hours or until clumps are observed. **(8)** The clumped lysate is pelleted at 4000 G for 30 min to separate cell debris. **(9)** The supernatant is filter-sterylyzed. **(10)** The lysate is concentrated using a 100kDa molecular-weight cut-off ultrafiltration column and replaced with SM2 buffer to obtain a cleaned-up final lysate of P4-EKORhE transducing particles.

Therefore, Figure 6 shows how the P4-EKORhE system can be optimised to yield 10^12^ cfu/mL of transducing particles by performing up to four enrichment cycles, utilising a nutrient-rich Terrific Broth (TB) and the ultrafiltration protocol as a physical means of separation and concentration (Figure 8).

### R4. Transduction of the multi-lysins cassette with P4-EKORhE

In Figure 7.A the antimicrobial potential of the multi-lysins cassette delivered by P4-EKORhE particles is demonstrated. The optical densities (OD_600_) of a transduced bacterial culture are measured over 5 hours and compared with natural phages T4 and K1F using the same target bacteria (Figure 7.B and 7.C).

Figure 7 shows that P4-EKORhE-multi-lysins particles exert antimicrobial activity proportional to the MOI (Multiplicity of Infection). Especially effective activity is observed at an MOI of 10. In the positive control, where the tetR repressor is inactivated with 200 nM aTc, the antimicrobial response was distinctive compared to the negative control in which active tetR repressor significantly inhibited lysin expression. The target strain’s response matched or exceeded that of the positive control over the 5-hour measurement period.

To facilitate discussion, Figure 7.C uses the relative area under the curve (rAuC) for the growth curves in Figure 7.A and 7.B to compare antimicrobial effectiveness among samples and MOIs for both P4-EKORhE and T4/K1F-GFP phages. These results show that infecting *E. coli* BW25114 with T4 produces a response similar to that observed with controllable P4-EKORhE.

Of notice, K1F phages specialise in infecting encapsulated strains of *E. coli*[21], so their effectiveness against normal strains such as BW25113 is expected to be compromised, as shown by the figure and corresponding rAuC values. Figure 7.A, 7.B and 7.C confirm the initial characterisation of the multi-lysins cassette in Figure 4. Notably, increasing the MOI from 10 to 100 for natural phages does not produce an appreciably different result, even though an MOI of 10 produces transient growth at about 1 hour post-infection, while P4-EKORhE delays growth until after 2.5 hours as reflected in the rAuC. Moreover, at an MOIs of 100, the antimicrobial activity in negative control strains of *E. coli* Marionette where the tetR repressor remains active in the absence of aTc.

Regarding the effect of MOI on the bacteriostatic activity of P4-EKORhE encoding the multi-lysins cassette, Figure 7.A, and literature [44], indicate that at high MOIs (100), “lysis-from-without” may contribute to the observed results, although the multi-lysins cassette plays the primary role at lower MOIs (10). This effect becomes apparent in the negative control results at MOIs of 100 without aTc. Figure 7.C shows that the differences in rAuC values for MOIs of 10 versus MOIs of 100 with P4-EKORhE do not scale equally between BW25113 and the Z1 controls, suggesting that “lysis-from-without” does not fully account for the differences. However, the non-induced Z1 strains display a notable increase in antimicrobial effectiveness only at an MOI of 100, which may result from “lysis-from-without” if the tetR repressor remains fully functional in the absence of aTc.

An alternative, although less likely, would be the effect of tetR repressor depletion. At high MOIs (*e.g*. 100), a single bacterial cell might be transduced multiple times, exhausting its tetR repressors and leading to leaky gene expression of the multi-lysins cassette. This phenomenon could contribute to the bacteriostatic effect observed at at high MOI but is not apparent at lower MOIs of 10 (compare the rAuC and the bacteriostatic times for the “Z1 + aTc” versus the “Z1 non-aTc”, when using the P4-EKORhE, in the rAuC plot of the Figure 7.C).

Whether high MOIs cause effects via “lysis-from-without” or by tetR repressor depletion remains speculative. Further experiments, for example using P4-EKORhE carrying blank genetic sequences, could clarify this contribution. Based on this data, it is possible to conclude that the substantial differences between aTc-induced and uninduced Z1 cells cannot be solely attributed to the multi-lysins cassette, as antimicrobial effectiveness—albeit milder—at low MOIs (10) is still observed, which is less pronounced than in the “Z1 Marionette + aTc” positive control.

If “lysis-from-without” occurred at lower MOIs, the 2.5-hour lysis delay in the negative control (Figure 7.A) would not be observed. In both replicas (Figure 7.A and Figure S2) for non-activated Z1 cells infected with P4-EKORhE, a mild antimicrobial activity, at an MOI of 100—indicated by a bump in growth at 2.5 hours starting at time = 0—which suggests that the effect in the negative controls at high MOIs likely results from “lysis-from-without” or tetR repressor depletion, is observed.

This data underscore the potential of P4-EKORhE particles as an antimicrobial alternative to natural phages. Figures 7.B and 7.C show BW25113 and Z1 Marionette, when infected with T4 phage, fail to grow during the the 5-hour measurement period—–similar to transduction with P4-EKORhE. These results support the feasibility of using transducing particles as an alternative to natural phages, with the advantage of full control over lysis mechanisms and without relying on viral replication in target bacteria.

### The P4-EKORhE on A549-tissue cultures as a model of infection

Experiments measuring optical cell densities in pure bacterial cultures are used as an initial assessment of transducing particle effectiveness, alongside natural phages. However, to develop a potential antimicrobial therapeutic, it is valuable to replicate experimental conditions similar to real scenarios. Therefore, an infection model is employed that uses A549 human lung epithelial immortalised cells to further assess the capacity of P4-EKORhE encoding the multi-lysins cassette to perform as an antimicrobial in a more complex environment.

In Figure 7.D, the ratio of bacteria to nuclei is demonstrated from images taken in two experiments: one where a 100 *μ*L of an SM2-buffer solution containing `P4-EKORhE` transducing particles harbouring the multi-lysins cassette is supplementated after 1 hour of bacterial incubation on the A549 cell layer, and another where a 100 *μ*L of blank SM2 buffer is added instead. The figure reveals that the differences in bacterial count after one hour of bacterial growth followed by two hours post-treatment with P4-EKORhE indicate the system’s ability to deliver the multi-lysins cassette in a complex environment. Figure 7.D shows a remarkable difference in average bacterial counts between samples treated with P4-EKORhE particles in SM2 buffer (target samples) and those treated with SM2 buffer only (negative control). However, the significance analysis for a sample size (n) of n = 6 yields only p = 0.31, suggesting the need for further optimization or larger sample sizes to substantiate the observed differences. The images in Figure 7.E further illustrate this remarkable difference in averages, and additional images used for bacteria/cell nuclei counts are available in Figure S12 for target samples and Figure S13 for negative controls.

In the context of evaluating the multi-lysins cassette as a gene-delivered antimicrobial, these results demonstrate the potential of P4-EKORhE as a high-yield transducing particle bioproduction system capable of delivering a genetically-encoded antimicrobial (the multi-lysins cassette) in a human cell culture infection model, thereby corroborating the results obtained from pure bacterial cultures.

## Conclusion

### Effectiveness of the multi-lysins cassette

Regarding the effectiveness of using lysins, this work demonstrates that the multi-lysins cassette—encoding a combination of lysins and lysis-accessory proteins (such as holins and spanins)—constructs a genetically encoded antimicrobial with synergistic effects by activating multiple lysis mechanisms. The results obtained suggest that the incorporation of additional kill-switches, such as nuclease elements (*e.g*. CRISPR-Cas9)[45] or alternative growth inhibitors that target divergent bacterial survival pathways, could further enhance the cassette’s potency.

### High-yield bioproduction with P4-EKORhE

The bio-engineered P4-EKORhE, a minimised, controllable, and conditionally propagating phage that employs temperate P2 phages with *knocked-out* cos-sites, produces virus-free P4-like transducing particles at remarkably high yields (ranging from 10^6^ - 10^10^ pfu/mL) while eliminating infectious wild-type P2 or P4 phages. The *ε* gene is used as a transactivator to exploit natural P4 phage proteins that activate temperate P2 phages, thereby enabling reproducible and controllable production of transducing particles. The further downstream processing optimizations—using nutrient-rich TB and an optimised hybrid fed-batch process—achieve yields greater than 10^12^ pfu/mL, demonstrating that the P4-EKORhE system provides an optimal procedure for high-yield production of transducing particles.

### Transduction of the multi-lysins cassette with P4-EKORhE

The transduction of the multi-lysins cassette via P4-EKORhE produces an antimicrobial effect that is not compromised but rather enhanced compared to the results obtained in transformed bacterial cell cultures. The results obtained from pure bacterial cultures closely resemble those found with natural phages such as T4 or K1F-GFP, highlighting the clinical potential of both P4-EKORhE and the multi-lysins cassette.

Moreover, this data indicate that antibiotic marker selection (used to maintain the transduced cosmids within bacterial hosts) does not impair performance at MOIs of 10 or 100, suggesting that antibiotic co-delivery is unnecessary for achieving similar or superior bacteriostatic effects compared to systems where kanamycin (50 *μ*g/mL) is added before aTc activation. An improved antimicrobial effect can be attributed to multiple transducing particles that likely transduce cosmids multiple times during the initial hours of infection. Additionally, at high MOIs (100), the phenomenon of “lysis-from-without” may enhance the effect, as indicated by comparisons between positive and negative control outcomes with P4-EKORhE.

The use of P4-EKORhE harbouring the multi-lysins cassette in A549 cell cultures contaminated with E. coli, show (despite with a p-value of 0.31) a notable reduction in average bacterial counts that correlates with results from pure bacterial cell cultures. The findings invite further experiments at higher MOIs, where a clearer difference between treated and untreated samples may be observed, especially considering that P4-EKORhE does not propagate in target host bacteria.

Ultimately, this data demonstrates that encoding the multi-lysins cassette in a cosmid packaged within a P4-EKORhE system produces a non-replicative, yet conditionally propagable, finely tunable antimicrobial agent as effective as natural phages. Therefore, this work provides a comprehensive proof-of-concept gene therapeutic where a phage-based transducing particle with a genetically encoded antimicrobial offers an alternative to natural phages by using P4-EKORhE as gene-delivery vehicle to produce high yields of transducing particles for delivering the multi-lysins cassette into target bacteria—–both in pure bacterial cultures and in a model of infection using A549 human epithelial cell cultures infected with *E. coli*. This approach opens the door to novel and effective antimicrobial therapeutics targeting a range of pathogenic bacterial strains.

## Materials and methods

### MM1. General Materials

All chemicals were purchased from Sigma Aldrich (now Merck) or ThermoFisher Scientific unless stated otherwise. Primers and gene fragments were commercially synthesised by Integrated DNA Technologies (IDT) or GENEWIZ, US. The stock and working concentrations of antibiotics used in this work are stated in Table S1. All strains used throughout this work are stated in Table S3.

For the construction of genetic devices (Supplementary Section SS4 and SS8), Q5 High-Fidelity Polymerase, Phusion High-Fidelity DNA Polymerase and NEBiuilder HiFi DNA Assembly Master Mix were purchased from New England Biolabs (NEB) and used according to protocol. All enzymes were stored at −20 °C and used with the buffers provided according to the manufacturer’s instructions.

### MM2. Mass-spectrometry and proteomics

In order to ascertain that bacterial cells are eliminated upon activation of the multi-lysins cassette when using anhydrous tetracycline (aTc), it is useful to identify these lysins on a western blot, only when the circuits are activated, in order to relate the correspondence between gene expression and a decrease in the optical cell densities, as shown in Figure 4. However, because the encoded lysins included in the multi-lysins cassette are multiple (*i.e*. up to six different coding sequences, Figure S11), when constructing the system it was unfeasible to include *Histidine-tags*, which would otherwise facilitate its purification and identification, because having repetitive DNA sequences within the cassette would generate genetic stability problems in the cosmids encoding repetitive sequences, which would lead to the impossibility of gene-synthesising it using a commercial supplier.

Therefore, in order to demonstrate the presence of the lysins, upon activation with aTc, and to relate its expression to its antimicrobial effect, an SDS-PAGE gel for expression verification (Protocol SS6.12), and later extraction for Mass-Spectrometry (MS) analysis was employed after the protein extraction of experimental samples (Protocol SS6.11). However, because bacterial cells would be disrupted upon activation of the lysins cassette, an SDS-PAGE gel did not yield bands sharp enough to be distinguished by raw inspection (*i.e*. to distinguish them among the remaining proteins in the lysate). Instead, a lower fraction band of the SDS-PAGE gel was used for protein extraction and MS, confirming the presence of the lysins and lysis-accessory proteins showcased in this work (Figure 4).

To prepare samples for MS (Supplementary Section SS3), induced and non-induced protein lysates from a culture of Marionette Z1 bacterial cells containing a P4-EKORhE cosmid *i.e*., in Figure 5, which harbours the aTc-inducible multi-lysins cassette showcased in this work, was processed and purified using a standard protein extraction protocol (see Protocol SS6.11). Two types of lysates were produced, one coming from the pellet, or insoluble phase (*i.e*. as lysins are expected to act within cell walls and membranes), and another which was considered a “raw lysate” *i.e*., containing both soluble and insoluble phases of the lysate.

Four different samples of pure bacterial cell cultures (*E. coli* Z1 cells) transformed with the multi-lysins cassette cosmid were grown in 1 mL Eppendorf at 37ºC, on shaking conditions, until reaching the exponential growth phase (about 0.4-0.5 OD_600_). Subsequently two of the samples were aTc-induced with 200 nM[46] of anhydrous tetracycline and grown for a further two hours to wait for the lysis of the bacterial cell culture. The bacterial cell cultures were treated with Lysis buffer (50 mM Tris-HCl, 150 mM NaCl, 1% Triton X-100, and 5 mM EDTA) and sonicated to make sure both induced and non-induced samples were treated equally and also so that non-induced samples are surely lysed (see Protocol SS6.11 for more details).

### MM3. Preparation of optical density measurements

In order to ascertain the antimicrobial effectiveness of the P4-EKORhE transducing particles encoding the multi-lysins cassette (Figure 7.A and 7.B), replicas of 96-well plates were utilised. For each well on a 96-well plate (Cellstar Multiwell Plates, Sigma-Aldrich), 100 *μ*L of bacterial culture of *Escherichia coli* at an OD_600_ = 0.1 were dissolved in doubly concentrated LB media, and half diluted with an additional 100 *μ*L of, either an SM2 buffer solution supplemented with P4-EKORhE-multi-lysin transducing particles or T4 or T7 phages (at MOIs of 10 or 100) or with an SM2 buffer only solution for blanks (*i.e*. for MOI = 0). The plates were measured on a plate reader measuring optical densities at about 600 nm to determine bacterial cell densities over time. When the bacterial cells being tested were the Z1 cells, and there was the intention to activate the multi-lysins cassette upon infection with P4-EKORhE, anhydrous tetracycline was added at a concentration of 200 nM[46]. For the transformed multi-lysins cassettes experiments (Figure 4 and Figure 5.D) the same methodology was employed but without doubling

LB media concentration and without adding any SM2 buffer solutions, but kanamycin (50 *μ*g/mL) was used instead to recover the cells after cosmid transformation and selection before activation. Before activation, cosmid-harbouring cells where resuspended in LB media and the multi-lysins cassette was activated using anhydrous tetracycline (aTc, 200 nM)[46] and L-rhamnose was added at a concentration of 0.2% (w/v)[25].

### MM4. Relative area under the curve as antimicrobial effectiveness

For a more accurate comparison between the antimicrobial effect of the P4-EKORhE transducing particles harbouring the multi-lysins cassette with the antimicrobial effect of natural phages T4 and K1F-GFP, the rAuC from the OD_600_ plots over the time span that goes from 0 to 5 hours were extracted for all plots. The relative area under the curve (rAuC) is plotted using the area under the curve (AuC), obtained using the scripted def AuC(x, y): function shown below, of the non-induced, or non-transduced (*i.e*. MOI = 0), sample and using it to subtract the mean values of the induced samples and the standard deviation normalised (by dividing its values) to the blank experiments with non-transduced or non-infected bacterial cells. It is understood that rAuC(MOI = 0) = 0 ± 1 *i.e*. for MOI = 0, the rAuC = 0 with an absolute standard deviation of 1. Therefore, MOIs of zero have not been included in the rAuC plots as these are a relative measure. For integration of each of the curves from the corresponding plots used to obtain the values of AuC, which can later be used to compute the rAuC as aforementioned, the Trapezoid Rule is used to interpolate the area under the curve using the equation encoded in the following script:

~~~
0    def AuC(x, y):
1      sum = 0
2      for i in range(1, len(x)):
3        h = x[i] - x[i-1]
4        sum += h * (y[i-1] + y[i]) / 2
5      return sum
~~~

Where x is time, and y are the point values of OD_600_ over time which form the basis of the total sum of the area under the curve against the x-axis from x=0 (*i.e*. time = 0) until the final value of x.

### MM5. Preparation of animal tissue cultures for transduction

In regards to the mammalian cell culture model of infection, cell lawns of A549 cells (A549 - ATCC CCL-185) were cultivated. The A549 cells were cultured using a base medium F12/K (Gibco/Invitrogen) with 10% fetal bovine serum (FBS) added to the base medium with a standard amount of p/s (*i.e*. penicillin/streptomycin), incubated at 37ºC with 5% CO_2_. The A549 cells were seeded in T25 flasks until transferred to 6-well plates (Cellstar Multiwell Plates, Sigma-Aldrich). When the growing A549 cells were transferred to the 6-well plates, using 2 mL of media per well, these were grown on top of microscopy coverslip slides that have been deposited on each of the 6-well, so as to be able to remove the coverslip slides for later preparation, following an immunohistochemistry protocol, for fluorescence imaging using DAPI, phalloidin and a GFP enhancer to count bacterial cells (*i.e*. transformed with a GFP plasmid) alongside cell nuclei (stained with DAPI) surrounded by the cell cytoskeleton (stained with phalloidin).

Once the 6-well plate with the lawn of A549 cells grew to about 90% confluency the media was exchanged with 2 mL Leibovitz CO_2_-supplemented media (L-15 Medium, Sigma-Aldrich) with 10% FBS only, without antibiotics, before adding bacteria to the media. Following it, 100 *μ*L of bacteria at an OD_600_ of 0.1 was added to the media and grown incubated at 37 ^*°*^C for one hour. After the hour, either a 100 *μ*L of a particular SM2 buffer solution supplemented with P4-EKORhE transducing particles harbouring the *multi-lysins cassette* at an MOI of 10 or a 100 *μ*L of a blank solution of SM2 buffer (*i.e*. for MOI = 0) was added to each well. Further incubation for one hour was effectuated before removing the media and starting the following immunohistochemistry protocol.

For the immunohistochemistry protocol, which is adapted from the literature[21], the liquid phase of the cell cultures grown in the 6-well plate was removed by aspiration. The 6-well plate was placed under the fume hood, and cells were fixed by adding 500 *μ*L of a solution of 4% PFA (paraformaldehyde) before placing the plate on ice for 15 minutes. The liquid was again removed from the plate and its wells were washed 3 times with PBS for 5 minutes each time. The cells were then permeabilised with 500 *μ*L of ice-cold PEM/0.05% saponin (PEM Buffer: 0.1M PIPES/1,4-Piperazinediethanesulfonic acid, Piperazine-1,4-bis(2-ethanesulfonic acid), Piperazine-N,N-bis(2-ethanesulfonic acid), 5 mM EGTA, 2mM Magnesium chloride. Brought to pH 6.8, using a NaOH solution) and later aspirated and washed three times with PBS for 5 minutes. The remains on each well were aspirated before being incubated for 15 minutes, at room temperature, with 500 *μ*L of a 50 mM solution of ammonium chloride. The wells containing the coverslip slides with the cells upon were then aspirated and washed one time, for 5 minutes, with PBS/0.05% saponin. Subsequently, 40 *μ*L of a mixture that contains 300 *μ*L of 0.05% saponin, 7.5 *μ*L of phalloidin and 1.5 *μ*L of a GFP enhancer was added on each of the six wells of the plate and incubated for 45 minutes in dark conditions. The wells were then washed twice, for 5 minutes, with PBS/0.05% saponin and once with PBS. After aspiration, a drip of Fluoroschield (Fluoroshield™ with DAPI, Sigma-Aldrich) was added on each coverslip glass slide on each well of the 6-well plate. The coverslips were then delicately rinsed with water before being put on microscope slides. The microscope slides with the coverslips were then let dry in dark conditions and after sealing with CoverGrip (CoverGrip™, Biotium) sealant these were finally stored at 4ºC until microscopy imaging.

For the microscopy (ZEISS Celldiscoverer 7 — Automated Live Cell Imaging), images were taken at random and those images that could count more than 60 nuclei per image were taken to ensure the homogeneity of the samples from which imaging was performed. Following the obtaining of such images, bacteria were counted in each image respective to the number of nuclei (blue nuclei stained with DAPI) together with bacteria expressing GFP (green dots stained with GFP enhancer).

### MM6. Upstream optimization and downstream processing with P4-EKORhE

For the production of P4-EKORhE-based transducing particles harbouring the multi-lysins cassette, several protocols were tested which have led to different results. The outputs of such protocols are shown in Figure S3.

The proposed protocol was established for producing transducing particles by growing an overnight (O/N) culture of *E. coli* Δcos:TriR-P2-c5545, that was pre-transformed with a P4-EKORhE-multi-lysins cosmid, in a culture media of Luria-Bertrani (LB) with 50 *μ*g/mL kanamycin (to select for the selection marker in the P4-EKORhE) and 10 *μ*g/mL trimethoprim (to select for the Δcos:TriR cassette in the c5545 strain harbouring the P2 prophage) at 37ºC. Then, a 10% (v/v) of the O/N culture was combined with LB media using 50 *μ*g/mL kanamycin and 10 *μ*g/mL trimethoprim. The culture was shaken for 3-4 hours at 150 rpm and 37°C grown until grown to exponential phase (OD_600_ to 0.2 - 0.5). During the exponential phase, the epsilon (*ε*) gene encoded within the P4-EKORhE cosmid was activated by adding 0.2% (w/v) rhamnose to the media and further incubated at 150 rpm and 37°C for 3 hours until lysis is observed. The debris from the lysate was pelleted at 4000 G for 30 minutes. The lysate was then filter-sterilised, with the option to store the filter-sterilised lysates at 4°C if desired.

After the lysate was obtained, a PEG8000 concentration protocol (Protocol SS7.2), or an ultrafiltration protocol[43], was effectuated in order to purify and concentrate the lysate samples with the conditionally propagating transducing particles. When using the PEG8000 protocol, the yields would not improve over 10^6^ transducing particles per millilitre (Figure S3, measured as pfu/mL).

An alternative strategy using ultrafiltration is shown to improve the yields to up to 10^10^ particles per millilitre (as it can be seen in the replica “P4-EKORhE, x1 LB broth + ultrafiltration” shown in Figure S3).

Fortunately, even higher yields of P4-EKORhE transducing particles were obtained when using Terrific Broth (TB) as a culturing media, and using a 4-step enrichment technique, followed by ultrafiltration (as depicted in Figure 8), with observed yields of up to 10^12^ particles per millilitre, as it is seen in the three replicas on the top row of the spot assay shown in Figure 6.

The protocol, using the showcased enrichment technique, is characteristic of P4-EKORhE because its capacity to be repressed and activated (*i.e*. a controllable P4-EKORhE system). This characteristic allows for the multiplication of the Δcos:TriR-P2-c5545 Marionette Z1 *Escherichia coli* lysogenic bacteria in a *fed-batch* fashion while maintaining a “dormant cosmid” before its activation with 0.2% (w/v) rhamnose. A high-concentration of producing bacteria harbouring the P4-EKORhE is thus possible, and the later activation of the P4-EKORhE rends the highest yields obtained throughout this work.

In this procedure, an O/N culture of Δcos:TriR-P2-c5545 E. coli lysogens, previously transformed with the P4-EKORhE cosmid, was grown in Luria-Bertani (LB) medium using 50 *μ*g/mL kanamycin (to select for the selection marker in the P4-EKORhE) and 10 *μ*g/mL trimethoprim (to select for the Δcos:TriR cassette in the c5545 strain harbouring the P2 prophage). Subsequently, a 10% (v/v) of the aforementioned overnight culture was combined with fresh LB media (with kanamycin and trimethoprim) and shaken for 2 hours at 150 rpm and 37°C until the culture was recalled at exponential growth phase (OD_600_ = 0.5).

Concurrently, another 4% (v/v) portion of the overnight culture was pre-mixed with the necessary amount of CaCl_2_ to make a final solution of 0.1 M in a flask containing a volume of Terrific Broth (TB) media, including kanamycin and trimethoprim at the aforementioned concentrations, and shaken for 3-4 hours at 150 rpm and 37°C, reaching the exponential growth phase. Pre-mixture of cells with CaCl_2_ is essential to avoid the precipitation of phosphate salts in the TB broth used for the culture. After reaching the exponential growth phase, 0.2% (w/v) rhamnose was added to the TB culture to induce the production of the *ε* gene, to activate the Δcos:TriR-P2 prophage. After 2 hours of lysis in the TB culture, a 4% (v/v) sample of bacteria from the LB culture was introduced into the lysing TB-grown culture, again pre-mixing it with the necessary amount of CaCl_2_ to maintain a final solution of 0.1 M in the growing TB culture, and compensating the rhamnose to maintain the 0.2% (w/v) proportion. Yet concurrently, a new exponentially growing LB culture was prepared by adding 10% (v/v) of the overnight culture, which was shaken in parallel for a further 2 hours at 150 rpm and 37°C until it reaching the “recall stage” at the exponential growth phase. This growing LB culture was also introduced into the ongoing lysis process of the TB culture, after 2 hours, pre-mixed with the necessary amount of CaCl_2_ to make a final solution of 0.1 M, while maintaining the proportion to a final 0.2% (w/v) rhamnose.

This re-lysing procedure can be repeated as many times as necessary (*e.g*. 0 to 3 times), ideally up to three additional times, to maximise the yield of transducing particles obtained. The lysis procedure was finished by adding 0.1 M MgCl_2_ and the culture shaken at 37 ºC for a further 1.5 hours until visible clumps were observed. Afterwards, the debris from the lysate was pelleted at 4000 G for 30 minutes, followed by filter-sterilization of the lysate with 0.22 *μ*m sterile syringe filters. After filtration, the lysate can be paused and stored at 4°C if desired, until further use. The lysate can then be ultrafiltrated using the Amicon 100 kDa cut-off columns by centrifuging at 3000 G. The recovered, non-eluted, residue contains the highly concentrated P4-EKORhE based transducing particles.

Before titering, the lysate was pre-treated with DNAseI from NEB using 10*μ*L lysate + 10*μ*L DNAseI buffer + 1*μ*L DNAseI + 79 *μ*L of water, incubated for one hour. Because P4 phages are conditionally propagating, lysates are titered using a spot assay (Supplementary Section SS7.4.1), on lawned plates of Δcos:TriR-P2-c5545 Marionette Z1 *E. coli* lysogenic bacteria with 5 mM Ca^++^.

## ACKNOWLEDGEMENTS

R.R.G. is grateful to Gianni Dehò and Federica Briani, from Università degli Studi di Milano, for kindly and promptly providing a comprehensive collection of P4 and P2 phage strains and host cells essential to completing this work. R.R.G. also acknowledges the Monash-Warwick Alliance’s financial sponsorship and support throughout the joint doctoral program. R.R.G. thanks Sahan B.W. Liyanagedera for his help in the laboratory and for scientific discussion, Satya Prakash for his help with difficult cloning, Gurneet Dhanoa and Josh Williams for their help with immunohistochemistry and fluorescence microscopy, and Cleidi Zampronio for her support with Mass-Spectrometry and the analysis of protein sequences. R.R.G. thanks to A.P.S., J.J.B. and his PhD viva jury composed of Paul Jaschke and Danish Malik for their valuable comments on his doctoral thesis that have been directly and indirectly implemented in this manuscript. sAJM.1505 was a gift from Christopher Voigt (Addgene # 108253).

## AUTHOR CONTRIBUTIONS

R.R.G., J.J.B. and A.J. conceived the project, R.R.G. designed, planned and executed experiments. R.R.G. prepared and revised the manuscript. R.R.G. created the bioengineered P4-EKORhE system via gene assembly. R.R.G. designed plasmids and genetic circuits for the creation of P4-EKORhE using Gianni Dehò’s P4vir1, synthetic constructs, and components from available A.J. constructs.

R.R.G. and A.J. constructed a functional prototype of the multi-lysins cassette from a previous prototype by A.J. for a purpose different to the one depicted in this work. R.R.G. modified, characterized the antimicrobial effectiveness of the multi-lysins cassette and repurposed it as a gene-delivered antimicrobial. R.R.G. performed optical density measurements, immunohistochemistry and fluorescence microscopy imaging. All E. coli Z1 host strains belong to A.J. lab. Phage K1F-GFP, host strain EV36, and reagents for immunohistochemistry and microscopy were provided by A.P.S. lab. Manuscript was revised by J.J.B. and A.J.

