## Supplementary material for "High-yield bioproduction of virus-free virus-like P4-EKORhE multi-lysin transducing particles as an antimicrobial gene therapeutic"

#### Supplementary Section SS1: Antibiotics, bacterial strains, and human cell lines

**Table S1.** List of antibiotics used in this work

| Antibiotic | Solvent | Stock (mg/mL) | Working ($\mu$g/mL) |
|---|---|---|---|
| Chloramphenicol (Chlo) | EtOH | 35 | 35 |
| Kanamycin (Kan) | Water | 50 | 50 |
| Trimethoprim (Tri) | DMSO | 10 | 10 |

**Table S2.** List of phage strains used in the work

| Strain | Origin | Purpose |
|---|---|---|
| P1vir1 | A kind gift from Baojun Wang (University of Edinburgh)[47]. | Exclusively virulent variant of P1 phage. This P1 virulent strain is used for P1 transduction in the construction of the *Escherichia coli* $\Delta$cos-c5546-Z1-Marionette strain. |
| P2vir1 | A kind gift of Gianni Dehò and Federica Briani (Universita degli Studio de Milano)[48]. | Exclusively virulent variant of P2 phage. This P2 virulent strain is used in this work to produce stocks of P2vir1 phages or to produce transducing particles contaminated with virulent P2vir1 phage particles. |
| P4vir1 | A kind gift of Gianni Dehò and Federica Briani (Universita degli Studio de Milano)[49] | Virulent P4 phage used in this work as a backbone for the essential region to construct P4-min and P4-EKORhE. |
| K1F-GFP | A kind gift of Josh Williams from Sagona lab (University of Warwick)[21]. | Used as a control natural phage to compare with the bioengineered P4-EKORhE. K1F-GFP is a bacterial virus specialised to infect encapsulated forms of *Escherichia coli* EV36 that has been modified to display a Green Fluorescent Protein (GFP) in its capsid. |

**Table S3.** List of bacterial strains and cell lines used in this work

| Strain | Origin | Purpose |
|---|---|---|
| *Escherichia coli* c5545 | A kind gift from Gianni Dehò and Federica Briani (Universita degli Studio de Milano). | P2 lysogen with *del1* deletion in *old* gene, permitting lambda red expression for Δcos *knock-out* using recombineering[50]. This P2 lysogenic strain used in this work to test transducing particle yields or to produce wild-type P2 phages. |
| *Escherichia coli* Z1 Marionette (sAJM.1505) | sAJM.1505 was a gift from Christopher Voigt (Addgene # 108253)[23] in Jaramillo lab. | Contains different cassettes to enable inducible promoters, of interest the Z1 cassette necessary to repress the lysins when no aTc is present. |
| *Escherichia coli* Δcos-c5545-Z1-Marionette | Constructed using P1 transduction of the Z1 Marionette cassette into Δ*cos c5545 E. coli*[50]. Jaramillo lab. | Production of virus-free P2 and/or P4 phage particles while containing the multi-lysins cassette under the tetR repressor (*i.e.* Z1 cassette) |
| *Escherichia coli* BW25113 | Keio Collection Parental Strain, CGSC[51]. | Proof-of-concept strain used for target host-bacteria validation strain and for the production of virulent P2vir1 phages. |
| *Escherichia coli* BL21(DE3) | Thermofisher (EC0114). | A host for virulent P2vir1 phage production. |
| DH5$\alpha$ Z1 cells | Jaramillo lab. | General cloning procedures while containing the multi-lysins cassette under the tetR repressor (*i.e.* Z1 cassette). |
| A549 human lung epithelial immortalised cells | A kind gift from Vicky Smith from Unnikrishnan lab (University of Warwick). | Used as a human cellular model of infection to test the antimicrobial effectiveness of P4-EKORhE against *E. coli*. |

### Supplementary Section SS2: Replicas

#### SS2.1. Optical densities.

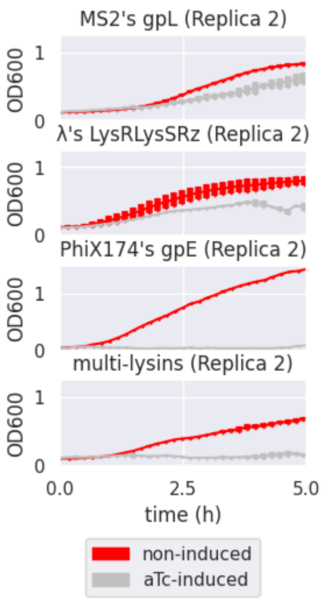


**Fig. S1. Effectiveness of a transformed multi-lysins cassette (replica).** Optical density ($OD_{600}$) measurements for bacterial survival profiles in pure bacterial cell cultures of *Escherichia coli* DH5$\alpha$ Z1 showing the antimicrobial effect of the multi-lysins cassette. The "red lines" represent the bacterial cells harbouring the multi-lysins cassette that have not been induced with anhydrous tetracycline (aTc). The "silver lines" represent bacterial cells harbouring the lysins cassette that have been induced with anhydrous tetracycline at the start of the culture. ** A primary replica is found in Figure 4. All plots use a sample number of n = 3. The thickness of each line represents variability among the "n" samples using the standard deviation.*

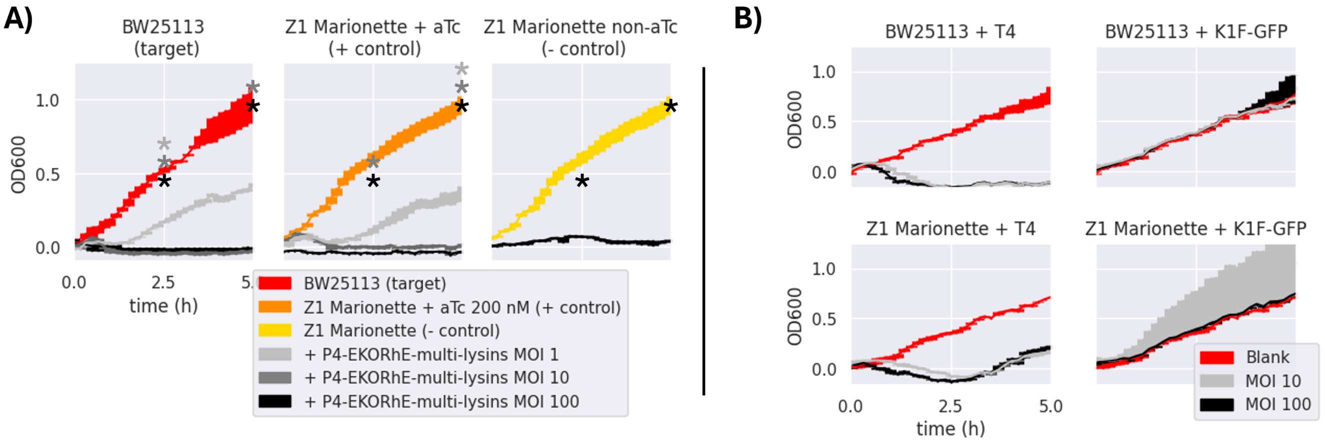


**Fig. S2. A) Antimicrobial activity of P4-EKORhE-multi-lysins (replica).** Optical density measurements of the target and control *Escherichia coli* strains *i.e.*, BW25113 and Z1 Marionette, respectively when exposed to P4-EKORhE-multi-lysins particles at different MOIs. No selection antibiotics were used for the realization of these experiments. * On this replica, for the (- control) the MOI of 10 is missing and done only on the replica shown in Figure 7.A with n=3. ** On this replica, the Z1 Marionete (- control) is the same as the Z1 Marionette + aTc 200 nM (+ control). These exceptions were due to these experiments being a preliminary test that has been used here to further corroborate the data beyond the n=3 samples demonstrated of each replica graph (*i.e.* which becomes n=6 considering both replicas). **B) Antimicrobial activity of enterobacteriophages to its respective hosts (replica).** Antimicrobial activity of enterobacteriophages to its respective target and control E coli strains *i.e.*, BW25113 and Z1 Marionette when exposed to natural enterobacteriophages containing a replicative virus at different MOIs. For the corresponding replica of the same set of experiments see Figure 7.A and Figure 7.B. n=3 for all samples and error bars are indicated by the thickness of each line.

**SS2.2. Spot-assays.** In the spot assay of Figure S3, it is shown the capacity of P4-EKORhE to propagate by means of a spot assay. The spot assay can be used to determine the yield of the lysate sample because the P4 phages are conditionally-propagable in the $\Delta$cos:TriR-P2-c5545 Z1 strain of *Escherichia coli* (see Table S3).

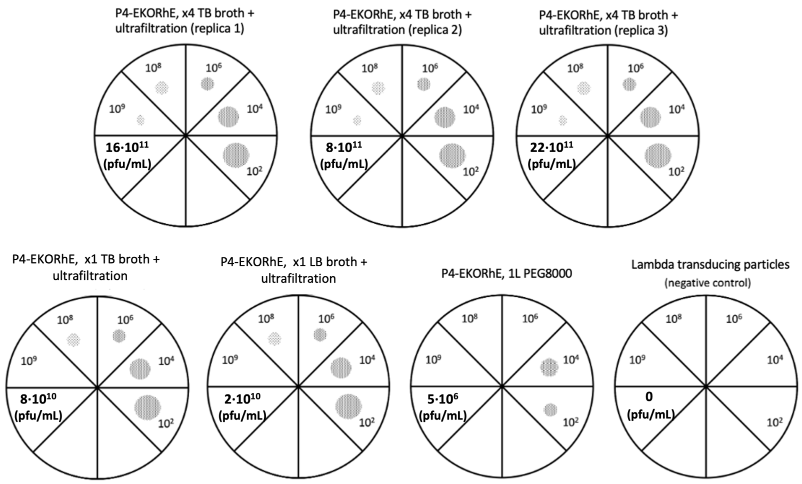


**Fig. S3. Virtual spot assay of P4-EKORhE on *Escherichia coli* $\Delta$cos:TriR-P2-c5545 Z1.** Assessment of the production yields of P4-EKORhE using varied protocols using TB broth, LB broth and/or PEG8000 or ultrafiltration as a concentration protocol, respectively. The values 1x or 4x refer to the number of enrichment cycles used for lysate production (see Figure 8). Virus-free, $\lambda$ phage-based transducing particles ($10^3$ cfu/mL) were used as a negative control due to their incapacity to propagate. The real plates with their corresponding spots are found in Figure S4. The final yields marked in pfu/mL require the consideration that each spot was constituted with 10 $\mu$L lysate.

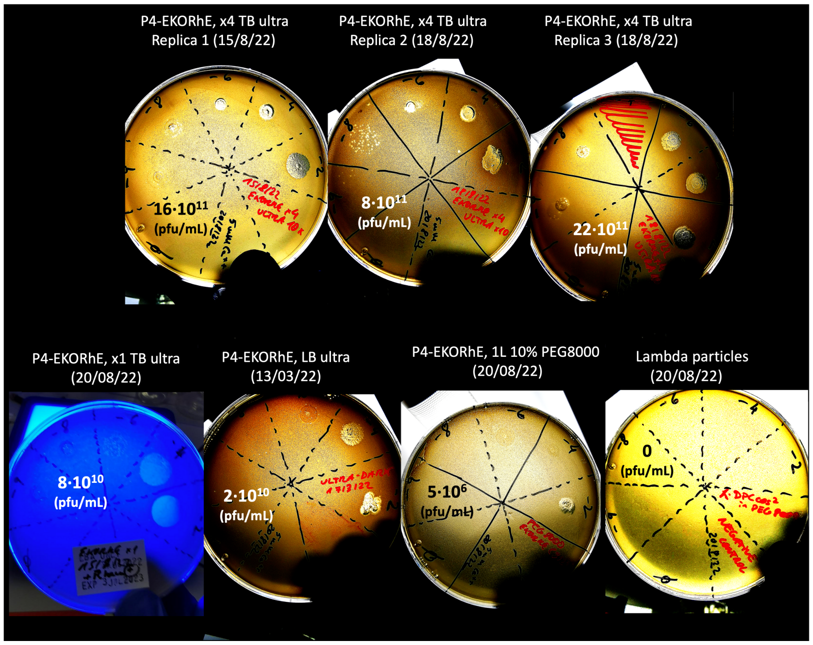


**Fig. S4. Spot assay of P4-EKORhE on E coli $\Delta$cos-5545.** Assessment of the production yields of P4-EKORhE using varied protocols using TB broth, LB broth and/or PEG8000 or ultrafiltration as a concentration protocol, respectively. The virtual plates with their corresponding spots are found in Figure S3. The final yields marked in pfu/mL require the consideration that each spot was constituted with 10 $\mu$L lysate.

#### Supplementary Section SS3: Mass-spectrometry and proteomics

**SS3.1. NanoLC-ESI-MS/MS Analysis.** Reversed-phase chromatography was used to separate tryptic peptides prior to Mass Spectrometric (MS) analysis. Two C18 columns were utilised, an Acclaim PepMap $\mu$-precolumn cartridge 300 $\mu$m i.d. x 5 mm 5 $\mu$m 100 Å (Thermo Fisher Scientific, Waltham, MA, USA) and a 75 $\mu$m x 40 cm 1.9 $\mu$m (Bruker nanoElute Forty Analytical column). The columns were installed on an Ultimate 3000 RSLCnano system (Thermo Fisher Scientific, Waltham, MA, USA). Mobile phase buffer A was composed of 0.1% formic acid in water and mobile phase B 0.1% formic acid in acetonitrile. Samples were loaded onto the $\mu$-precolumn equilibrated in 2% aqueous acetonitrile containing 0.1% Trifluoroacetic and peptides were eluted onto the analytical column at 350 nL/min by increasing the mobile phase B concentration from 4% phase B to 25% over 36 min, then to 35% phase B over 10 minutes, and to 90% phase B over 3 minutes, followed by a 10-minute re-equilibration at 4% phase B. Ultimate 3000 RSLCnano was coupled online to a hybrid timsTOF Pro (Bruker Daltonics, Germany) via a CaptiveSpray nano-electrospray ion source[52]. The timsTOF Pro was operated in Data-Dependent Parallel Accumulation-Serial Fragmentation (PASEF) mode. Peptides were separated by ion mobility depending on their collisional cross sections and charge states. The method settings were as follows: mass range 100 to 1700 m/z, ion mobility range 1/K0 Start 0.6 Vs/cm$^2$ End 1.6 Vs/cm$^2$, Ramp rate 9.42 Hz and Duty cycle 100%.

**SS3.2. Mass-Spectrometry data analysis.** The raw data from Mass Spectrometry (MS) was searched using label-free quantitation by FragPipe version 18.0 (https://fragpipe.nesvilab.org/) against the *Escherichia coli* database (https://www.uniprot.org/proteomes), the coding sequences of the proteins included in the multi-lysins cassette (*i.e.* MS2's gpL, PhiX174's gpE, Lambda's LysR, Lambda's LysS and Lambda's Rz), and the common contaminant database. The gene product of *aphA1*, the Aminoglycoside 3'-phosphotransferase (UniProt P00551[53]), which is the kanamycin-resistant selection marker used to construct and select all the lysis- and lysis-accessory- encoding plasmids used throughout this work, including the P4-EKORhE, was used as a positive control. For the database search, peptides were generated from a tryptic digestion with up to two missed cleavages, and carbamidomethylation of cysteines as fixed modifications. Oxidation of methionine and acetylation of the protein N-terminus were added as variable modifications. MS/MS data was filtered using a false-discover rate of 0.01. Scaffold software version 5 (https://www.proteomesoftware.com/products/scaffold-5) was used to analyse the results. The coding aminoacid sequences of the proteins, included in the multi-lysins cassette (*i.e.* MS2's gpL, PhiX174's gpE, Lambda's LysR, Lambda's LysS and Lambda's Rz), used to confirm the analysis are compiled in the Supplementary Section SS8 for Genetic and aminoacid sequences.

**SS3.3. Normalised Total Spectra.** The Normalised Total Spectra (NTS) was obtained from the presence of peptides belonging to the query sequences shown above (*i.e.* the lysins and lysis-accessory proteins studied in this work) with a 95% probability match, using the Scaffold software version 5 (https://www.proteomesoftware.com/products/scaffold-5)[54] with the data obtained from the aforementioned samples blasted on a Mass Spectrometer. To further indagate into the process of normalization that Scaffold software version 5 software uses to report its results, find online an explanation of "Spectrum count normalization in scaffold"[55]. For the obtaining of the values shown in Results R1, for the NTS of the query sequences identified, MSFRagger search was performed using Label Free Quantitation (LFQ)- The software uses the peak area of all tryptic peptides from a specific protein to calculate the ratio of this protein in different samples. The Scaffold version 5 software uses these quantitative ratio values to display the NTS results using automated output from MS data as their input.

**sp|000004|MS2-gpL (100%), 8,870.6 Da**
**MS2-gpL protein**
**4 exclusive unique peptides, 5 exclusive unique spectra, 10 total spectra, 30/75 amino acids (40% coverage)**

M E T R F P Q Q S Q   Q T P A S T N R R R   P F K H E D Y P C R   R Q Q R S S T L Y V   L I F L A I F L S K   F T N Q L L L S L L
E A V I R T V T T L   Q Q L L T

**sp|000005|PhiX174-gpE (100%), 10,602.7 Da**
**PhiX174-gpE protein**
**1 exclusive unique peptides, 1 exclusive unique spectra, 1 total spectra, 9/91 amino acids (10% coverage)**

M V R W T L W D T L   A F L L L L S L L L   P S L L I M F I P S   T F K R P V S S W K   A L N L R K T L L M   A S S V R L K P L N
C S R L P C V Y A Q   E T L T F L L T Q K   K T C V K N Y V R K   E

**sp|000002|Lambda-lysR (95%), 17,825.9 Da**
**Lambda-lysR protein**
**1 exclusive unique peptides, 1 exclusive unique spectra, 1 total spectra, 13/158 amino acids (8% coverage)**

M V E I N N Q R K A   F L D M L A W S E G   T D N G R Q K T R N   H G Y D V I V G G E   L F T D Y S D H P R   K L V T L N P K L K
S T G A G R Y Q L L   S R W W D A Y R K Q   L G L K D F S P K S   Q D A V A L Q Q I K   E R G A L P M I D R   G D I R Q A I D R C
S N I W A S L P G A   G Y G Q F E H K A D   S L I A K F K E A G   G T V R E I D V

**sp|000003|Lambda-lysS (100%), 11,521.6 Da**
**Lambda-lysS protein**
**2 exclusive unique peptides, 2 exclusive unique spectra, 2 total spectra, 17/107 amino acids (16% coverage)**

M K M P E K H D L L   A A I L A A K E Q G   I G A I L A F A M A   Y L R G R Y N G G A   F T K T V I D A T M   C A I I A W F I R D
L L D F A G L S S N   L A Y I T S V F I G   Y I G T D S I G S L   I K R F A A K K A G   V E D G R N Q

**sp|000001|Lambda-Rz (100%), 17,230.1 Da**
**Lambda-Rz protein**
**10 exclusive unique peptides, 19 exclusive unique spectra, 45 total spectra, 77/153 amino acids (50% coverage)**

M S R V T A I I S A   L V I C I I V C L S   W A V N H Y R D N A   I T Y K A Q R D K N   A R E L K L A N A A   I T D M Q M R Q R D
V A A L D A K Y T K   E L A D A K A E N D   A L R D D V A A G R   R R L H I K A V C Q   S V R E A T T A S G   V D N A A S P R L A
D T A E R D Y F T L   R E R L I T M Q K Q   L E G T Q K Y I N E   Q C R

**sp|000006|aphA1 (100%), 30,980.1 Da**
**(same as APH(3')-I family aminoglycoside O-phosphotransferase [Escherichia coli])**
**5 exclusive unique peptides, 6 exclusive unique spectra, 10 total spectra, 66/271 amino acids (24% coverage)**

M S H I Q R E T S C   S R P R L N S N M D   A D L Y G Y K W A R   D N V G Q S G A T I   Y R L Y G K P D A P   E L F L K H G K G S
V A N D V T D E M V   R L N W L T E F M P   L P T I K H F I R T   P D D A W L L T T A   I P G K T A F Q V L   E E Y P D S G E N I
V D A L A V F L R R   L H S I P V C N C P   F N S D R V F R L A   Q A Q S R M N N G L   V D A S D F D D E R   N G W P V E Q V W K
E M H K L L P F S P   D S V V T H G D F S   L D N L I F D E G K   L I G C I D V G R V   G I A D R Y Q D L A   I L W N C L G E F S
P S L Q K R L F Q K   Y G I D N P D M N K   L Q F H L M L D E F   F

**sp|P0A7W1|RS5_ECOLI (100%), 17,603.0 Da**
**30S ribosomal protein S5 OS=Escherichia coli (strain K12) OX=83333 GN=rpsE PE=1 SV=2**
**32 exclusive unique peptides, 83 exclusive unique spectra, 645 total spectra, 164/167 amino acids (98% coverage)**

M A H I E K Q A G E   L Q E K L I A V N R   V S K T V K G G R I   F S F T A L T V V G   D G N G R V G F G Y   G K A R E V P A A I
Q K A M E K A R R N   M I N V A L N N G T   L Q H P V K G V H T   G S R V F M Q P A S   E G T G I I A G G A   M R A V L E V A G V
H N V L A K A Y G S   T N P I N V V R A T   I D G L E N M N S P   E M V A A K R G K S   V E E I L G K

**sp|P62399|RL5_ECOLI (100%), 20,302.7 Da**
**50S ribosomal protein L5 OS=Escherichia coli (strain K12) OX=83333 GN=rplE PE=1 SV=2**
**44 exclusive unique peptides, 107 exclusive unique spectra, 495 total spectra, 174/179 amino acids (97% coverage)**

M A K L H D Y Y K D   E V V K K L M T E F   N Y N S V M Q V P R   V E K I T L N M G V   G E A I A D K K L L   D N A A A D L A A I
S G Q K P L I T K A   R K S V A G F K I R   Q G Y P I G C K V T   L R G E R M W E F F   E R L I T I A V P R   I R D F R G L S A K
S F D G R G N Y S M   G V R E Q I I F P E   I D Y D K V D R V R   G L D I T I T T T A   K S D E E G R A L L   A A F D F P F R K

**Fig. S5. Peptide coverage of the proteins found with Mass-Spectrometry analysis of bacteria activated with the multi-lysins cassette.** The peptide coverage includes data that uses the number of peptides found with over 95% probability for each of the lysins studied MS2's gpL, PhiX174's gpE, and $\lambda$'s LysR, *lysS* and *Rz*, when inducing the multi-lysins cassette. The gene product of *aphA1* is used as a positive control because the cosmid expresses aphA1 as a kanamycin resistance selection marker. The 30S and 50S ribosomal proteins are also included as controls due to their necessary presence in the bacterial host (*Escherichia coli* Marionette Z1) used for the analysis of gene expression. **aTc-induced:** The protein extract comes from bacteria for which cosmids containing the multi-lysins cassette have been activated by adding aTc (anhydrous tetracycline) to the culture media. **Non-induced:** The protein extract comes from bacteria for which cosmids containing the multi-lysins cassette have not been activated. **Insoluble fraction:** Protein fraction extracted from the pellet of the lysed cells only upon induction with the inducer anhydrous tetracycline (*i.e.* aTc). **Raw fraction:** Protein fraction extracted from samples containing both, the supernatant and the pellet, from lysed cells upon induction with the inducer.

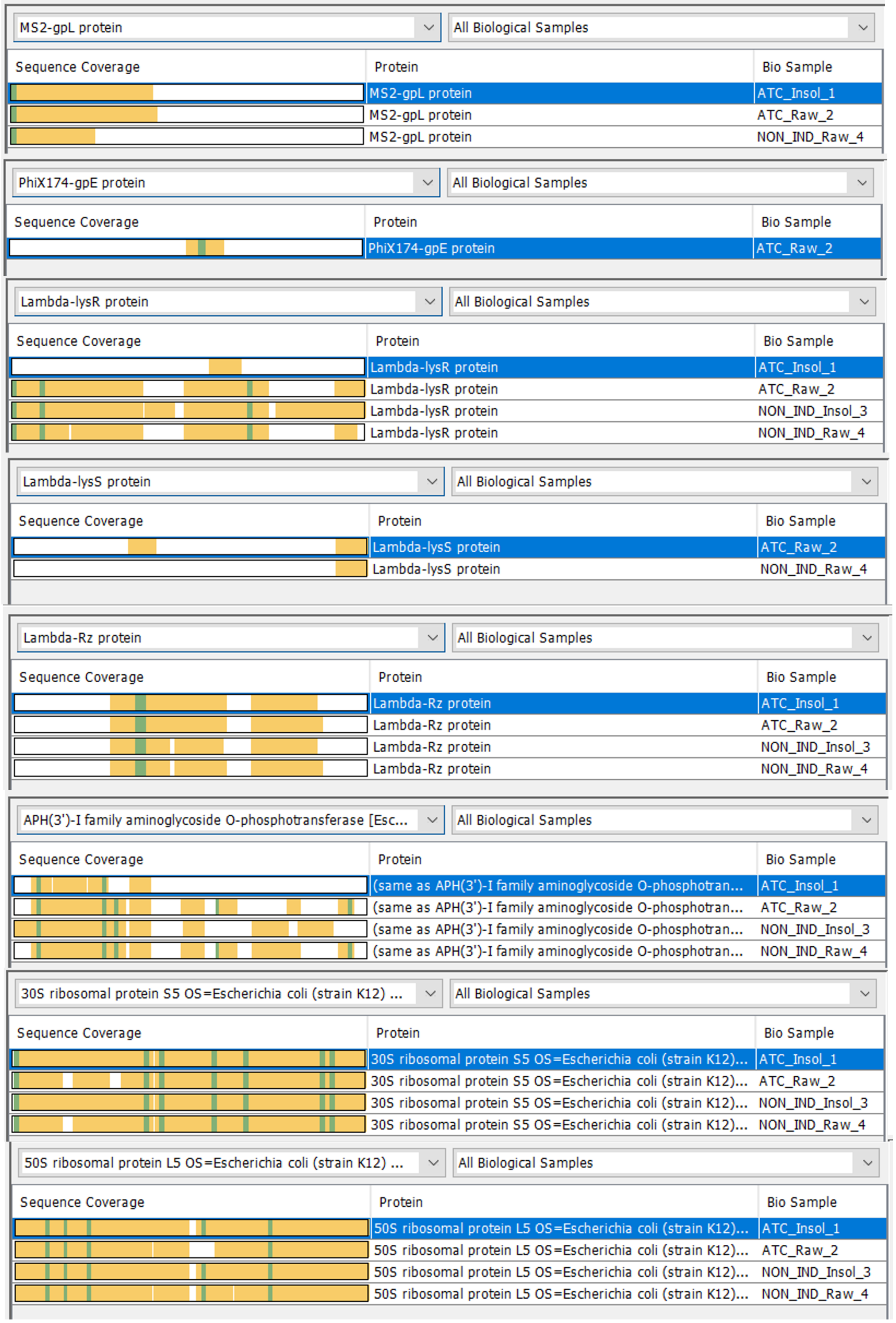


**Fig. S6. Peptide coverage obtained from a Mass-Spectrometry analysis of 4 different samples, two induced and two non-induced: one corresponding to the raw-phase and the other to the insoluble-phase only).** The peptide coverage includes data that uses the number of peptides found with over 95% probability for each of the lysins studied MS2's gpL, PhiX174's gpE, and $\lambda$'s LysR, *lysS* and *Rz*, when inducing the multi-lysins cassette. No peptides were found for LysR and LysS. The gene product of *aphA1* is used as a positive control because the cosmid expresses aphA1 as a kanamycin resistance selection marker. The 30S and 50S ribosomal proteins are also included as controls due to their necessary presence in the bacterial host (*Escherichia coli* Marionette Z1) used to for the analysis of gene expression. **aTc-induced:** The protein extract comes from bacteria for which cosmids containing the multi-lysins cassette have been activated by adding aTc (anhydrous tetracycline) to the culture media. **Non-induced:** The protein extract comes from bacteria for which cosmids containing the multi-lysins cassette have not been activated. **Insoluble fraction:** Protein fraction extracted from the pellet of the lysed cells only upon induction with the inducer anhydrous tetracycline (*i.e.* aTc). **Raw fraction:** Protein fraction extracted from samples containing both, the supernatant and the pellet, from lysed cells upon induction with the inducer.

#### Supplementary Section SS4: Genetic maps and construction of genetic devices

##### SS4.1. Cryptic promoters.

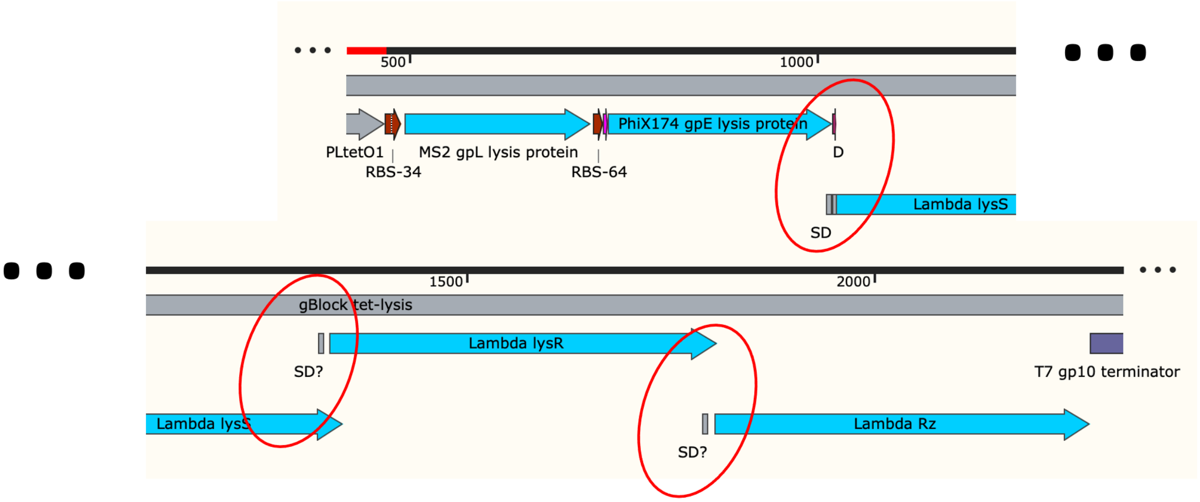


**Fig. S7. Scheme sequence for which *cryptic promoters* can be found in the multi-lysins cassette.** The contiguous nature in which the lysins are arranged gives the possibility to the presence of *cryptic promoters* that may promote the "leakage" or "constitutive expression" of the later genes displayed in the contiguous polycistronic arrangement of genes *e.g.* the LysS, LysR and the Rz proteins *i.e.*, see "SD?" notations through the sequence.

##### SS4.2. Rhamnose-inducible epsilon ($\varepsilon$) cassette.

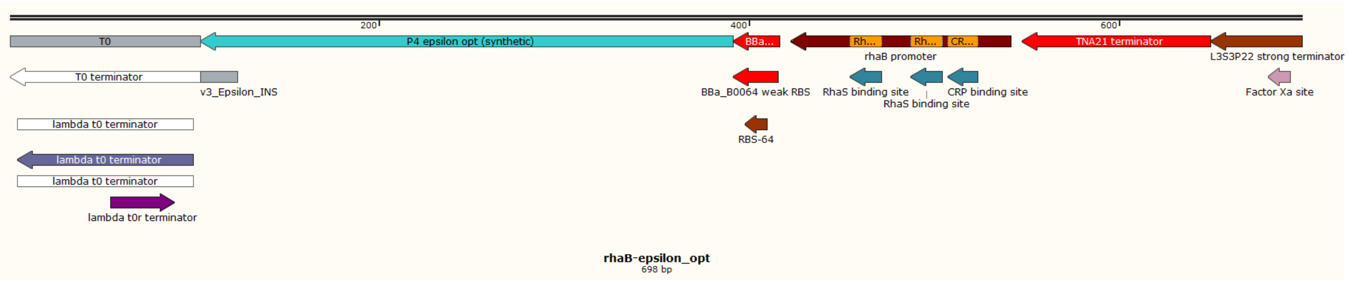


**Fig. S8. A rhamnose-inducible, degenerate, codon-optimised epsilon ($\varepsilon$) cassette.** This cassette is codon-optimised in a way that prevents its recombination with the natural $\varepsilon$ from P4 phage, while still being degenerate to its original sequence and therefore enabling its natural full activity.

**SS4.3. Construction of P4-EKORhE.** The complete version of the P4-EKORhE harbouring the multi-lysins cassette is described in Results, R2, and its fully featured genetic map is shown in Figure 5.A and 5.B. Its antimicrobial effect is demonstrated in Figure 7. The following procedure below which demonstrates the means for its construction.

For the construction of P4-EKORhE, it is necessary to construct an intermediary cosmid, the P4-min. Figure S9, below, schematizes the two-step assembly of P4-min *i.e.*, the essential region of P4 phage natural genome[11], (see featured map of P4-EKORhE in Figure 5) and the further incorporation of the multi-lysins cassette, which is described as follows.

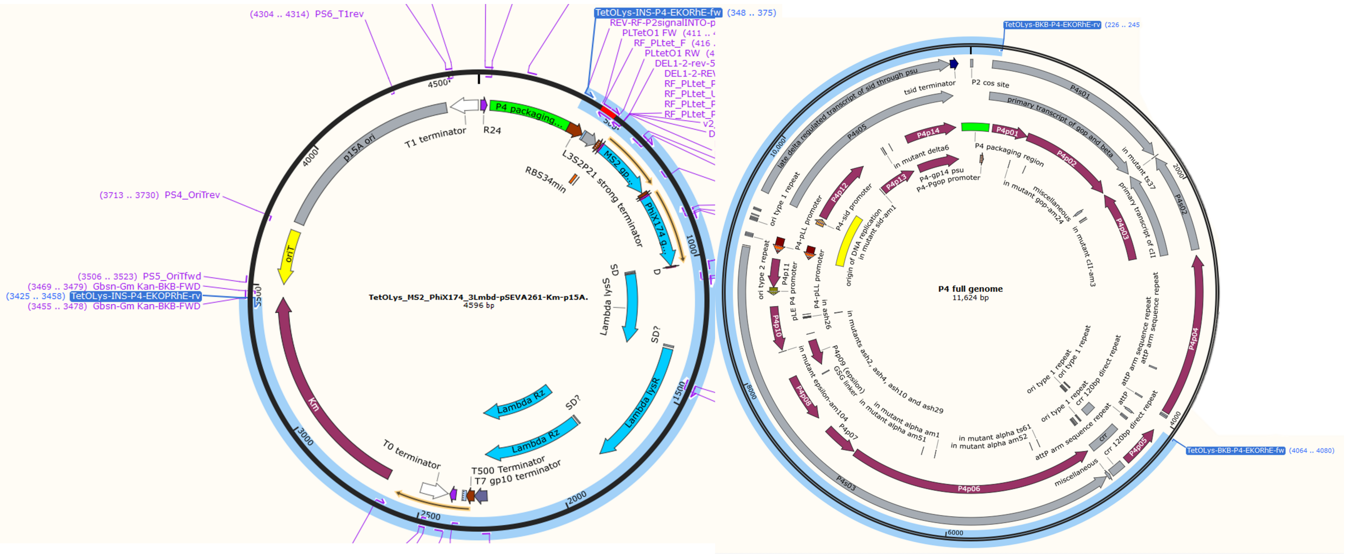


**Fig. S9. Two-step assembly of P4-min with the multi-lysins cassette.** The P4-min is generated by a large deletion on the non-essential region on the P4 full genome (genetic circuit at the right side of the image) that is used to assemble the multi-lysins cassette (genetic circuit at the left side of the image), using the corresponding set of primers.

Table S4 shows the primers used to which this section makes reference, for both the extraction of P4-min out of the P4 phage full genome; and for the 3-step assembly of P4-EKORhE (Figure S10). Subsequently comes the incorporation of the multi-lysins cassette from the p15A cosmid shown at the beginning of this section which covers from the terminators flanking upstream the *tet* promoter (aTc-inducible), til the kanamycin resistance gene flanking downstream the multi-lysins cassette, see images on S11. Once the P4-min cosmid, containing the multi-lysins cassette and the kanamycin-resistance marker is assembled, afterwards the natural epsilon ($\varepsilon$) gene product is *knocked-out* from the P4-min region. For the $\varepsilon$ (also called "P4p09") gene coding sequence *knock-out*, the backbone belonging to the full-natural P4 phage genome is modified using the following primers for selective *knock-out* and finally assembled together with the remaining fragments of P4-min described below, using a general gene assembly protocol (Protocol SS6.6), as depicted further below in Figure S10. The priming sequences responsible for the $\varepsilon$ *knock-out* are shown as "EpsilonKOfwd55" and "EpsilonKOrev55" in Table S4. Simultaneous to the natural $\varepsilon$ *knock-out* generation, proceeds the insertion of a degenerate codon-optimised rhamnose-inducible $\varepsilon$ (as aforementioned, see it in Figure S8). The map of the "rhamnose-inducible, degenerate, codon-optimised $\varepsilon$ cassette" is shown in Figure S8. The sequence of the "rhamnose-inducible, degenerate, codon-optimised epsilon($\varepsilon$) cassette", which was used for assembly to form the final version of the P4-EKORhE, is found in the Supplementary Section SS8.3. The primers used for the partition of the P4-min genome into "assemblable" parts, in combination with the primers shown before for the $\varepsilon$ *knock-out* are shown in Table S4 as "RhamE-EKORhE-P4-BKB-fw", "RhamE-EKORhE-P4-BKB-rv", "RhamE-EKORhE-P4-INS-fw" and "RhamE-EKORhE-P4-INS-rv"

A final three-step assembly (see Figure S10 between the "rhaB-epsilon_opt" gene fragment, and the two remaining fragments within the "P4-min with the multi-lysins cassette", using a combination of all the aforementioned primer sequences, yields the final version of the bioengineered, virus-free, virus-like conditionally-propagating, P4-EKORhE. For the full-featured genetic map refer to Figure 5. For the genetic sequence, refer to the in the Supplementary Sections SS8.5 and SS8.6.

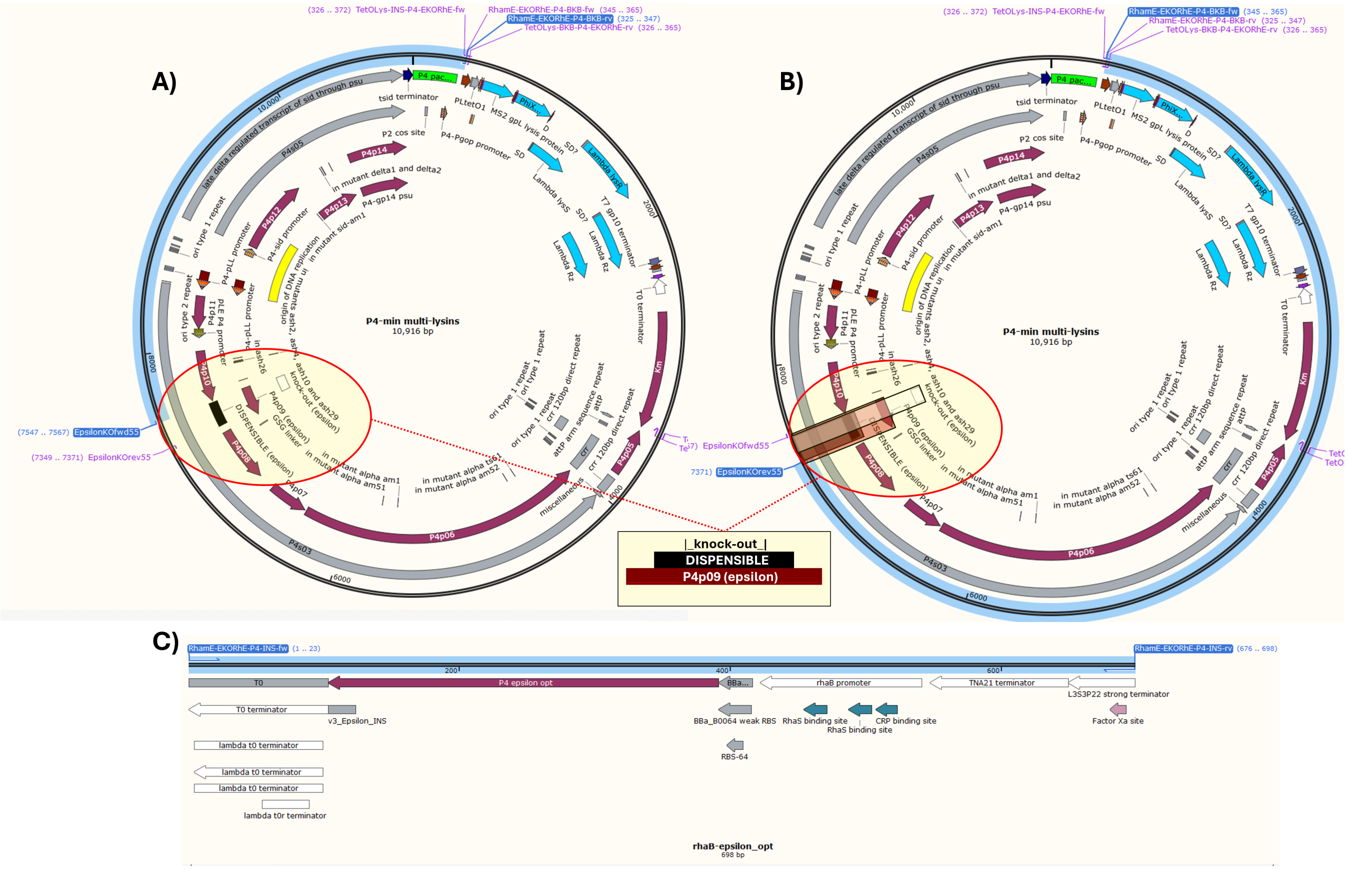


**Fig. S10. Three-step assembly of P4-EKORhE with the multi-lysins cassette.** The P4-EKORhE is generated by combining the highlighted three fragments, using the corresponding set of primers. **A)** The part covering the upstream "dispensible" region, within which the *knock-out* is performed, of only part of the "dispensible region" of the epsilon ($\varepsilon$) coding sequence until downstream the end of the P4 packaging signal. **B)** The part covering the downstream part of the $\varepsilon$ *knock-out* until upstream the tet-promoter of the multi-lysins cassette which is downstream the P4 packaging signal (cos-site). **C)** The part covering the "rhaB-epsilon_opt" gene fragment.

**SS4.4. Construction of the multi-lysins cassette.** For the construction of the multi-lysins cassette, a combination of lysins and lysis-accessory coding sequences put contiguously under the expression of an anhydrous tetracycline (aTc)-inducible promoter (see Figure 3.A), has been constructed in order to find potential synergies in the potency of intracellularly-mediated lysis in target host bacteria that could be improving the effectiveness of genetically-encoded antimicrobial as an alternative to standard antimicrobial molecules or extracellularly delivered lysis proteins. For this purpose, a selection of lysins (gpL, gpE and LysR) and lysis-accessory proteins (*lysS* and *Rz*) have been put together as a means to construct a more effective antimicrobial gene-delivered to target bacterial cells by means of transduction.

The gpE single-gene lysis protein (UniProt P03639[36]) from the PhiX174 phage acts by inhibiting a peptidoglycan biosynthesis enzyme encoded in *mraY*, undermining a critical process for the constitution of the bacterial cell-wall, thus leading to cell wall failure at septation[37]. The **MS2's gpL autolysin** is a 75-residue helical protein used by the MS2 phage (*Emesvirus zinderi, Escherichia phage MS2*), a ssRNA virus which infects gram-negative *Escherichia coli*, as single-gene lysis (Sgl) protein[56][57].

The gpL protein (UniProt P03609[35]) has the particularity of being a cistronic protein which is encoded in the +1 frame of the major coat protein of phage MS2, despite its activity is completely independent as a Sgl[56][58]. In this work, the **$\lambda$'s LysR endolysin**, and the lysis-accessory gene products, the **LysS holin/antiholin** and the **Rz/Rz1 spanins**, respectively, are encoded within phage $\lambda$'s lysis cassette and included downstream to the MS2's gpL and PhiX174's gpE proteins to constitute the *multi-lysins cassette*.

The first protein of the $\lambda$'s cassette, the LysR (UniProt C6ZCX1[59]), is an *endolysin* with transglycosylase activity with bacteriolytic activity that acts by degrading the peptidoglycan in the cell wall of the target host bacteria and coordinates with the holin (*lysS* gene product) and spanin (*Rz* gene product) proteins to concatenate the programmed target host bacteria lysis to facilitate the release of viral particles at the end of the viral cycle. The LysS protein (UniProt P03705[32]) is rather two gene products encoded in different frames of the same *lysS* gene. The *lysS* gene encodes two isoforms: a *holin*, and an *antiholin* (which counteracts the activity of the holin as a regulation mechanism), through different translation start sites which differ by the fact that the holin has 3 transmembrane regions whereas the antiholin lacks the first transmembrane region[33]. Instead, the gene products of *lysS* remain harmlessly accumulated within the cytoplasmic membrane until they attain a crucial threshold concentration, which then initiates the creation of micron-scale pores (holes). In regards to the the *Rz* gene products, the Rz (UniProt P00726[39]) and the Rz1 (UniProt Q37935[40]) spanin proteins are of particular interest because these represent a unique example of two genes located in different reading frames in the same nucleotide sequence and which are necessary for the gene to perform its function (*i.e.* one spanin would not be able to act without the other). While Rz corresponds to the i-spanin (acting on the inner membrane as an integral cytoplasmic membrane protein), Rz1 corresponds to the o-spanin which acts on the outer membrane as an outer membrane lipoprotein, forming what is termed in the literature as the "spanin complex"[41].

Below the fully featured map of a basic cosmid harbouring the multi-lysins cassette (Figure S11):

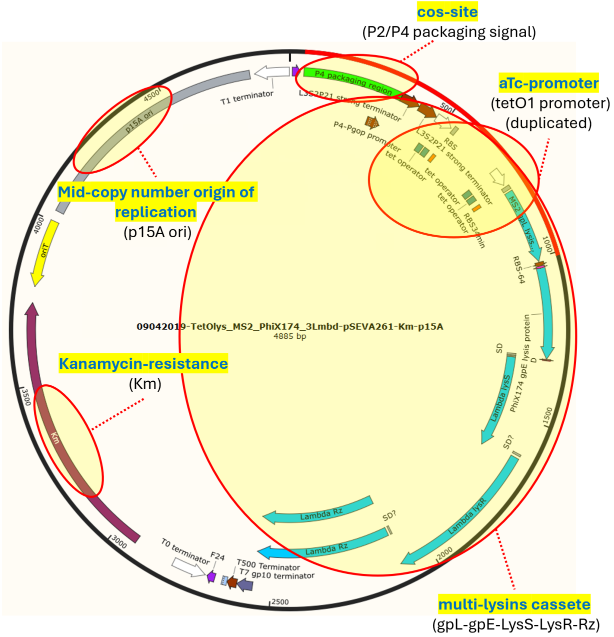


**Fig. S11. The multi-lysins cassette on a backbone containing a packageable "cos" site.** The "TetOlys_MS2_PhiX174_3Lmbd-pSEVA261-Km-p15A" is a cosmid characterized by having a "cos" site (compatible for packaging within P2 or P4 phage-based transducing particles), the multi-lysins cassette (containing MS2's gpL, PhiX174's gpE and $\lambda$'s LysS, LysR and Rz) controlled by the tetO1 promoter (*i.e.* induced with the addition of anhydrous tetracycline/aTc), a kanamycin selection marker, and a p15A origin of replication. The "SD" and "SD?" labels contained within the construct are predicted "Shine-Delgarno" sequences that correspond to the multi-cistronic nature of the $\lambda$ phage lysis operon (*lysS*, *lysR* and *Rz*) and the likely presence of unidentified *cryptic promoters*. The construction of this cosmid (not detailed in this manuscript) shows that during the incorporation of a functional tetO1 promoter, a duplication of the promoter has occurred which does not affect functionality (see Figure 4 for characterization) For a sequencing confirmation, see sequencing tag "90DJ25_61971875_61971875.ab1" in the Supplementary Section SS8.4.

#### Supplementary Section SS5: Immunohistochemistry imaging

+ SM2 buffer

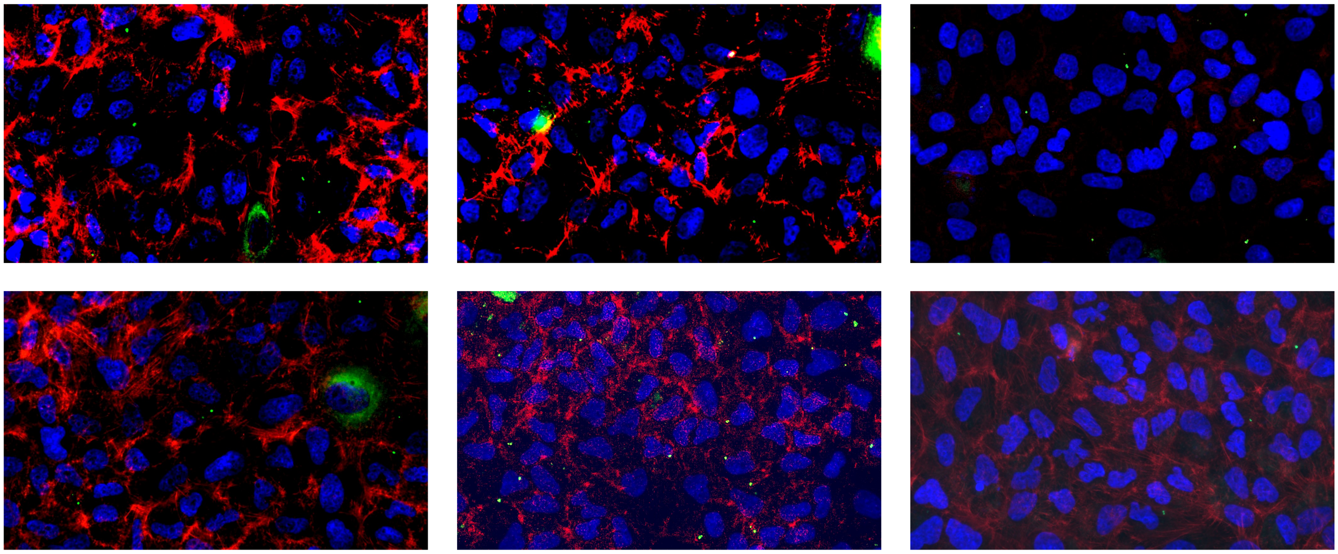

**Fig. S12. Immunohistochemistry imaging using DAPI, Phalloidin and GFP enhancers when no transducing particles were added (SM2 only)** Images n=6 used to quantify the immunohistochemistry imaging. Revert back to Figure 7.E in Results, R4.

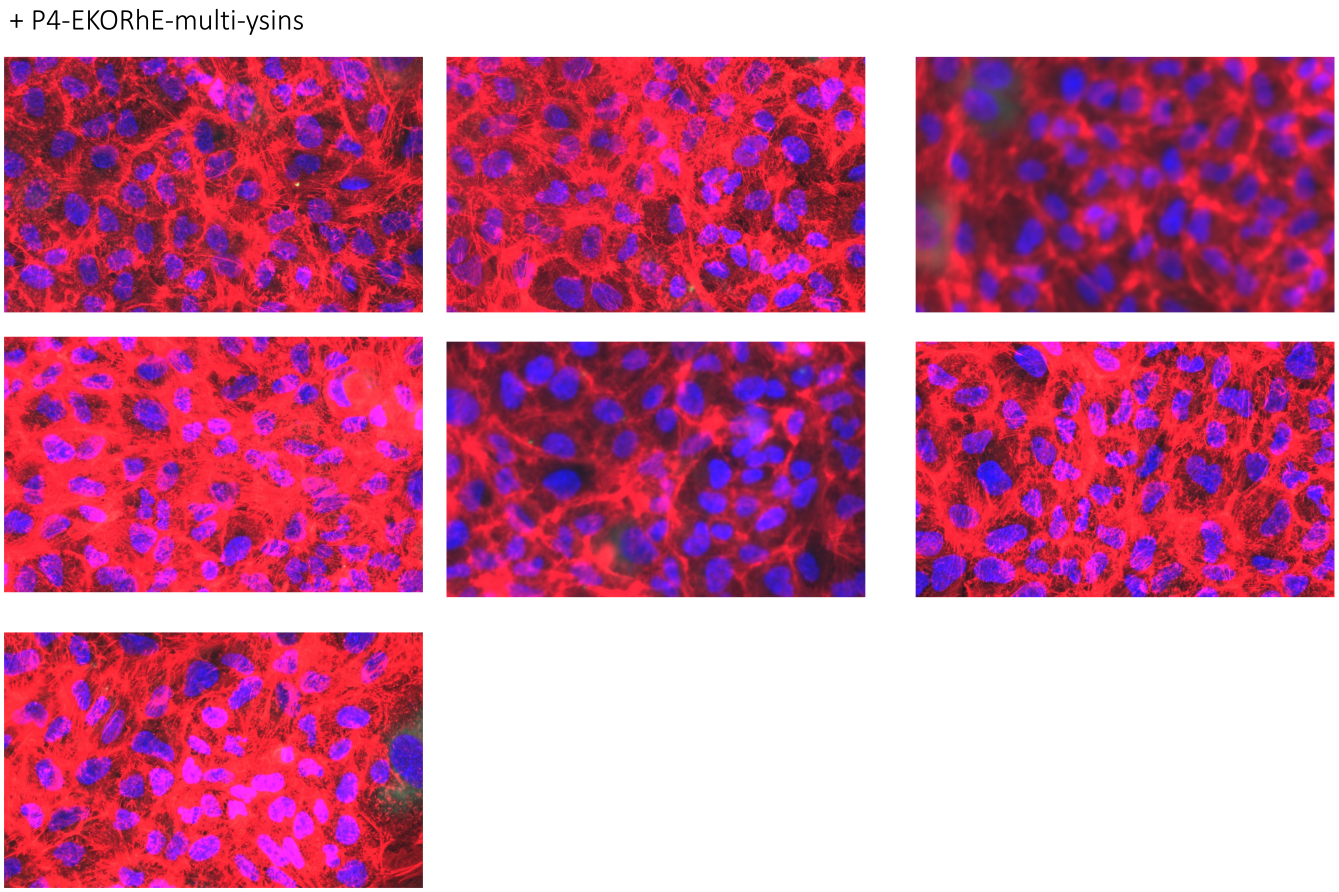


**Fig. S13. Immunohistochemistry imaging using DAPI, Phalloidin and GFP enhancers when P4-EKORhE-multi-lysins transducing particles were added (P4-EKORhE in SM2 buffer)** Images n=7 used to quantify the immunohistochemistry imaging. Revert back to Figure 7.E in Results, R4.

#### Supplementary Section SS6: General standard methods

This section briefly describes the general methods used in this study. Further details are found in the manufacturer's instructions if required.

**SS6.1. Statistical significance.** A measure of statistical significance is given using a standard *t-test* using the values of average and standard deviation and including the number of samples "n" used to calculate the latter. The process has been automated using the SciPy libraries "scipy.stats.ttest_ind_from_stats" for a *t-test* using two independent samples to calculate the likelihood that the null hypothesis is valid using the obtained *p-value*. Throughout this work, whenever the obtained *p-value* is lower than 0.05 the null hypothesis is disregarded and the conclusion that there is a significant difference between the two independent samples is accepted with a relative confidence level of 95%. To see the *t-test* code used for statistical analysis using the SciPy libraries refer to the following link: https://docs.scipy.org/doc/scipy/reference/generated/scipy.stats.ttest_ind_from_stats.html .

**SS6.2. Plasmid DNA Extraction and Isolation.** The GeneJET Miniprep Kit was used to extract plasmid DNA from bacterial cell cultures that had been grown overnight. The bacterial culture, ranging from 1.5 mL to 5 mL, was centrifuged at 13,000 rpm, 5 minutes using a table-top microcentrifuge to create a cell pellet. This pellet was then mixed with 250 $\mu$l of Resuspension Solution containing RNase A to resuspend the cells. Following resuspension, plasmid DNA extraction was conducted on the resuspended cells using the GeneJet Plasmid Miniprep kit, with detailed instructions provided by the manufacturer. Subsequent to extraction, the concentration and purity of the DNA were gauged using a Nanodrop.

**SS6.3. DNA Purification.** The Qiagen PCR purification kit (Qiagen) was employed for the purification of PCR products. This purification was performed as a preliminary step before carrying out subsequent downstream procedures. The DNA purification process involved a series of sequential actions, including binding, washing, and centrifugation using a QIAquick column. The resultant purified DNA was then extracted from the column and transferred into a fresh 1.5 mL microcentrifuge tube utilising Qiagen elution buffer. For a comprehensive protocol, the manufacturer's guidelines can be consulted. In specific instances, the Monarch DNA clean-up kit from NEB was utilised following the instructions provided by the manufacturer.

**SS6.4. Agarose Gel Electrophoresis.** Agarose gel electrophoresis was carried out utilising the wide Mini-Sub Cell GT gel tank manufactured by Bio-Rad. The process involved the use of TAE buffer and visualization was achieved through a UVP Ultraviolet transilluminator emitting light at 365 nm or a blue light transilluminator. DNA samples were combined with 6X DNA Loading Dye from NEB and typically introduced onto agarose gels of 0.5% (w/v) concentration. These gels were prepared by dissolving agarose in 100 mL of TAE buffer and a 1/10000 proportion of SybrSafe Gel Stain 10000X (Thermo Fisher). Adjustments to the agarose concentration were made as needed. When necessary, DNA fragments were isolated from the agarose gels using the GeneJet Gel Extraction Kit from Thermo Fisher Scientific, following the instructions provided by the manufacturer.

**SS6.5. Polymerase Chain Reaction (PCR).** PCR amplification was carried out using either Q5 High-Fidelity Master Mix (NEB) or Phusion HF Master mix (NEB), following the instructions provided by the manufacturers, unless otherwise specified. The essential elements for the PCR mixture were mixed on ice and subsequently moved to an available thermocycler (either the T100 PCR thermal cycler from Bio-Rad or the Eppendorf 5332 Mastercycler from Eppendorf). Following completion of the reaction, the PCR products were placed through agarose gel electrophoresis. The NEB Tm Calculator online tool was utilised to determine the annealing temperature for each primer set. Modification of the extension time was done based on the size of the amplicon, typically allowing for a 30-second extension per kilobase pair of amplified DNA.

**SS6.6. Gibson Assembly.** Gene assembly for the construction of plasmids and cosmids was performed using either a custom Gibson master mix or the NEBuilder HiFi DNA Assembly as stated previously. A typical reaction contained 100 -120 ng of linearised vector and a 3x molar excess of DNA insert topped up to 10 $\mu$L of nuclease-free water before the addition of 10 $\mu$L of 2X NEBuilder HiFi DNA Assembly Master Mix. Reaction mixtures were incubated at 50°C for 1 hour, and a 1 - 2.5 $\mu$L aliquot was used for the transformation of an appropriate competent *E. coli* strain.

**SS6.7. Preparation of chemically competent *E.coli*.** Overnight cultures of each strain were grown in LB medium. For each strain, 10 mL of fresh LB medium was prepared in a 50 mL flask. The bacterial culture was incubated overnight at 37 °C with shaking conditions at 220 rpm. On the following day, a 1% (v/v) of the overnight culture was used to grow a new culture in LB media and incubated at 37 °C until they reached an $OD_{600}$ of 0.5 – 0.6. The cells were harvested by centrifugation (3,000 G, 10 min, 4 °C) and the cell pellet was resuspended in 10 mL ice-cold 30 mM $CaCl_2$ solution. The cells were then centrifuged at 3,000 G for 5 min at 4 °C and suspended in 3 mL ice-cold 85 mM $CaCl_2$ solution containing 10% glycerol. Aliquots of 50 $\mu$L were transferred into 1.5 mL Eppendorf tubes and stored at –80 °C until further use or directly used for transformation.

**SS6.8. Preparation of electro-competent *E.coli*.** Overnight cultures of each strain were grown in LB medium. For each strain, 10 mL of fresh LB medium was prepared in a 50 mL flask. On the following day, a 1% (v/v) of the overnight culture was used to grow a new culture in LB media and incubated at 37 °C until they reached an $OD_{600}$ of 0.5 – 0.6. The cells were then transferred to 15 mL Falcon conical tubes and pelleted by centrifugation for 5 minutes at 4,000 G. After promptly removing and pouring off the supernatant, the cells underwent washing. Each tube received 10 mL of chilled 10% glycerol, and vigorous vortexing was performed to resuspend the pellet. The tubes were then centrifuged for 10 minutes, with supernatant removal following promptly. This washing process was repeated for at least four cycles in 10% glycerol. The cells were resuspended in around 100 $\mu$L of 10% glycerol to create a 100x concentration of the initial culture. These concentrated cell suspensions were divided into 30-50 $\mu$l aliquots in 1.5 mL Eppendorf tubes. Finally, the samples were either frozen or directly used for electroporation.

**SS6.9. Transformation of electro-competent *E. coli*.** An 50 - 100 $\mu$L aliquot of electrocompetent *E. coli* was gently thawed on ice. Around 10 ng of plasmid, cosmid, or 1 - 2.5 $\mu$L of ligation mix was added and mixed. The mix was then moved to a pre-chilled electroporation cuvette and electroporation was done using a Bio-Rad GenePulser Electroporator (settings: 2.5 kV, 100 Ω, and 25 $\mu$F). Immediately after electroporation, 950 $\mu$L of SOC medium (2% tryptone, 0.5% yeast extract, 10 mM NaCl, 2.5 mM KCl, 10 mM MgCl2, 10 mM MgSO4, and 20 mM glucose) was added and the mixture was incubated at 37 °C (or 30ºC in some instances) with agitation for 1 hour. The cells were then spread on LB agar plates with the corresponding antibiotics for selection and incubated overnight at a temperature depending on the strain/plasmid. The following day, single colonies were selected and cultured to extract DNA for verification through sequencing.

**SS6.10. Transformation of chemically-competent *E.coli*.** An 50 - 100 $\mu$L aliquot of electrocompetent *E. coli* was gently thawed on ice. Around 20-50 ng of plasmid, cosmid, or 5 $\mu$L of ligation mix was added and mixed. The mixture was kept on ice for 30 minutes before transferring the tube to a water bath at 42°C for heat shock for 42- 45 seconds. The tube was placed back on ice for 2 - 5 minutes, immediately after, 950 $\mu$L of SOC medium (2% tryptone, 0.5% yeast extract, 10 mM NaCl, 2.5 mM KCl, 10 mM MgCl2, 10 mM MgSO4, and 20 mM glucose) was added and the mixture was incubated at 37 °C (or 30ºC in some instances) with agitation for 1 hour. The cells were then spread on LB agar plates with the corresponding antibiotics for selection and incubated overnight at a temperature depending on the strain/plasmid. The following day, single colonies were selected and cultured to extract DNA for verification through sequencing.

**SS6.11. Protein extraction.** Following protein expression, cell cultures were pelleted at 7000 RPM using a table-top centrifuge for 10 minutes at 4°C. The collected cell pellets were then suspended in Lysis Buffer (50 mM Tris-HCl, 150 mM NaCl, 1% Triton X-100, and 5 mM EDTA) at a 5% (v/v) ratio while maintaining a temperature of 4°C. Subsequently, the bacterial cells, once resuspended, underwent lysis through sonication with the Sonics Vibra-Cell VCX130 ultrasonic processor with sonication settings of amplitude = 80%, pulse on = 10 s, pulse off = 30 s, and time = 3 min. To separate the soluble and insoluble components, the cell lysates were subjected to centrifugation at 20,000 rpm for 30 minutes at 4°C. The resulting soluble fraction was carefully poured off and stored on ice, along with the insoluble fraction. In preparation for SDS PAGE analysis, varying dilutions of either the soluble or insoluble fractions were combined with -mercaptoethanol, and the mixture was boiled for 10 minutes before being loaded onto a running gel.

**SS6.12. SDS PAGE for expression verification.** The protein samples were mixed with 6X protein loading dye, and deposited onto pre-cast 4-20% Novex Gels from Invitrogen. The SDS-PAGE was realised using a specialised Mini-Gel Tank designed for Novex gels by Invitrogen. This procedure utilised a protein gel buffer with a composition of 25 mM Tris, 190 mM glycine, and 3.5 mM SDS at a pH of 8.3. Following electrophoresis, the SDS-PAGE gel was immersed in a tray containing Instant Blue Coomassie stain from Abcam. The gel was gently shaken for one hour for the staining process to complete. Post-staining, the gel was washed overnight with tap water using the same tray before the final visualization step.

#### Supplementary Section SS7: Phage Methods

**SS7.1. Preparation of natural phages.** For the comparison of the effectiveness of P4-EKORhE encoding the multi-lysins cassette respective to natural phages, phages T4 and K1F-GFP were propagated in *Escherichia coli* B cells as host (for T4) and EV36 strains (for K1F-GFP) and recovered by PEG8000 precipitation (see Protocol SS7.2). After the PEG8000 purified phages, a caesium chloride equilibrium centrifugation, followed by dialysis against SM buffer (25 mM Tris-HCl, 8 mM $MgSO_4$, 1M NaCl) and dialysis against SM2 buffer (25 mM Tris-HCl, 8 mM $MgSO_4$, 100 mM NaCl) was effectuated (see Protocol SS7.3). All phage were titered by plaque assay (SS7.4), using *Escherichia coli* B strains (for T4) or EV36 strains (for K1F-GFP), plating O/N at 37 °C. The phages were finally prepared at MOIs of 10 and 100 using SM2 buffer dilutions. K1F is a phage that can particularly infect EV36 strains, the K1F-GFP is a variant of phage K1F that has been genetically modified to contain a Green Fluorescent Protein (GFP) on its capsid for other purposes not related to this work and which are not of relevance for discussion.

**SS7.2. Natural phage propagation.** To propagate T4 or K1F phages, T4 was propagated using B strains of *Escherichia coli*, while K1F was propagated using an EV36 strain. First, *Escherichia coli* strains were incubated in 100 mL of Lysogeny Broth (LB) in a sterile conical flask and incubated overnight at 37 °C on a rotating table. Next, 5 $\mu$L of phage was added to a 10 mL diluted *Escherichia coli* culture with an optical density ($OD_{600}$) of 0.15 in a 50 mL tube. The mixture was incubated at 37 °C with rotation and the culture was observed for clearance every 30 minutes, adjusting the phage concentration or $OD_{600}$ of the *Escherichia coli* culture if needed. Once clearance was achieved, the culture was centrifuged at 3220 G at 4 °C for 15 minutes and the supernatant was transferred to a new tube, which served as the phage solution. A 1 mL of the cleared phage was added to a new *Escherichia coli* culture at a slightly higher $OD_{600}$ and gradually increased the $OD_{600}$ and volume of the new cultures until cleared culture with an $OD_{600}$ > 2.0 and a volume of 1.5 L. The conical flasks were filled only up to one-third of their capacity when using a rotating table. The culture was poured into six 250 mL centrifuge bottles and centrifuged at 4000 G at 4 °C for 15 minutes using rotor JLA-16.250 in an Avanti centrifuge. The supernatant of the culture was then transferred to sterile blue cap bottles, 0.2 M NaCl added, and incubated on ice for 1 hour. Then, a NaCl solution was poured into six 250 mL centrifuge bottles and centrifuged at 5000 g, at 4 °C, for 45 minutes to remove bacterial remnants. The supernatant was transferred to sterile blue cap bottles and 10% w/v PEG8000 was added, ensuring regular agitation until the PEG8000 was dissolved. The solution was incubated on ice for a minimum of 2 hours, or preferably overnight at 4 °C. Subsequently, the PEG8000 solution was poured into six 250 mL centrifuge bottles and centrifuged at 25,000 G at 4 °C for 60 minutes to precipitate the phage particles. Finally, the resulting pellets were resuspended in SM buffer (25 mM Tris-HCl, 8 mM $MgSO_4$, 1M NaCl) with a total volume of approximately 10 mL and the final solution was stored at 4 °C.

**SS7.3. Caesium chloride purification.** Once phages are propagated and recovered in PEG8000, a CsCl density gradient was prepared by mixing CsCl with sterile water to achieve different densities: 9.6 g CsCl + 7.5 mL sterile H2O for a density of 1.7 g/mL, 6.6 g CsCl + 8.0 mL sterile H2O for a density of 1.5 g/mL, and 5.4 g CsCl + 8.5 mL sterile H20 for a density of 1.4 g/mL. CsCl was then added to the phage solution to achieve a density of 1.3 g/mL, aiming for an approximate concentration of 0.5 g CsCl per mL of solution. The density gradient was stacked into a polymer centrifuge tube, starting with the highest density at the bottom. To prevent tube collapse during centrifugation, a maximum gap of 5mm was left at the top, ensuring sufficient space for the phage solution. A balance was prepared, using a CsCl solution with the same density as the gradient. Both the sample and balance tubes were required to be accurate within 0.01 g. The CsCl gradient was then subjected to centrifugation at 150,000 G and 4 °C for 20 hours, utilising rotor SW28 and a Beckman L-90K centrifuge. As a result, a grey-bluish band formed in the middle third of the tube. The band was extracted by piercing a large gauge syringe needle just below it, and the purified phage was stored at 4 °C. Subsequently, a dialysis tube was placed in a beaker under cold running water for 3 hours, approximately 15 cm in length. One end of the dialysis tube was sealed with a bag clip, and the extracted phage band was added to the tube. Care was taken to prevent air from entering, and the other end of the tube was tightly sealed. The dialysis tube, containing the phage, was then placed in a large beaker with approximately 900 mL of SM buffer (25 mM Tris-HCl, 8 mM $MgSO_4$, 1M NaCl), ensuring that the stretch of the tube between the clips was fully submerged. The beaker was left at 4 °C overnight. The dialysis tube was subsequently transferred to another beaker with approximately 900 mL of SM2 buffer (25 mM Tris-HCl, 8 mM $MgSO_4$, 100 mM NaCl), and it was placed on a magnetic stirrer for gentle mixing at room temperature for 2 hours. This step was repeated with a fresh SM2 buffer. After the dialysis process, the purified phage was retrieved and stored in appropriate aliquots at -20 °C.

**SS7.4. Plaque Assay.** For the plaque assay, a culture of *E.coli* is grown overnight (O/N) and refreshed at a 1/10 dilution in LB (Luria-Bertrani) media until $OD_{600}$ = 0.2 - 0.5 at 37ºC, adding 10 mM $Ca^{++}$. A serial dilution of a phage lysate is prepared in LB media. A 200 $\mu$L of a diluted phage sample is mixed 1:1 with 200 $\mu$L of the exponentially growing LB culture of *E.coli*, in replicas of three, and incubated at 37ºC for 10 minutes. Afterwards, the incubated premixed solution of 400 $\mu$L of bacteria-phage with 10 mM $CaCl_2$ is further mixed with 3 mL of semi-solid LB agar (LB with 0.75% agar, at 42ºC). Rapidly afterwards, a pre-incubated Petri dish plate containing 20 mL of solid LBA (Luria-Bertrani media with 1.5 % agar) at 37ºC is used as a base to pour the 3.4 mL solution of bacteria-phage in semi-solid LB agar. Rapidly, the poured semi-solid LB agar is evenly distributed throughout the Petri dish plate. The is let cool down til the semi-solid LB agar containing the mixture of bacteria-phage is solidified. Afterwards, the plate is incubated O/N at 37ºC for the appearance of plaques. A correlation between the sample and its respective dilution was used to quantify the concentration of phage particles present in the diluted sample by counting the number of plaques in the plate.

***SS7.4.1. Spot assay.*** Because the P4-EKORhE system for the production of transducing particles relies on the use of a conditionally propagating P4-phage (*i.e.* the P4-EKORhE in Figure 5), it is possible to use this conditionally propagating phage for the counting of the number of transducing particles, using a spot assay. For the spot assay, a bacterial culture of $\Delta$cos:TriR-P2-c5545 Z1 bacterial cells is grown overnight and refreshed at a 1/10 dilution in LB (Luria-Bertrani) media with 10 $\mu$g/mL Trimethoprim until $OD_{600}$ = 0.2 - 0.5 at 37ºC. Afterwards a premixed solution of 300 $\mu$L of bacterial cells with 0.067 M $CaCl_2$ and further mixed with 3 mL of semi-solid soft-LB-TBA (Terrific-Broth media with 50% premixed LB with 1.5% agar) at 42ºC, rapidly afterwards, a pre-incubated Petri dish containing 20 mL of solid LBA (Luria-Bertrani media with 1.5 % agar) at 37ºC is used as a base to pour the 3.3 mL solution of bacteria in soft-LB-TBA. Rapidly before the poured semi-solid soft-LB-TBA containing the premixed bacterial culture is solidified, multiple 10 $\mu$L drops of a serially diluted sample of P4-EKORhE-based transducing particles lysate are distributed throughout the plate (see Figure S3 and Figure S4), and a correlation between the sample and its respective dilution used to quantify the concentration of transducing particles present in the sample.

#### Supplementary Section SS8: Genetic and aminoacid sequences

##### SS8.1. Primers used in this work.

**Table S4.** Primers used for the construction of P4-EKORhE with the multi-lysins cassette

| Primer | Sequence | Purpose |
|---|---|---|
| TetOLys-BKB-P4-EKORhE-fw FW primer | 5’ tgc tgg atg aat ttt tct aac ttg gaa gta aga atg g 5’ | for the extraction of P4-min ready and assembly with the multi-lysins cassette |
| TetOLys-BKB-P4-EKORhE-rv RV primer | 5’ ttc tgg aat ttg gta ccg agc ttt aac tct ctc atg cca c 3’ | for the extraction of P4-min ready and assembly with the multi-lysins cassette |
| TetOLys-INS-P4-EKORhE-fw FW primer | 5’ gtg gca tga gag agt taa agc tcg gta cca aat tcc aga aaa gag gc 3’ | for extraction of the multi-lysins cassette and assembly with P4-min |
| TetOLys-INS-P4-EKOPRhE-rv RV primer | 5’ gca cca ttc tta ctt cca agt tag aaa aat tca tcc agc atc aga tga aat tgc 3’ | for extraction of the multi-lysins cassette and assembly with P4-min |
| EpsilonKOfwd55 FW primer | 5’ gct cat tca gca caa aat caa ggg gct ttt tta tta cgc ac 3’ | for the selective *knock-out* of the natural epsilon ($\varepsilon$) gene of P4 |
| EpsilonKOrev55 RV primer | 5’ tgc gta ata aaa aag ccc ctt gat ttt gtg ctg aat gag ctg 3’ | for the selective *knock-out* of the natural epsilon ($\varepsilon$) gene of P4 |
| RhamE-EKORhE-P4-BKB-rv RV primer | 5’ tct atc aac agg agt cca agc ttt aac tct ctc atg cca cg 3’ | for extraction of the P4-min cosmid containing the multi-lysins cassette |
| RhamE-EKORhE-P4-BKB-fw FW primer | 5’ tag cgg cct tca ata att ggc tat aaa aat agg cgt atc acg agg 3’ | for extraction of the P4-min cosmid containing the multi-lysins cassette |
| RhamE-EKORhE-P4-INS-fw FW primer: | 5’ gtg gca tga gag agt taa agc ttg gac tcc tgt tga tag atc c 3’ | for extraction of the rhamnose-inducible epsilon($\varepsilon$) cassette |
| RhamE-EKORhE-P4-INS-rv RV primer | 5’ tga tac gcc tat ttt tat agc caa tta ttg aag gcc gct aac g 3’ | for extraction of the rhamnose-inducible epsilon($\varepsilon$) cassette |

##### SS8.2. Mass spectrometry query aminoacid sequences.

```
>Lambda-Rz protein
MSRVTAIISALVICIIVCLSWAVNHYRDNAITYKAQRDKNARELKLANAAITDMQMRQRDVAALDAKYTKEL
ADAKAENDALRDDVAAGRRRLHIKAVCQSVREATTASGVDNAASPRLADTAERDYFTLRERLITMQKQLEGT
QKYINEQCR*
>Lambda_R protein
MVEINNQRKAFLDMLAWSEGTDNGRQKTRNHGYDVIVGGELFTDYSDHPRKLVTLNPKLKSTGAGRYQLLSR
WWDAYRKQLGLKDFSPKSQDAVALQQIKERGALPMIDRGDIRQAIDRCSNIWASLPGAGYGQFEHKADSLIA
KFKEAGGTVREIDV*
>Lambda_S protein
MKMPEKHDLLAAILAAKEQGIGAILAFAMAYLRGRYNGGAFTKTVIDATMCAIIAWFIRDLLDFAGLSSNLA
YITSVFIGYIGTDSIGSLIKRFAAKKAGVEDGRNQ*
>MS2-gpL protein
```

```
METRFPQQSQQTPASTNRRRPFKHEDYPCRRQQRSSTLYVLIFLAIFLSKFTNQLLLSLLEAVIRTVTTLQQ
LLT*
>PhiX174-gpE protein
MVRWTLWDTLAFLLLLSLLLPSLLIMFIPSTFKRPVSSWKALNLRKTLLMASSVRLKPLNCSRLPCVYAQET
LTFLLTQKKTCVKNYVRKE*
MS_query_seqs
>Aminoglycoside 3'-phosphotransferase (aphA1) - Kanamycin resistance protein
MSHIQRETSCSRPRLNSNMDADLYGYKWARDNVGQSGATIYRLYGKPDAPELFLKHGKGSVANDVTDEMVRL
NWLTEFMPLPTIKHFIRTPDDAWLLTTAIPGKTAFQVLEEYPDSGENIVDALAVFLRRLHSIPVCNCPFNSD
RVFRLAQAQSRMNNGLVDASDFDDERNGWPVEQVWKEMHKLLPFSPDSVVTHGDFSLDNLIFDEGKLIGCID
VGRVGIADRYQDLAILWNCLGEFSPSLQKRLFQKYGIDNPDMNKLQFHLMLDEFF*
```

**SS8.3. Rhamnose-inducible epsilon ($\varepsilon$).** Degenerate, codon-optimised, rhamnose-inducible epsilon ($\varepsilon$) genetic sequence:

```
LOCUS       Exported File            698 bp ds-DNA    linear   SYN 29-SEP-2024
DEFINITION  .
ACCESSION   .
VERSION     .
KEYWORDS    rhaB-epsilon_opt
SOURCE      synthetic DNA sequence
  ORGANISM  unspecified
REFERENCE   1  (bases 1 to 698)
  AUTHORS   Robert Ramirez-Garcia, Alfonso Jaramillo
  TITLE     Direct Submission
  JOURNAL   Exported 29 Jul 2024
FEATURES             Location/Qualifiers
     source          1..698
                     /organism="unspecified"
                     /mol_type="genomic DNA"
     terminator      complement(1..103)
                     /note="T0 terminator"
                     /note="color: #ffffff; direction: LEFT"
     misc_feature    1..103
                     /note="T0"
                     /note="color: #a6acb3"
     terminator      5..99
                     /note="lambda t0 terminator"
                     /note="color: #ffffff"
     terminator      complement(5..99)
                     /note="lambda t0 terminator"
                     /note="color: #666699; direction: LEFT"
     terminator      5..99
                     /gene="
                     "
                     /note="lambda t0 terminator"
                     /note="transcription terminator from phage lambda"
                     /note="color: #ffffff"
     terminator      55..89
                     /note="lambda t0r terminator"
                     /note="color: #800080; direction: RIGHT"
     CDS             complement(104..391)
                     /codon_start=1
                     /note="P4 epsilon opt (synthetic)"
                     /note="color: #33cccc"
                     /translation="MRNKKAPQTVSARHDAREHLSIEAYHKLNRASAVSRFVGGDLIHR
                     ELSGLHQLYIPHIFSYLNEDIDFVLNELKAKGLCRDFLAQQKDRGDRTHV"
     misc_feature    104..123
                     /note="v3_Epsilon_INS"
```

```
                     /note="color: #a6acb3"
     RBS             complement(392..416)
                     /note="BBa_B0064 weak RBS"
                     /note="color: #ff0000; direction: LEFT"
     RBS             complement(392..415)
                     /note="BBa_B0064 weak RBS"
                     /note="color: #ff0000; direction: LEFT"
     RBS             complement(398..409)
                     /note="RBS-64"
                     /note="color: #993300; direction: LEFT"
     promoter        complement(423..541)
                     /note="rhaB promoter"
                     /note="This reverse directional feature has 8 segments:
                     1:423..423/#800000/+1
                     2:424..454/#800000
                     3:455..471/#ff9900/RhaSop
                     4:472..487/#800000
                     5:488..504/#ff9900/RhaSop
                     6:505..507/#800000
                     7:508..523/#ff9900/CRPop
                     8:524..541/#800000"
     protein_bind    complement(455..471)
                     /note="RhaS binding site"
                     /note="color: #31849b; direction: LEFT"
     protein_bind    complement(488..504)
                     /note="RhaS binding site"
                     /note="color: #31849b; direction: LEFT"
     protein_bind    complement(508..523)
                     /note="CRP binding site"
                     /note="color: #31849b; direction: LEFT"
     terminator      complement(548..649)
                     /note="TNA21 terminator"
                     /note="color: #ff0000; direction: LEFT"
     terminator      complement(650..698)
                     /note="L3S3P22 strong terminator"
                     /note="color: #993300; direction: LEFT"
     CDS             complement(681..692)
                     /codon_start=1
                     /note="Factor Xa site"
                     /note="color: #cc99b2
                     Cleavage site after base 680"
                     /translation="IEGR"
ORIGIN
        1 cttggactcc tgttgataga tccagtaatg acctcagaac tccatctgga tttgttcaga
       61 acgctcggtt gccgccgggc gtttttttatt ggtgagaatc cagttaaacg tgggtgcgat
      121 cgccgcggtc tttctgctgc gccaggaaat cacggcacag gcctttcgct ttcagctcat
      181 tcagcacgaa gtcgatgtct tcgttcaggt aggagaagat gtgcgggata tacagctggt
      241 gcagaccgct cagttcacgg tggatcaggt cgccgccaac gaagcggctg acggcgctgg
      301 cacggttcag tttgtgatac gcttcgatgg acaggtgttc gcgcgcgtcg tgacgcgcgg
      361 aaacggtctg cggtgctttt ttgttacgca tctagtattt cccctctttc tctagagatc
      421 tccacgacca gtctaaaaag cgcctgaatt cgcgaccttc tcgttactga caggaaaatg
      481 ggccattggc aaccagggaa agatgaacgt gatgatgttc acaatttgct gaattgtggc
      541 cgggcccccg taatgacctt tatacgactg acccaaataa aaaaagccac cgttgcaact
      601 taagagtcac taacggcagc ttatgcgaat agtgttgcca cttgctcaag ggagaccaga
      661 aacaaaaaaa ggccgcgtta gcggccttca ataattgg
//
```

**SS8.4. The multi-lysins cassette cosmid.** Genetic sequence of the multi-lysins cassette cosmid used for Figure S11 and for the construction of the P4-EKORhE:

```
LOCUS       Exported File           4885 bp ds-DNA    circular SYN 09-APR-2019
DEFINITION  .
ACCESSION   .
VERSION     .
KEYWORDS    09042019-TetOlys_MS2_PhiX174_3Lmbd-pSEVA261-Km-p15A
SOURCE      synthetic DNA construct
  ORGANISM  synthetic DNA construct
REFERENCE   1  (bases 1 to 4885)
  AUTHORS   Robert Ramirez-Garcia, Alfonso Jaramillo
  TITLE     Direct Submission
  JOURNAL   Exported 22 Jul 2024
FEATURES             Location/Qualifiers
     source          1..4885
                     /note="color: #ffffff"
     source          1013..2648
                     /note="color: #ffffff"
     source          1013..2648
                     /note="color: #ffffff"
     source          1028..1033
                     /note="color: #ffffff"
     source          1310..1313
                     /note="color: #ffffff"
     source          2556..2603
                     /note="color: #ffffff"
     primer_bind     9..32
                     /note="R24"
                     /note="color: #a020f0; direction: RIGHT"
     misc_feature    44..348
                     /note="P4 packaging region"
                     /note="color: #00ff00"
     promoter        274..320
                     /note="P4-Pgop promoter"
                     /note="This forward directional feature has 6 segments:
                     1:274..282/#993300
                     2:283..288/#993300/-35
                     3:289..304/#993300
                     4:305..310/#993300/-10
                     5:311..319/#993300
                     6:320..320/#993300/+1"
     terminator      349..409
                     /note="L3S2P21 strong terminator"
                     /note="color: #993300; direction: RIGHT"
     terminator      410..470
                     /note="L3S2P21 strong terminator"
                     /note="color: #993300; direction: RIGHT"
     promoter        471..524
                     /note="promoter pLtetO12"
                     /note="color: #ffffff; direction: RIGHT"
     protein_bind    471..489
                     /gene="tetO"
                     /bound_moiety="tetracycline repressor TetR"
                     /note="tet operator"
                     /note="
                     "
                     /note="color: #31849b"
```

```
     protein_bind    496..514
                     /gene="tetO"
                     /bound_moiety="tetracycline repressor TetR"
                     /note="tet operator"
                     /note="
                     "
                     /note="color: #31849b"
     RBS             532..543
                     /note="color: #a6acb3"
     RBS             532..543
                     /note="RBS34min"
                     /note="color: #ff6600"
     promoter        706..759
                     /note="promoter pLtetO12"
                     /note="color: #ffffff; direction: RIGHT"
     protein_bind    706..724
                     /gene="tetO"
                     /bound_moiety="tetracycline repressor TetR"
                     /note="tet operator"
                     /note="
                     "
                     /note="color: #31849b"
     protein_bind    731..749
                     /gene="tetO"
                     /bound_moiety="tetracycline repressor TetR"
                     /note="tet operator"
                     /note="
                     "
                     /note="color: #31849b"
     RBS             767..778
                     /note="color: #a6acb3"
     RBS             767..778
                     /note="RBS34min"
                     /note="color: #ff6600"
     CDS             785..1012
                     /codon_start=1
                     /note="MS2 gpL lysis protein"
                     /note="color: #00ccff"
                     /translation="METRFPQQSQQTPASTNRRRPFKHEDYPCRRQQRSSTLYVLIFLA
                     IFLSKFTNQLLLSLLEAVIRTVTTLQQLLT"
     RBS             1016..1027
                     /note="RBS-64"
                     /note="color: #993300; direction: RIGHT"
     misc_RNA        1028..1033
                     /note="binding site of Lin28a"
                     /note="color: #ff00ff; direction: RIGHT"
     CDS             1034..1309
                     /codon_start=1
                     /note="PhiX174 gpE lysis protein"
                     /note="color: #00ccff"
                     /translation="MVRWTLWDTLAFLLLLSLLLPSLLIMFIPSTFKRPVSSWKALNLR
                     KTLLMASSVRLKPLNCSRLPCVYAQETLTFLLTQKKTCVKNYVRKE"
     misc_feature    1302..1307
                     /note="SD"
                     /note="color: #a6acb3"
     misc_feature    1310..1313
                     /note="intergenic spacer in phi-X174"
```

```
                     /note="color: #a6acb3"
     CDS             1310..1313
                     /codon_start=3
                     /note="D"
                     /note="color: #993366"
                     /translation=""
     CDS             1314..1637
                     /codon_start=1
                     /note="Lambda lysS"
                     /note="color: #00ccff"
                     /translation="MKMPEKHDLLAAILAAKEQGIGAILAFAMAYLRGRYNGGAFTKTV
                     IDATMCAIIAWFIRDLLDFAGLSSNLAYITSVFIGYIGTDSIGSLIKRFAAKKAGVEDG
                     RNQ"
     misc_feature    1607..1612
                     /note="SD?"
                     /note="color: #a6acb3"
     CDS             1621..2097
                     /codon_start=1
                     /note="Lambda lysR"
                     /note="color: #00ccff"
                     /translation="MVEINNQRKAFLDMLAWSEGTDNGRQKTRNHGYDVIVGGELFTDY
                     SDHPRKLVTLNPKLKSTGAGRYQLLSRWWDAYRKQLGLKDFSPKSQDAVALQQIKERGA
                     LPMIDRGDIRQAIDRCSNIWASLPGAGYGQFEHKADSLIAKFKEAGGTVREIDV"
     misc_feature    2080..2085
                     /note="SD?"
                     /note="color: #a6acb3"
     CDS             2094..2555
                     /codon_start=1
                     /note="Lambda Rz"
                     /note="color: #00ccff"
                     /translation="MSRVTAIISALVICIIVCLSWAVNHYRDNAITYKAQRDKNARELK
                     LANAAITDMQMRQRDVAALDAKYTKELADAKAENDALRDDVAAGRRRLHIKAVCQSVRE
                     ATTASGVDNAASPRLADTAERDYFTLRERLITMQKQLEGTQKYINEQCR"
     CDS             2094..2554
                     /codon_start=1
                     /note="Lambda Rz"
                     /note="color: #00ccff"
                     /translation="MSRVTAIISALVICIIVCLSWAVNHYRDNAITYKAQRDKNARELK
                     LANAAITDMQMRQRDVAALDAKYTKELADAKAENDALRDDVAAGRRRLHIKAVCQSVRE
                     ATTASGVDNAASPRLADTAERDYFTLRERLITMQKQLEGTQKYINEQCR"
     terminator      2556..2603
                     /note="T7 gp10 terminator"
                     /note="color: #666699; direction: RIGHT"
     terminator      2604..2633
                     /note="T500 Terminator"
                     /note="color: #993300; direction: RIGHT"
     misc_feature    2634..2648
                     /note="BioBrick suffix"
                     /note="This feature has 3 segments:
                     1:2634..2643/#a6ccff/ChangedSpeItoNheI(com...
                     2:2644..2644/#a6ccff/obliterationofEagI/NotI
                     3:2645..2648/#a6ccff"
     primer_bind     2672..2695
                     /note="F24"
                     /note="color: #a020f0; direction: RIGHT"
     terminator      complement(2702..2804)
                     /note="T0 terminator"
```

```
                     /note="color: #ffffff; direction: LEFT"
     CDS             2932..3747
                     /note="Km"
                     /note="color: #993366; direction: RIGHT"
     rep_origin      complement(3797..4001)
                     /direction=LEFT
                     /note="oriT"
                     /note="color: #ffff00"
     misc_feature    4033..4765
                     /note="p15A ori"
                     /note="color: #a6acb3"
     terminator      complement(4780..4884)
                     /note="T1 terminator"
                     /note="color: #ffffff; direction: LEFT"
ORIGIN
        1 ttaattaaag cggataacaa tttcacacag gaggccgcct aggatgcgtt ttcctgcctc
       61 attttctgca aaccgcgcca ttcccggcgc ggtctgagcg tgtcagtgca actgcattaa
      121 aaccgccccg caaagcgggc gggcgaggcg gggaaagcac cgcgcgcaaa cccagaagtt
      181 agttaattat ttgtgtagtc aaagtgcctt gactacatac ctcgttaata cattggagca
      241 taatgaagaa aatctatggc ctatggtcca aaactgtctt ttttgatggc actatcctga
      301 aaaatatgca aaaaatagat tgatgtaagg tggttcttgt cagtgtcgct cggtaccaaa
      361 ttccagaaaa gaggcctccc gaaagggggg cctttttttcg ttttggtccc tcggtaccaa
      421 attccagaaa agaggcctcc cgaaaggggg gcctttttttc gttttggtcc tccctatcag
      481 tgatagagat tgacatccct atcagtgata gagatactga gcactctaga gaaagaggag
      541 aaaggagata tgaaatatct gctgccgacc gcagcagccg gcttactgct gttagcagcc
      601 cagcccgcta tggccagcgc ccagattcag aaagcagaac agaacgatgt gaaactggca
      661 ccgcctaccg atgtgctggg aaaaccctgg cgactagtga attcgtccct atcagtgata
      721 gagattgaca tccctatcag tgatagagat actgagcact ctagagaaag aggagaaagg
      781 agatatggag acccgattcc ctcagcaatc gcagcaaact ccggcatcta ccaacagacg
      841 ccggccattc aagcatgagg attacccatg tcgaagacaa caaagaagtt caactcttta
      901 tgtattgatc ttcctcgcga tctttctctc gaaatttacc aatcaattgc ttctgtcgct
      961 actggaagcg gtgatccgca cagtgacgac tttacagcaa ttgcttactt aaggtaaaga
     1021 ggggaaagga gatatggtac gctggacttt gtgggatacc ctcgctttcc tcctgttgct
     1081 cagtttattg ctgccgtcat tgctgatcat gttcatcccg tcaacattca aacggcctgt
     1141 ctcatcatgg aaggcgctga atttacggaa aacactgtta atggcgtcga gcgtccggct
     1201 gaagccgctg aattgttcgc gtttaccttg cgtgtacgcg caggaaacac tgacgttctt
     1261 actgacgcag aagaaaacgt gcgtcaaaaa ttacgtgcgg aaggagtgat gtaatgaaga
     1321 tgccagaaaa acatgacctg ttggccgcca ttctcgcggc aaaggaacaa ggcatcgggg
     1381 caatccttgc gtttgcaatg gcgtaccttc gcggcagata taatggcggt gcgtttacaa
     1441 aaacagtaat cgacgcaacg atgtgcgcca ttatcgcctg gttcattcgt gaccttctcg
     1501 acttcgccgg actaagtagc aatctcgctt atataacgag cgtgtttatc ggctacatcg
     1561 gtactgactc gattggttcg cttatcaaac gcttcgctgc taaaaaagcc ggagtagaag
     1621 atggtagaaa tcaataatca acgtaaggcg ttcctcgata tgctggcgtg gtcggaggga
     1681 actgataacg gacgtcagaa aaccagaaat catggttatg acgtcattgt aggcggagag
     1741 ctatttactg attactccga tcaccctcgc aaacttgtca cgctaaaccc aaaactcaaa
     1801 tcaacaggcg ccggacgcta ccagcttctt tcccgttggt gggatgccta ccgcaagcag
     1861 cttggcctga aagacttctc tccgaaaagt caggacgctg tggcattgca gcagattaag
     1921 gagcgtggcg ctttacctat gattgatcgt ggtgatatcc gtcaggcaat cgaccgttgc
     1981 agcaatatct gggcttcact gccgggcgct ggttatggtc agttcgagca taaggctgac
     2041 agcctgattg caaaattcaa agaagcgggc ggaacggtca gagagattga tgtatgagca
     2101 gagtcaccgc gattatctcc gctctggtta tctgcatcat cgtctgcctg tcatgggctg
     2161 ttaatcatta ccgtgataac gccattacct acaaagccca gcgcgacaaa aatgccagag
     2221 aactgaagct ggcgaacgcg gcaattactg acatgcagat gcgtcagcgt gatgttgctg
     2281 cgctcgatgc aaaatacacg aaggagttag ctgatgctaa agctgaaaat gatgctctgc
     2341 gtgatgatgt tgccgctggt cgtcgtcggt tgcacatcaa agcagtctgt cagtcagtgc
     2401 gtgaagccac caccgcctcc ggcgtggata atgcagcctc cccccgactg gcagacaccg
     2461 ctgaacggga ttatttcacc ctcagagaga ggctgatcac tatgcaaaaa caactggaag
```

```
     2521 gaacccagaa gtatattaat gagcagtgca gataactagc ataacccctt ggggcctcta
     2581 aacgggtctt gaggggtttt ttgagacaaa caaaagaatg gaatcaaagt taatgctagc
     2641 agccgccgca ggcatgcaag cttgcggccg cgtcgtgact gggaaaaccc tggcgactag
     2701 tcttggactc ctgttgatag atccagtaat gacctcagaa ctccatctgg atttgttcag
     2761 aacgctcggt tgccgccggg cgttttttat tggtgagaat ccaggggtcc ccaataatta
     2821 cgatttaaat ttgtgtctca aaatctctga tgttacattg cacaagataa aaatatatca
     2881 tcatgaacaa taaaactgtc tgcttacata aacagtaata caaggggtgt tatgagccat
     2941 attcagcgtg aaacgagctg tagccgtccg cgtctgaaca gcaacatgga tgcggatctg
     3001 tatggctata aatgggcgcg tgataacgtg ggtcagagcg gcgcgaccat ttatcgtctg
     3061 tatggcaaac cggatgcgcc ggaactgttt ctgaaacatg gcaaaggcag cgtggcgaac
     3121 gatgtgaccg atgaaatggt gcgtctgaac tggctgaccg aatttatgcc gctgccgacc
     3181 attaaacatt ttattcgcac cccggatgat gcgtggctgc tgaccaccgc gattccgggc
     3241 aaaaccgcgt ttcaggtgct ggaagaatat ccggatagcg gcgaaaacat tgtggatgcg
     3301 ctggccgtgt ttctgcgtcg tctgcatagc attccggtgt gcaactgccc gtttaacagc
     3361 gatcgtgtgt ttcgtctggc ccaggcgcag agccgtatga acaacggcct ggtggatgcg
     3421 agcgattttg atgatgaacg taacggctgg ccggtggaac aggtgtggaa agaaatgcat
     3481 aaactgctgc cgtttagccc ggatagcgtg gtgacccacg gcgattttag cctggataac
     3541 ctgattttcg atgaaggcaa actgattggc tgcattgatg tgggccgtgt gggcattgcg
     3601 gatcgttatc aggatctggc cattctgtgg aactgcctgg gcgaatttag cccgagcctg
     3661 caaaaacgtc tgtttcagaa atatggcatt gataatccgg atatgaacaa actgcaattt
     3721 catctgatgc tggatgaatt tttctaataa ttaattggac cgcggtccgc gcgttgtcct
     3781 tttccgctgc ataaccctgc ttcggggtca ttatagcgat tttttcggta tatccatcct
     3841 ttttcgcacg atatacagga ttttgccaaa gggttcgtgt agactttcct tggtgtatcc
     3901 aacggcgtca gccgggcagg ataggtgaag taggcccacc cgcgagcggg tgttccttct
     3961 tcactgtccc ttattcgcac ctggcggtgc tcaacgggaa tcctgctctg cgaggctggc
     4021 cgtaggccgg ccctagaaat attttatctg attaataaga tgatcttctt gagatcgttt
     4081 tggtctgcgc gtaatctctt gctctgaaaa cgaaaaaacc gccttgcagg gcggtttttc
     4141 gaaggttctc tgagctacca actctttgaa ccgaggtaac tggcttggag gagcgcagtc
     4201 accaaaactt gtcctttcag tttagcctta accggcgcat gacttcaaga ctaactcctc
     4261 taaatcaatt accagtggct gctgccagtg gtgcttttgc atgtctttcc gggttggact
     4321 caagacgata gttaccggat aaggcgcagc ggtcggactg aacggggggt tcgtgcatac
     4381 agtccagctt ggagcgaact gcctacccgg aactgagtgt caggcgtgga atgagacaaa
     4441 cgcggccata acagcggaat gacaccggta aaccgaaagg caggaacagg agagcgcacg
     4501 agggagccgc cagggggaaa cgcctggtat ctttatagtc ctgtcgggtt tcgccaccac
     4561 tgatttgagc gtcagatttc gtgatgcttg tcaggggggc ggagcctatg gaaaaacggc
     4621 ttttgccgcg gccctctcac ttccctgtta agtatcttcc tggcatcttc caggaaatct
     4681 ccgccccgtt cgtaagccat ttccgctcgc cgcagtcgaa cgaccgagcg tagcgagtca
     4741 gtgagcgagg aagcggaata tatccggcgc gcccagctgt ctagggcggc ggatttgtcc
     4801 tactcaggag agcgttcacc gacaaacaac agataaaacg aaaggcccag tctttcgact
     4861 gagcctttcg ttttatttga tgcct
//
```

Trace sequence "90DJ25_61971875_61971875.ab1":

```
>90DJ25_61971875_61971875.ab1  (1604 bp)
GGGGAATCCGACAACACAGATAAACGAAAGGCCCAGTCTTTCGACTGAGCCTTTCGTTTTATTTGATGCCTTTAATTA
AAGCGGATAACAATTTCACACAGGAGGCCGCCTAGGATGCGTTTTCCTGCCTCATTTTCTGCAAACCGCGCCATTCCC
GGCGCGGTCTGAGCGTGTCAGTGCAACTGCATTAAAACCGCCCCGCAAAGCGGGCGGGCGAGGCGGGGAAAGCACCGC
GCGCAAACCCAGAAGTTAGTTAATTATTTGTGTAGTCAAAGTGCCTTGACTACATACCTCGTTAATACATTGGAGCAT
AATGAAGAAAATCTATGGCCTATGGTCCAAAACTGTCTTTTTTGATGGCACTATCCTGAAAAATATGCAAAAAATAGA
TTGATGTAAGGTGGTTCTTGTCAGTGTCGCTCGGTACCAAATTCCAGAAAAGAGGCCTCCCGAAAGGGGGGCCTTTTT
TCGTTTTGGTCCCTCGGTACCAAATTCCAGAAAAGAGGCCTCCCGAAAGGGGGGCCTTTTTTCGTTTTGGTCCTCCCT
ATCAGTGATAGAGATTGACATCCCTATCAGTGATAGAGATACTGAGCACTCTAGAGAAAGAGGAGAAAGGAGATATGA
AATATCTGCTGCCGACCGCAGCAGCCGGCTTACTGCTGTTAGCAGCCCAGCCCGCTATGGCCAGCGCCCAGATTCAGA
AAGCAGAACAGAACGATGTGAAACTGGCACCGCCTACCGATGTGCTGGGAAAACCCTGGCGACTAGTGAATTCGTCCC
TATCAGTGATAGAGATTGACATCCCTATCAGTGATAGAGATACTGAGCACTCTAGAGAAAGAGGAGAAAGGAGATATG
GAGACCCGATTCCCTCAGCAATCGCAGCAAACTCCGGCATCTACCAACAGACGCCGGCCATTCAAGCATGAGGATTAC
CCATGTCGAAGACAACAAAGAAGTTCAACTCTTTATGTATTGATCTTCCTCGCGATCTTTCTCTCGAAATTTACCAAT
```

```
CAATTGCTTCTGTCGCTACTGGAAGCGGTGATCCGCACAGTGACGACTTTACAGCAATTGCTTACTTAAGGTAAAGAG
GGGAAAGGAGATATGGTACGCTGGACTTTGTGGGATACCCTCGCTTTCCTCCTGTTGCTCAGTTTATTGCTGCCGTCA
TTGCTGATCATGTTCATCCCGTCAACATTCAAACGGCCTGTCTCATCATGGAAGGCGCTGAATTTACGGAAAACCTGT
TAATGGCGTCAAGCGTCCGGCTGAACCGCTGAATGTTCCCGTTACCTTGCGGGTCGCCCGGAAACCTGACTTCTTACG
GACCAAAAGAAAACGTGGTCAAAATTACGTGGGGAAGGGGTGGTAAAGAAAAGCCCAAAAAAAAACCCGTTTGCCCCC
TTTCTCCGGCAAAAAAAAAGGCTGGGGACACCTTGTTTTTGGGGGGCCCCTCTCGGCAAAATAGGGGGGTGGCTTTAA
AAAAAATAAAAAACCACGAAGGGCCCCTTTACCGTGTTTCCCGTCCTCCTCTCCCCGAAAAAAAACCCTCTTAATAAA
GGGTGTTTCCTCGTCGTTGTGGGTCACCCTCTCAAAAAAAAAAA
```

##### SS8.5. Original P4-EKORhE with the multi-lysins cassette.

```
LOCUS       Exported File          11439 bp ds-DNA    circular SYN 29-JUL-2024
DEFINITION  .
ACCESSION   .
VERSION     .
KEYWORDS    26082021-ORIGINAL-P4-EKORhE-multilysins
SOURCE      synthetic DNA construct
  ORGANISM  synthetic DNA construct
REFERENCE   1  (bases 1 to 11439)
  AUTHORS   Robert Ramirez-Garcia
  TITLE     Direct Submission
  JOURNAL   Exported 29 Jul 2024
COMMENT     Designed by Robert Ramirez-Garcia on 26-8-2021
FEATURES             Location/Qualifiers
     source          1..11439
                     /organism="synthetic DNA construct"
                     /mol_type="other DNA"
     source          1044..4153
                     /note="color: #ffffff"
     source          1044..3054
                     /note="color: #ffffff"
     source          1044..3054
                     /note="color: #ffffff"
     source          1434..1439
                     /note="color: #ffffff"
     source          1716..1719
                     /note="color: #ffffff"
     source          2962..3009
                     /note="color: #ffffff"
     misc_feature    3..307
                     /note="P4 packaging region"
                     /note="color: #00ff00"
     misc_feature    101..119
                     /note="P2 cos site"
                     /note="color: #a6acb3"
     promoter        233..279
                     /note="P4-Pgop promoter"
                     /note="This forward directional feature has 6 segments:
                     1:233..241/#993300
                     2:242..247/#993300/-35
                     3:248..263/#993300
                     4:264..269/#993300/-10
                     5:270..278/#993300
                     6:279..279/#993300/+1"
     misc_feature    346..448
                     /note="T0"
                     /note="color: #a6acb3"
     terminator      complement(350..444)
```

```
                     /note="lambda t0 terminator"
                     /note="color: #666699; direction: LEFT"
     terminator      350..444
                     /note="lambda t0 terminator"
                     /note="color: #ffffff"
     terminator      400..434
                     /note="lambda t0r terminator"
                     /note="color: #800080; direction: RIGHT"
     CDS             complement(449..736)
                     /codon_start=1
                     /note="P4 epsilon opt (synthetic)"
                     /note="color: #33cccc"
                     /translation="MRNKKAPQTVSARHDAREHLSIEAYHKLNRASAVSRFVGGDLIHR
                     ELSGLHQLYIPHIFSYLNEDIDFVLNELKAKGLCRDFLAQQKDRGDRTHV"
     misc_feature    449..468
                     /note="v3_Epsilon_INS"
                     /note="color: #a6acb3"
     RBS             complement(737..761)
                     /note="BBa_B0064 weak RBS"
                     /note="color: #ff0000; direction: LEFT"
     RBS             complement(737..760)
                     /note="BBa_B0064 weak RBS"
                     /note="color: #ff0000; direction: LEFT"
     promoter        complement(768..886)
                     /note="rhaB promoter"
                     /note="This reverse directional feature has 8 segments:
                     1:768..768/#800000/+1
                     2:769..799/#800000
                     3:800..816/#ff9900/RhaSop
                     4:817..832/#800000
                     5:833..849/#ff9900/RhaSop
                     6:850..852/#800000
                     7:853..868/#ff9900/CRPop
                     8:869..886/#800000"
     protein_bind    complement(800..816)
                     /note="RhaS binding site"
                     /note="color: #31849b; direction: LEFT"
     protein_bind    complement(833..849)
                     /note="RhaS binding site"
                     /note="color: #31849b; direction: LEFT"
     protein_bind    complement(853..868)
                     /note="CRP binding site"
                     /note="color: #31849b; direction: LEFT"
     misc_feature    884
                     /note="G>T Mutation in BQK537"
                     /note="color: #000000"
     terminator      complement(893..994)
                     /note="TNA21 terminator"
                     /note="color: #ff0000; direction: LEFT"
     terminator      complement(995..1043)
                     /note="L3S3P22 strong terminator"
                     /note="color: #993300; direction: LEFT"
     CDS             complement(1026..1037)
                     /codon_start=1
                     /note="Factor Xa site"
                     /note="color: #cc99b2
                     Cleavage site after base 1025"
```

```
                     /translation="IEGR"
     misc_feature    complement(1044..3077)
                     /note="MCS"
                     /note="color: #a6acb3; direction: LEFT"
     misc_feature    1044..3054
                     /note="gBlock tet-lysis"
                     /note="color: #a6acb3"
     terminator      1044..1104
                     /note="L3S2P21 strong terminator"
                     /note="color: #993300; direction: RIGHT"
     misc_feature    1105..1111
                     /note="BioBrick prefix"
                     /note="color: #99ccff"
     misc_feature    1112..1165
                     /note="PLtetO1"
                     /note="color: #a6acb3; direction: RIGHT"
     protein_bind    1112..1130
                     /gene="tetO"
                     /bound_moiety="tetracycline repressor TetR"
                     /note="tet operator"
                     /note="
                     "
                     /note="color: #31849b"
     protein_bind    1137..1155
                     /gene="tetO"
                     /bound_moiety="tetracycline repressor TetR"
                     /note="tet operator"
                     /note="
                     "
                     /note="color: #31849b"
     RBS             1166..1184
                     /note="RBS-34"
                     /note="This forward directional feature has 2 segments:
                     1:1166..1172/#993300/biobrickscar1
                     2:1173..1184/#993300/RBS"
     RBS             1173..1184
                     /note="RBS34min"
                     /note="color: #ff6600"
     misc_RNA        1185..1190
                     /note="binding site of Lin28a"
                     /note="color: #ff00ff; direction: RIGHT"
     CDS             1191..1418
                     /codon_start=1
                     /note="MS2 gpL lysis protein"
                     /note="color: #00ccff"
                     /translation="METRFPQQSQQTPASTNRRRPFKHEDYPCRRQQRSSTLYVLIFLA
                     IFLSKFTNQLLLSLLEAVIRTVTTLQQLLT"
     misc_feature    1196
                     /note="Silent mutation"
                     /note="color: #a6acb3"
     RBS             1422..1433
                     /note="RBS-64"
                     /note="color: #993300; direction: RIGHT"
     misc_RNA        1434..1439
                     /note="binding site of Lin28a"
                     /note="color: #ff00ff; direction: RIGHT"
     CDS             1440..1715
```

```
                     /codon_start=1
                     /note="PhiX174 gpE lysis protein"
                     /note="color: #00ccff"
                     /translation="MVRWTLWDTLAFLLLLSLLLPSLLIMFIPSTFKRPVSSWKALNLR
                     KTLLMASSVRLKPLNCSRLPCVYAQETLTFLLTQKKTCVKNYVRKE"
     misc_feature    1708..1713
                     /note="SD"
                     /note="color: #a6acb3"
     CDS             1716..1719
                     /codon_start=3
                     /note="D"
                     /note="color: #993366"
                     /translation=""
     misc_feature    1716..1719
                     /note="intergenic spacer in phi-X174"
                     /note="color: #a6acb3"
     CDS             1720..2043
                     /codon_start=1
                     /note="Lambda lysS"
                     /note="color: #00ccff"
                     /translation="MKMPEKHDLLAAILAAKEQGIGAILAFAMAYLRGRYNGGAFTKTV
                     IDATMCAIIAWFIRDLLDFAGLSSNLAYITSVFIGYIGTDSIGSLIKRFAAKKAGVEDG
                     RNQ"
     misc_feature    2013..2018
                     /note="SD?"
                     /note="color: #a6acb3"
     CDS             2027..2503
                     /codon_start=1
                     /note="Lambda lysR"
                     /note="color: #00ccff"
                     /translation="MVEINNQRKAFLDMLAWSEGTDNGRQKTRNHGYDVIVGGELFTDY
                     SDHPRKLVTLNPKLKSTGAGRYQLLSRWWDAYRKQLGLKDFSPKSQDAVALQQIKERGA
                     LPMIDRGDIRQAIDRCSNIWASLPGAGYGQFEHKADSLIAKFKEAGGTVREIDV"
     misc_feature    2486..2491
                     /note="SD?"
                     /note="color: #a6acb3"
     CDS             2500..2961
                     /codon_start=1
                     /note="Lambda Rz"
                     /note="color: #00ccff"
                     /translation="MSRVTAIISALVICIIVCLSWAVNHYRDNAITYKAQRDKNARELK
                     LANAAITDMQMRQRDVAALDAKYTKELADAKAENDALRDDVAAGRRRLHIKAVCQSVRE
                     ATTASGVDNAASPRLADTAERDYFTLRERLITMQKQLEGTQKYINEQCR"
     CDS             2500..2960
                     /codon_start=1
                     /note="Lambda Rz"
                     /note="color: #00ccff"
                     /translation="MSRVTAIISALVICIIVCLSWAVNHYRDNAITYKAQRDKNARELK
                     LANAAITDMQMRQRDVAALDAKYTKELADAKAENDALRDDVAAGRRRLHIKAVCQSVRE
                     ATTASGVDNAASPRLADTAERDYFTLRERLITMQKQLEGTQKYINEQCR"
     terminator      2962..3009
                     /note="T7 gp10 terminator"
                     /note="color: #666699; direction: RIGHT"
     terminator      3010..3039
                     /note="T500 Terminator"
                     /note="color: #993300; direction: RIGHT"
     misc_feature    3040..3054
```

```
                     /note="BioBrick suffix"
                     /note="This feature has 3 segments:
                     1:3040..3049/#a6ccff/ChangedSpeItoNheI(com...
                     2:3050..3050/#a6ccff/obliterationofEagI/NotI
                     3:3051..3054/#a6ccff"
     primer_bind     3078..3101
                     /note="F24"
                     /note="color: #a020f0; direction: RIGHT"
     terminator      complement(3108..3210)
                     /note="T0 terminator"
                     /note="color: #ffffff; direction: LEFT"
     terminator      3112..3206
                     /gene="
                     "
                     /note="lambda t0 terminator"
                     /note="transcription terminator from phage lambda"
                     /note="color: #ffffff"
     promoter        3239..3337
                     /note="KanR-aph(3’)-Ia promoter"
                     /note="color: #808000; direction: RIGHT"
     CDS             3338..4153
                     /note="Km"
                     /note="color: #993366; direction: RIGHT"
     CDS             complement(4186..4521)
                     /codon_start=1
                     /transl_table=11
                     /locus_tag="P4p05"
                     /product="hypothetical protein"
                     /note="P4p05"
                     /note="Predicted by GeneMark"
                     /note="color: #993366"
                     /db_xref="GeneID:1261089"
                     /protein_id="NP_597796.1"
                     /translation="MSHIGRMPEVKNRMFTLHPLFTTYHSEIKGENRKVNSVNSSFEKK
                     FFMWIKDALVWIRATDPNLLIKNDLTFRFIDGLKIDWSEKQWGHKRGHISLLCFIICFL
                     SLTYFDV"
     misc_feature    4381..4680
                     /note="crr"
                     /note="crr (cis required region for replication)"
                     /note="color: #a6acb3"
     repeat_region   4381..4500
                     /note="crr 120bp direct repeat"
                     /note="color: #a6acb3"
     repeat_region   4561..4680
                     /note="crr 120bp direct repeat"
                     /note="color: #a6acb3"
     gene            complement(4726..7059)
                     /locus_tag="P4p06"
                     /note="P4p06"
                     /note="color: #a6acb3; direction: LEFT"
                     /db_xref="GeneID:1261095"
     CDS             complement(4726..7059)
                     /codon_start=1
                     /transl_table=11
                     /note="P4p06"
                     /note="color: #993366"
                     /translation="MKMNVTATVSHALGHWPRILPALGIQVLKNRHQPCPVCGGSDRFR
```

```
                     FDDREGRGTWYCNQCGAGDGLKLVEKVFGVSPSDAAAKVAAVTGSLPPADPAVTTAAVD
                     ETDAARKNAAALAQTLMAKTRTGTGNAYLTRKGFPGRECRMLTGTHRAGGVSWRAGDLV
                     VPLYDDSGELVNLQLISADGRKRTLKGGQVRGTCHTLEGQNQAGKRLWIAEGYATALTV
                     HHLTGETVMVALSSVNLLSLASLARQKHPACQIVLAADRDLSGDGQKKAAAAADACEGV
                     VALPPVFGDWNDAFTQYGGEATRKAIYDAIRPPAESPFDTMSEAEFSAMSTSEKAMRIY
                     EHYGEALAVDANGQLLSRYENGVWKVLPPQDFARDVAGLFQRLRAPFSSGKVASVVDTL
                     KLIIPQQEAPSRRLIGFRNGVLDTQNGTFHPHSPSHWMRTLCDVDFTPPVDGETLETHA
                     PAFWRWLDRAAGGRAEKRDVILAALFMVLANRYDWQLFLEVTGPGGSGKSIMAEIATLL
                     AGEDNATSATIETLESPRERAALTGFSLIRLPDQEKWSGDGAGLKAITGGDAVSVDPKY
                     RDAYSTHIPAVILAVNNNPMRFTDRSGGVSRRRVIIHFPEQIAPQERDPQLKDKITREL
                     AVIVRHLMQKFSDPMLARSLLQSQQNSDEALNIKRDADPTFDFIGYLETLPQTSGMYMG
                     NASIIPRNYRKYLYHAYLAYMEANGYRNVLSLKMFGLGLPVMLKEYGLNYEKRHTKQGI
                     QTNLTLKEESYGDWLPKCDDPTTA"
     CDS             complement(7074..7394)
                     /codon_start=1
                     /transl_table=11
                     /locus_tag="P4p07"
                     /product="hypothetical protein"
                     /note="P4p07"
                     /note="ORF106 (AA 1-106)"
                     /note="color: #993366"
                     /db_xref="UniProtKB/Swiss-Prot:P10278"
                     /db_xref="GeneID:1261093"
                     /protein_id="NP_042037.1"
                     /translation="MKTPLPPVLRAALYRRAVACAWLTVCERQHRYPHLTLESLEAAIA
                     AELEGFYLRQHGEEKGRQIACALLEDLMESGPLKAAPSLSFLGLVVMDELCARHIKAPV
                     LH"
     CDS             complement(7530..7985)
                     /codon_start=1
                     /transl_table=11
                     /locus_tag="P4p08"
                     /product="hypothetical protein"
                     /note="P4p08"
                     /note="ORF151 (AA 1-151)"
                     /note="color: #993366"
                     /db_xref="UniProtKB/Swiss-Prot:P05464"
                     /db_xref="GeneID:1261086"
                     /protein_id="NP_042038.1"
                     /translation="MFDFPQPGEIYRSAGFPDVAVVGILEDGIPWEMPYRCPDIVWNPY
                     RRKFSILVRILADGRTTDIPLGRFLREFTCDRPDLFKRSPVNRHAVLKEMAGDPELQKW
                     REKYLDIYPQDTVPVSRAAPVAREWREIPRTEPDPETTPDNSYRNYL"
     misc_feature    7874..7882
                     /note="GSG linker"
                     /note="color: #a6acb3"
     CDS             complement(7978..8087)
                     /codon_start=1
                     /note="P4-gp9 epsilon (KO)"
                     /note="color: #ff0000"
                     /translation="RNKKAPLILC*MS*KPKACAAIFSPSRKTGETGRMF"
     misc_feature    8047..8069
                     /note="DISPENSIBLE"
                     /note="color: #000000"
     CDS             complement(8083..8499)
                     /codon_start=1
                     /transl_table=11
                     /locus_tag="P4p10"
                     /product="putative CI repressor"
```

```
                     /note="P4p10"
                     /note="cI gene product (AA 1-137)"
                     /note="color: #993366"
                     /db_xref="UniProtKB/Swiss-Prot:P05462"
                     /db_xref="GeneID:1261091"
                     /protein_id="NP_042040.1"
                     /translation="MMVWCVVSRADGIPCILPASAHYAAESMVAQAGQPPGWPVSCEAG
                     ILTPVWAIAIERENSGDSVICYSQEAAIMATTLTPSHPEFVFVFAAVRRADRHPRICML
                     RTVAGDERSARRSLVRDYVLSLAARLPVVEVSRA"
     promoter        complement(8608..8678)
                     /note="pLE P4 promoter"
                     /note="This reverse directional feature has 6 segments:
                     1:8608..8608/#808000/+1
                     2:8609..8613/#808000
                     3:8614..8619/#808000/-10
                     4:8620..8637/#808000
                     5:8638..8643/#808000/-35
                     6:8644..8678/#808000"
     CDS             complement(8679..8945)
                     /codon_start=1
                     /transl_table=11
                     /locus_tag="P4p11"
                     /product="transcriptional regulator"
                     /note="P4p11"
                     /note="ORF88 product (AA 1-88) (put. DNA-binding protein)"
                     /note="color: #993366"
                     /db_xref="GOA:P12552"
                     /db_xref="UniProtKB/Swiss-Prot:P12552"
                     /db_xref="GeneID:1261090"
                     /protein_id="NP_042041.1"
                     /translation="MQAVFSSPSPAPVTPLMPLPDITQERFLRVPEVMHLCGLSRSTIY
                     ELIRKGEFPPQVSLGGKNVAWLHSEVTAWMAGRIAGRKRGYDA"
     rep_origin      9001..9724
                     /note="origin of DNA replication"
                     /note="color: #ffff00"
     promoter        complement(9002..9148)
                     /note="P4-pLL promoter"
                     /note="This reverse directional feature has 8 segments:
                     1:9002..9002/#800000/+1
                     2:9003..9008/#800000
                     3:9009..9014/#800000/-10
                     4:9015..9029/#800000
                     5:9030..9035/#800000/-35
                     6:9036..9040/#800000
                     7:9041..9070/#ff6600/Ogr/deltaop
                     8:9071..9148/#800000/Coxbindingoperators"
     promoter        complement(9002..9148)
                     /note="P4-pLL promoter"
                     /note="This reverse directional feature has 8 segments:
                     1:9002..9002/#800000/+1
                     2:9003..9008/#800000
                     3:9009..9014/#800000/-10
                     4:9015..9029/#800000
                     5:9030..9035/#800000/-35
                     6:9036..9040/#800000
                     7:9041..9070/#ff6600/Ogr/deltaop
                     8:9071..9148/#800000/Coxbindingoperators"
```

```
     repeat_region   9091..9100
                     /note="ori type 2 repeat"
                     /note="color: #a6acb3"
     repeat_region   9101..9110
                     /note="ori type 2 repeat"
                     /note="color: #a6acb3"
     repeat_region   9111..9120
                     /note="ori type 2 repeat"
                     /note="color: #a6acb3"
     promoter        9402..9475
                     /note="P4-sid promoter"
                     /note="This forward directional feature has 11 segments:
                     1:9402..9407/#800000
                     2:9408..9425/#ff9900/op1
                     3:9426..9436/#800000
                     4:9437..9442/#800000/-35
                     5:9443..9445/#800000
                     6:9446..9459/#ff9900/op2
                     7:9460..9463/#ff9900/-10
                     8:9464..9465/#800000/-10
                     9:9466..9471/#800000
                     10:9472..9472/#800000/+1
                     11:9473..9475/#800000"
     gene            9472..11375
                     /locus_tag="P4s05"
                     /note="P4s05"
                     /note="color: #a6acb3; direction: RIGHT"
                     /db_xref="GeneID:1261100"
     precursor_RNA   9472..11375
                     /locus_tag="P4s05"
                     /note="late delta regulated transcript of sid through psu"
                     /note="color: #a6acb3; direction: RIGHT"
                     /db_xref="GeneID:1261100"
     gene            9498..10232
                     /note="P4p12"
                     /note="color: #a6acb3; direction: RIGHT"
     CDS             9498..10232
                     /codon_start=1
                     /note="P4-gp12 sid"
                     /note="color: #993366"
                     /translation="MSDHTIPEYLQPALAQLEKARAAHLENARLMDETVTAIERAEQEK
                     NALAQADGNDADDWRTAFRAAGGVLSDELKQRHIERVARRELVQEYDNLAVVLNFERER
                     LKGACDSTATAYRKAHHHLLSLYAEHELEHALNETCEALVRAMHLSILVQENPLANTTG
                     HQGYVAPEKAVMQQVKSSLEQKIKQMQISLTGEPVLRLTGLSAATLPHMDYEVAGTPAQ
                     RKVWQDKIDQQGAELKARGLLS"
     gene            10229..10729
                     /locus_tag="P4p13"
                     /note="P4p13"
                     /note="color: #a6acb3; direction: RIGHT"
                     /db_xref="GeneID:1261088"
     CDS             10229..10729
                     /codon_start=1
                     /note="P4-gp13 delta"
                     /note="color: #993366"
                     /translation="MIYCPSCGHVAHTRRAHFMDDGTKIMIAQCRNIYCSATFEASESF
                     FSDSKDSGMEYISGKQRYRDSLTSASCGMKRPKRMLVTGYCCRRCKGLALSRTSRRLSQ
                     EVTERFYVCTDPGCGLVFKTLQTINRFIVRPVTPDELAERLHEKQELPPVRLKTQSYSL
```

```
                     RLE"
     CDS             10803..11375
                     /codon_start=1
                     /transl_table=11
                     /locus_tag="P4p14"
                     /product="amber mutation-suppressing protein"
                     /note="P4p14"
                     /note="psu gene product (AA 1-190)"
                     /note="color: #993366"
                     /db_xref="UniProtKB/Swiss-Prot:P05460"
                     /db_xref="GeneID:1261094"
                     /protein_id="NP_042044.1"
                     /translation="MESTALQQAFDTCQNNKAAWLQRKNELAAAEQEYLRLLSGEGRNV
                     SRLDELRNIIEVRKWQVNQAAGRYIRSHEAVQHISIRDRLNDFMQQHGTALAAALAPEL
                     MGYSELTAIARNCAIQRATDALREALLSWLAKGEKINYSAQDSDILTTIGFRPDVASVD
                     DSREKFTPAQNMIFSRKSAQLASRQSV"
     gene            10803..11375
                     /locus_tag="P4p14"
                     /note="P4p14"
                     /note="color: #a6acb3; direction: RIGHT"
                     /db_xref="GeneID:1261094"
     CDS             10806..11375
                     /codon_start=1
                     /note="P4-gp14 psu"
                     /note="color: #993366"
                     /translation="ESTALQQAFDTCQNNKAAWLQRKNELAAAEQEYLRLLSGEGRNVS
                     RLDELRNIIEVRKWQVNQAAGRYIRSHEAVQHISIRDRLNDFMQQHGTALAAALAPELM
                     GYSELTAIARNCAIQRATDALREALLSWLAKGEKINYSAQDSDILTTIGFRPDVASVDD
                     SREKFTPAQNMIFSRKSAQLASRQSV"
     terminator      11376..11439
                     /note="tsid terminator"
                     /note="color: #000080; direction: RIGHT"
ORIGIN
        1 gcatgcgttt tcctgcctca ttttctgcaa accgcgccat tcccggcgcg gtctgagcgt
       61 gtcagtgcaa ctgcattaaa accgccccgc aaagcgggcg ggcgaggcgg ggaaagcacc
      121 gcgcgcaaac cgacaagtta gttaattatt tgtgtagtca aagtgccttc agtacatacc
      181 tcgttaatac attggagcat aatgaagaaa atctatggcc tatggtccaa aactgtcttt
      241 tttgatggca ctatcctgaa aaatatgcaa aaaatagatt gatgtaaggt ggttcttgtc
      301 agtgtcgcaa gatccttaag aattcgtggc atgagagagt taaagcttgg actcctgttg
      361 atagatccag taatgacctc agaactccat ctggatttgt tcagaacgct cggttgccgc
      421 cgggcgtttt ttattggtga gaatccagtt aaacgtgggt gcgatcgccg cggtctttct
      481 gctgcgccag gaaatcacgg cacaggcctt tcgctttcag ctcattcagc acgaagtcga
      541 tgtcttcgtt caggtaggag aagatgtgcg ggatatacag ctggtgcaga ccgctcagtt
      601 cacggtggat caggtcgccg ccaacgaagc ggctgacggc gctggcacgg ttcagtttgt
      661 gatacgcttc gatggacagg tgttcgcgcg cgtcgtgacg cgcggaaacg gtctgcggtg
      721 ctttttttgtt acgcatctag tatttcccct ctttctctag agatctccac gaccagtcta
      781 aaaagcgcct gaattcgcga ccttctcgtt actgacagga aaatgggcca ttggcaacca
      841 gggaaagatg aacgtgatga tgttcacaat ttgctgaatt gtggccgggc ccccgtaatg
      901 acctttatac gactgaccca aataaaaaaa gccaccgttg caacttaaga gtcactaacg
      961 gcagcttatg cgaatagtgt tgccacttgc tcaagggaga ccagaaacaa aaaaaggccg
     1021 cgttagcggc cttcaataat tggctcggta ccaaattcca gaaaagaggc ctcccgaaag
     1081 gggggccttt tttcgttttg gtccgaattc gtccctatca gtgatagaga ttgacatccc
     1141 tatcagtgat agagatactg agcactctag agaaagagga gaaaggagat atggagaccc
     1201 gattccctca gcaatcgcag caaactccgg catctaccaa cagacgccgg ccattcaagc
     1261 atgaggatta cccatgtcga agacaacaaa gaagttcaac tctttatgta ttgatcttcc
     1321 tcgcgatctt tctctcgaaa tttaccaatc aattgcttct gtcgctactg gaagcggtga
     1381 tccgcacagt gacgacttta cagcaattgc ttacttaagg taaagagggg aaaggagata
```

```
1441 tggtacgctg gactttgtgg gataccctcg ctttcctcct gttgctcagt ttattgctgc
1501 cgtcattgct gatcatgttc atcccgtcaa cattcaaacg gcctgtctca tcatggaagg
1561 cgctgaattt acggaaaaca ctgttaatgg cgtcgagcgt ccggctgaag ccgctgaatt
1621 gttcgcgttt accttgcgtg tacgcgcagg aaacactgac gttcttactg acgcagaaga
1681 aaacgtgcgt caaaaattac gtgcggaagg agtgatgtaa tgaagatgcc agaaaaacat
1741 gacctgttgg ccgccattct cgcggcaaag gaacaaggca tcggggcaat ccttgcgttt
1801 gcaatggcgt accttcgcgg cagatataat ggcggtgcgt ttacaaaaac agtaatcgac
1861 gcaacgatgt gcgccattat cgcctggttc attcgtgacc ttctcgactt cgccggacta
1921 agtagcaatc tcgcttatat aacgagcgtg tttatcggct acatcggtac tgactcgatt
1981 ggttcgctta tcaaacgctt cgctgctaaa aaagccggag tagaagatgg tagaaatcaa
2041 taatcaacgt aaggcgttcc tcgatatgct ggcgtggtcg gagggaactg ataacggacg
2101 tcagaaaacc agaaatcatg gttatgacgt cattgtaggc ggagagctat ttactgatta
2161 ctccgatcac cctcgcaaac ttgtcacgct aaacccaaaa ctcaaatcaa caggcgccgg
2221 acgctaccag cttctttccc gttggtggga tgcctaccgc aagcagcttg gcctgaaaga
2281 cttctctccg aaaagtcagg acgctgtggc attgcagcag attaaggagc gtggcgcttt
2341 acctatgatt gatcgtggtg atatccgtca ggcaatcgac cgttgcagca atatctgggc
2401 ttcactgccg ggcgctggtt atggtcagtt cgagcataag gctgacagcc tgattgcaaa
2461 attcaaagaa gcgggcggaa cggtcagaga gattgatgta tgagcagagt caccgcgatt
2521 atctccgctc tggttatctg catcatcgtc tgcctgtcat gggctgttaa tcattaccgt
2581 gataacgcca ttacctacaa agcccagcgc gacaaaaatg ccagagaact gaagctggcg
2641 aacgcggcaa ttactgacat gcagatgcgt cagcgtgatg ttgctgcgct cgatgcaaaa
2701 tacacgaagg agttagctga tgctaaagct gaaaatgatg ctctgcgtga tgatgttgcc
2761 gctggtcgtc gtcggttgca catcaaagca gtctgtcagt cagtgcgtga agccaccacc
2821 gcctccggcg tggataatgc agcctccccc cgactggcag acaccgctga acgggattat
2881 ttcaccctca gagagaggct gatcactatg caaaaacaac tggaaggaac ccagaagtat
2941 attaatgagc agtgcagata actagcataa ccccttgggg cctctaaacg ggtcttgagg
3001 ggttttttga gacaaacaaa agaatggaat caaagttaat gctagcagcc gccgcaggca
3061 tgcaagcttg cggccgcgtc gtgactggga aaaccctggc gactagtctt ggactcctgt
3121 tgatagatcc agtaatgacc tcagaactcc atctggattt gttcagaacg ctcggttgcc
3181 gccgggcgtt ttttattggt gagaatccag ggtccccaa taattacgat ttaaatttgt
3241 gtctcaaaat ctctgatgtt acattgcaca agataaaaat atatcatcat gaacaataaa
3301 actgtctgct tacataaaca gtaatacaag gggtgttatg agccatattc agcgtgaaac
3361 gagctgtagc cgtccgcgtc tgaacagcaa catggatgcg gatctgtatg gctataaatg
3421 ggcgcgtgat aacgtgggtc agagcggcgc gaccatttat cgtctgtatg gcaaaccgga
3481 tgcgccggaa ctgtttctga aacatggcaa aggcagcgtg gcgaacgatg tgaccgatga
3541 aatggtgcgt ctgaactggc tgaccgaatt tatgccgctg ccgaccatta aacattttat
3601 tcgcaccccg gatgatgcgt ggctgctgac caccgcgatt ccgggcaaaa ccgcgtttca
3661 ggtgctggaa gaatatccgg atagcggcga aaacattgtg gatgcgctgg ccgtgtttct
3721 gcgtcgtctg catagcattc cggtgtgcaa ctgcccgttt aacagcgatc gtgtgtttcg
3781 tctggcccag gcgcagagcc gtatgaacaa cggcctggtg gatgcgagcg attttgatga
3841 tgaacgtaac ggctggccgg tggaacaggt gtggaaagaa atgcataaac tgctgccgtt
3901 tagcccggat agcgtggtga cccacggcga ttttagcctg gataacctga ttttcgatga
3961 aggcaaactg attggctgca ttgatgtggg ccgtgtgggc attgcggatc gttatcagga
4021 tctggccatt ctgtggaact gcctgggcga atttagcccg agcctgcaaa aacgtctgtt
4081 tcagaaatat ggcattgata atccggatat gaacaaactg caatttcatc tgatgctgga
4141 tgaatttttc taacttggaa gtaagaatgg tgccgaaggc cggactcaaa catcaaaata
4201 agttaatgat aaaaaacaaa taataaaaca caacaatgaa atatgccccc ttttgtgccc
4261 ccactgtttt tctgaccaat ctattttcag cccatcaata aatcggaaag ttaaatcatt
4321 tttaatcagt aagtttggat ccgtagctcg gatccaaacc agtgcatctt ttatccacat
4381 aaaaaatttt ttttcgaaag aactgttcac actgttcacc tttctgtttt ctccttttat
4441 ttcagagtga taggtggtga ataatgggtg aagggtgaac attcgattct tcacctccgg
4501 cattctgccg atgtgactca taccggtgat taatcctccg cactgaaatc actcaggaag
4561 aaaaaagttt tttttgattt gattgttcac actgttcacc tttcgttttt ctcttttaat
4621 ttcagtgtga taacgggtga atatacggtg aagggtgaac agtggattgt tcaccttcgg
4681 gggatatcgg gataaaaaaa gaccggcaga tgccggtcag gtgggtcagg ctgttgtagg
4741 gtcgtcacat tttggcagcc agtcgccgta gctttcctct ttcagcgtca ggttggtctg
4801 tatcccctgt ttggtatggc gtttctcgta attcagtccg tattccttca gcatcaccgg
```

4861 cagccccagc ccgaacattt tcagactgag tacattccgg tagccgtttg cctccatgta
4921 ggccagatag gcgtgataga ggtatttacg gtaattgcgc gggatgatac tggcgttccc
4981 catatacatg ccgctggtct gcggcagggt ttccagatag ccgataaaat caaacgtcgg
5041 gtcggcatcc cgtttgatgt tcagtgcctc gtctgagttc tgctgggact gaagcagtga
5101 ccgggcgagc atcgggtcgc tgaacttctg catcaggtga cgcacgatga ccgccagctc
5161 gcgggtgatt ttgtccttaa gctgcgggtc gcgctcctgc ggggctatct gttccgggaa
5221 gtgaataatc acccgtcggc gtgacacgcc gccgctgcgg tcggtgaagc gcatcgggtt
5281 attgttcacg gccagaatca ccgccgggat gtgcgtggag tacgcatccc ggtatttcgg
5341 gtcaacggac accgcatcgc cgccggtgat ggccttgagt ccggcaccgt cgccgctcca
5401 tttttcctgg tccggcaggc gtatcagtga gaagccagtt aacgcggcac gttcacgcgg
5461 ggattccagc gtctcgatgg tggccgacgt ggcgttatcc tccccggcca gcagggtggc
5521 tatttcggcc atgatacttt tgccgctgcc gccgggaccg gtcacctcca gaaagagctg
5581 ccagtcgtag cggtttgcca gcaccataaa cagtgcagcc agaatcacgt cgcgtttttc
5641 cgcacggcca ccggcggcac ggtcaagcca gcgccagaag gcgggggcgt gggtttccag
5701 cgtttcaccg tccaccggcg gggtgaaatc cacatcgcac agggtgcgca tccagtgtga
5761 cggactgtgc gggtggaacg tgccgttctg cgtgtcgagc acgccgttac gaaagccaat
5821 caggcggcgg gaggggggctt cctgctgcgg aataatcagc ttcagggtat ccaccacgga
5881 ggccaccttc ccggaggaga acggcgcacg cagacgctga aacagcccgg ccacatcccg
5941 ggcaaagtcc tgtggcggca gcaccttcca gacaccattt tcatagcggg acagaagctg
6001 gccgttggca tcgaccgcga gcgcctcgcc gtaatgctca tagatacgca tggccttttc
6061 gctggtactc atggcggaaa actccgcttc gctcatggtg tcgaacgggc tttcagccgg
6121 tggccggatg gcatcgtaaa tggccttacg ggtggcctcc ccgccgtact gcgtgaaggc
6181 atcattccag tcaccgaaga ccggcggcag ggcaacaaca ccttcacacg catctgcggc
6241 tgcggcggct tttttctggc cgtcaccact gaggtcacgg tcagcggcaa ggacaatctg
6301 acaggcgggg tgcttctgcc gggcaaggct ggccagagaa aggaggttca cggaagaaag
6361 cgccaccatc accgtttcac cggtcaggtg atgtacggta agtgcggtcg cgtatccctc
6421 cgctatccac agacgttttc cggcctgatt ctgtccttca agggtgtgac aggtgcccct
6481 gacctgtccg cctttcaggg tgcgcttacg gccgtcagca ctgattaact gaaggttaac
6541 cagttcgccg ctgtcgtcat acagtggcac cacaaggtca ccggcgcgcc agctcacgcc
6601 accggctctg tgtgtgccgg tcagcatccg gcattcccgg ccgggaaagc ccttgcgggt
6661 caggtaggcg ttaccggttc cggtacgggt tttcgccatc agggtttgtg ccagtgcggc
6721 ggcgttcttc cgggcagcgt ctgtttcatc aacggcggcg gtcgtcactg ccgggtcagc
6781 cggtggcagg ctgccggtca cggcagccac ctttgcggcc gcgtcggacg gggaaacacc
6841 aaaaaccttt tcaaccagtt tcaggccgtc accggcacca cactgattgc agtaccaggt
6901 gccgcgcccc tccctgtcat caaaacggaa gcggtcactc ccgccacaga ccggacaggg
6961 ctgatgacgg ttcttcagca cctgaatccc cagcgccggg agaatacgcg gccagtggcc
7021 gagcgcatgg ctgacggtgg cggttacgtt cattttcatg gtgttgttct ccttcagtgc
7081 agtaccggcg cttttatgtg acgggcacag agttcatcca tcacaaccag cccgagaaag
7141 gacagcgacg gcgcggcctt caggggggccg gattccatta aatcttccag cagggcacag
7201 gctatctgac gccctttttc ctcaccgtgc tggcgcagat aaaagccttc cagctcagcg
7261 gcgatggccg cctccagtga ctcaagggtg agatgcgggt agcggtgctg acgttcgcac
7321 acggtcagcc aggcacaggc gacagcgcga cggtaaaggg cagcgcgtaa gacgggcggt
7381 aagggtgttt tcatttgctt ttctccctgt gacagatgac tgcattccgt gccggttgca
7441 ttaactgata aggcatatct gcgtctcctg aagacgtgcg tatccctgcg cgaatacgca
7501 catttaattt ttcgggggtc gtttttttaat tacagataat tgcggtaact gttatccggg
7561 gtggtttccg ggtcaggctc cgtgcgggga atttcccgcc attcccgcgc caccggtgct
7621 gcccggctga ccggaacagt gtcctgcggg taaatatcca gatatttttc ccgccatttc
7681 tgtaattccg ggtctccggc catttctttc agtaccgcat gccggtttac ggggctgcgt
7741 ttaaacaggt caggacggtc acaggtaaat tcccgcagaa aacgccccag cgggatgtct
7801 gtggtgcgtc cgtcagcgag gatacgcaca aggatactga atttacggcg gtacgggttc
7861 cagacaatgt ccgggcagcg gtacggcatt tcccacggaa taccgtcttc cagaatgccg
7921 accacggcca catcgggaaa accggcagaa cggtaaatct caccgggctg gggaaaatca
7981 aacatgcgtc ctgtctcccc ggtctttctg ctgggcgaga aaatcgcggc acaggccttt
8041 ggctttcagc tcattcagca caaaatcaaa ggggcttttt tattacgcac gggacacctc
8101 caccaccggc agacgggcag caagggagag cacatagtca cggacaaggg aacggcgggc
8161 actgcgttca tcaccggcga cggtgcgaag catacagata cggggatgac ggtctgcgcg
8221 acggacagcc gcaaacacaa agacaaattc agggtgtgag gggtaaggg ttgtagccat

```
     8281 gatggcagcc tcctgtgaat agcaaataac gctatcgccg gagttctcac gctcgatggc
     8341 gatagcccag acgggggtga gaataccggc ttcacaggat accggccagc ccggaggctg
     8401 ccccgcctga gctaccattg actctgcggc ataatgagcg gacgcgggca ggatgcacgg
     8461 aatgccatct gcacgactga ccacacacca caccataatc tggcgctctg tggcattgat
     8521 tgcgacacaa aaaaagacgc gtggcgcgtc atatgtcgcc tgtgaattgc tcgggttctc
     8581 acgcccggct gccgattttg cggcaggcga aaaactatat ccgcaaatgc cggaaaaagg
     8641 caagccagaa aaagggagtt tttgcagagc gggcatcatc atgcgtcgta cccccgtttg
     8701 cgtccggcaa tgcgtccggc catccatgcg gtgacttcag agtgcagcca ggccacattt
     8761 ttaccgccaa gactcacctg cggcggaaat tccccccttac ggatgagttc gtagatggtc
     8821 gagcgtgaca ggccgcacag gtgcatcact tccggcacac gtaaaaaacg ctcctgcgtg
     8881 atgtccggca gcggcatcag tggcgtcact ggggcgggag acggggaaga aaaaacagct
     8941 tgcatcgggc tacctcgtta atgtccatac agcaccggat aagtccgtcc ggcttcgggt
     9001 agcgctttat tttgtgaata ttttcagcag acgcaacagg ggggatttgt tcaggctgtc
     9061 ttacaatggc tgtgtgtttt ttgttcatct ccacttaaag tcatttaaag ccacttaaag
     9121 caatttgtaa tttttatagt gaaatacaaa tcgtttcttc ttattcattc ccggcgaatt
     9181 aataaaaaca aacagtagta aacagcacaa aaagcccatc aacgggtgaa cagtggtgaa
     9241 cagacggtga acagtcatta ctgcgattgt tcacccttta acttactgta ttacttatct
     9301 tttttattaa ggtgaacaga ggtgaacagt aaaatataaa aaaacaaaca gtaagccggt
     9361 ttttcctgcg accttttcct ggcttgccgg tctgaggatg agtctcctgt gtcagggctg
     9421 gcacatctgc aatgcgtcgt gttgttgtcc ggtgtacgtc acaattttct taacctgaag
     9481 tgacgaggag ccggaaaatg tctgaccaca ctatccctga atatctgcaa cccgcactgg
     9541 cacaactgga aaaggccaga gccgcccatc ttgagaacgc ccgcctgatg gatgagaccg
     9601 tcacggccat tgaacgggca gagcaggaaa aaaatgcgct ggcgcaggcc gacggaaacg
     9661 acgctgacga ctggcgcacg gcctttcgtg cagccggtgg tgtcctgagc gacgagctga
     9721 aacagcgcca cattgagcgc gtggcacgcc gggagctggt acaggaatat gacaatctgg
     9781 ccgtggtgct gaatttcgaa cgtgaacgcc tgaaaggggc gtgtgacagc acggccaccg
     9841 cctaccggaa ggcacatcat caccttctga gtctgtatgc agagcatgag ctggaacacg
     9901 ccctgaatga aacctgtgag gcgcttgtcc gggcaatgca tctgagcatt ctggtacagg
     9961 aaaatccgct cgccaacacc accggccatc agggctacgt cgcaccggaa aaggctgtca
    10021 tgcagcaggt gaaatcatcg ctggaacaga aaattaaaca gatgcaaatc agcctcaccg
    10081 gcgagccggt tctccggctg accggactgt cagcggcaac actcccgcac atggattatg
    10141 aggtggcagg cacaccggca cagcgcaagg tgtggcagga caaaatagac cagcagggag
    10201 cagagcttaa ggccagaggg ctgctgtcat gatttactgt ccgtcgtgtg gacatgttgc
    10261 tcacacccgt cgcgcacatt tcatggacga tggcaccaag ataatgattg cacagtgccg
    10321 gaatatttat tgctctgcga catttgaagc gagtgaaagc tttttctctg acagtaaaga
    10381 ttcaggaatg gaatacattt caggcaaaca gagataccgc gattcactga cgtcagcctc
    10441 ctgcggtatg aaacgcccga aaagaatgct tgttaccgga tattgttgtc ggagatgtaa
    10501 aggccttgca ctgtcaagaa catcgcggcg tctgtctcag gaagtcaccg agcgttttta
    10561 tgtgtgcacg gatccgggct gtggtctggt gtttaaaacg cttcagacca tcaaccgctt
    10621 cattgtccgc ccggtcacgc cggacgaact ggcagaacgc ctgcatgaaa aacaggaact
    10681 gccgccagta cggttaaaaa cacaatcata ttcgctgcgt ctggaatgag ggctgccggt
    10741 taacaccggc cgtcgccgca caccgtattt ttattcttca gcatgatgag aaagagataa
    10801 cgatggaaag cacagcctta cagcaggcct ttgacacctg tcagaataac aaagcagcat
    10861 ggctgcaacg caaaaatgag ctggcagcgg ccgaacagga atatctgcgg cttctgtcag
    10921 gagaaggcag aaacgtcagt cgcctggacg aattacgcaa tattatcgaa gtcagaaaat
    10981 ggcaggtgaa tcaggccgcc ggtcgttata ttcgttcgca tgaagccgtt cagcacatca
    11041 gcatccgcga ccggctgaat gattttatgc agcagcacgg cacagcactg gcggccgcac
    11101 tggcaccgga gctgatgggc tacagtgagc tgacggccat tgcccgaaac tgtgccatac
    11161 agcgtgccac agatgccctg cgtgaagccc ttctgtcctg gcttgcgaag ggtgaaaaaa
    11221 ttaattattc cgcacaggat agcgacattt taacgaccat cggattcagg cctgacgtgg
    11281 cttcggtgga tgacagccgt gaaaaattca cccctgcgca gaacatgatt ttttcgcgta
    11341 aaagtgcgca actggcatca cgtcagtcag tgtaaaattc cccgaaaatc cgcccgtttt
    11401 tactgaaaaa agccatgcat cgataaggtg catggcttt
//
```

#### SS8.6. Whole-cosmid sequencing result of the stable version of P4-EKORhE with the multi-lysins cassette.

```
LOCUS       Exported File          11432 bp ds-DNA    circular SYN 29-JUL-2024
```

```
DEFINITION  .
ACCESSION   .
VERSION     .
KEYWORDS    Ramirez_Garcia_y9v_1_P4-EKORhE-A_pLann_11kb_featured_REVseq
SOURCE      synthetic DNA construct
  ORGANISM  synthetic DNA construct
REFERENCE   1  (bases 1 to 11432)
  AUTHORS   Robert Ramirez-Garcia
  TITLE     Direct Submission
  JOURNAL   Exported 29 Jul 2024
  COMMENT     Designed by Robert Ramirez-Garcia on 26-8-2021
     Whole-cosmid sequenced using Plasmidsaurus service
   with Oxford Nanopore Technologies https://www.plasmidsaurus.com/
FEATURES             Location/Qualifiers
     source          1..11432
                     /organism="synthetic DNA construct"
                     /mol_type="other DNA"
     source          2948..2995
                     /note="color: #ffffff"
     misc_feature    2..126
                     /note="P4 packaging region"
                     /note="color: #00ff00"
     misc_feature    3..307
                     /note="P4 packaging region"
                     /note="color: #00ff00"
     misc_feature    3..232
                     /note="P4 packaging region"
                     /note="color: #00ff00"
     misc_feature    101..119
                     /note="P2 cos site"
                     /note="color: #a6acb3"
     promoter        233..279
                     /note="P4-Pgop promoter"
                     /note="This forward directional feature has 6 segments:
                     1:233..241/#993300
                     2:242..247/#993300/-35
                     3:248..263/#993300
                     4:264..269/#993300/-10
                     5:270..278/#993300
                     6:279..279/#993300/+1"
     terminator      351..445
                     /gene="
                     "
                     /note="lambda t0 terminator"
                     /note="transcription terminator from phage lambda"
                     /note="color: #ffffff"
     terminator      401..435
                     /label="lambda t0 terminator"
                     /note="lambda t0 terminator pLannotate"
                     /note="pLannotate"
                     /note="color: #ffffff; direction: RIGHT"
     misc_feature    450..469
                     /note="v3_Epsilon_INS"
                     /note="color: #a6acb3"
     CDS             complement(453..737)
                     /label="Epsilon"
                     /note="P4-gp9 epsilon"
```

```
                     /note="pLannotate"
                     /note="color: #993366; direction: LEFT"
     RBS             complement(738..762)
                     /note="BBa_B0064 weak RBS"
                     /note="color: #ff0000; direction: LEFT"
     RBS             complement(738..761)
                     /note="BBa_B0064 weak RBS"
                     /note="color: #ff0000; direction: LEFT"
     RBS             complement(744..755)
                     /note="RBS-64"
                     /note="color: #993300; direction: LEFT"
     promoter        complement(769..887)
                     /note="rhaB promoter"
                     /note="This reverse directional feature has 8 segments:
                     1:769..769/#800000/+1
                     2:770..800/#800000
                     3:801..817/#ff9900/RhaSop
                     4:818..833/#800000
                     5:834..850/#ff9900/RhaSop
                     6:851..853/#800000
                     7:854..869/#ff9900/CRPop
                     8:870..887/#800000"
     promoter        complement(769..887)
                     /note="rhaB promoter"
                     /note="This reverse directional feature has 7 segments:
                     1:769..800/#800000
                     2:801..817/#ff9900/RhaSop
                     3:818..833/#800000
                     4:834..850/#ff9900/RhaSop
                     5:851..853/#800000
                     6:854..869/#ff9900/CRPop
                     7:870..887/#800000"
     promoter        770..885
                     /label="rhaB promoter"
                     /note="rhaB promoter pLannotate"
                     /note="pLannotate"
                     /note="color: #ffffff"
     protein_bind    complement(801..817)
                     /note="RhaS binding site"
                     /note="color: #31849b; direction: LEFT"
     protein_bind    complement(834..850)
                     /note="RhaS binding site"
                     /note="color: #31849b; direction: LEFT"
     misc_feature    836..884
                     /note="Seq BQK537 Part 2 - wt-Delta-Epsilon num1 colony-A"
                     /note="color: #000000"
     protein_bind    complement(854..869)
                     /note="CRP binding site"
                     /note="color: #31849b; direction: LEFT"
     misc_feature    886..911
                     /note="Seq BQK537 Part 3 - wt-Delta-Epsilon num1 colony-A"
                     /note="color: #000000"
     terminator      complement(894..995)
                     /note="TNA21 terminator"
                     /note="color: #ff0000; direction: LEFT"
     CDS             complement(1008..1019)
                     /codon_start=1
```

```
                     /product="Factor Xa recognition and cleavage site"
                     /note="Factor Xa site"
                     /note="color: #cc99b2
                     Cleavage site after base 1007"
                     /translation="IEGR"
     promoter        1072..1145
                     /note="pLtet vII weak promoter 0.05x PLtet full"
                     /note="This forward directional feature has 6 segments:
                     1:1072..1091/#800000
                     2:1092..1092/#800000/mutationT->G
                     3:1093..1114/#800000
                     4:1115..1120/#800000/-10
                     5:1121..1125/#800000
                     6:1126..1145/#800000/Lutzlinkerfrom+1"
     promoter        1072..1145
                     /note="PLtetO-1 promoter"
                     /note="This forward directional feature has 6 segments:
                     1:1072..1090/#993300
                     2:1091..1096/#000000/-35
                     3:1097..1113/#993300
                     4:1114..1119/#000000/-10
                     5:1120..1125/#993300
                     6:1126..1145/#993300/Lutzlinkerafterpromoter"
     promoter        1072..1145
                     /note="PLtetO-1 promoter full weak 0.14x"
                     /note="This forward directional feature has 8 segments:
                     1:1072..1089/#993300
                     2:1090..1090/#993300
                     3:1091..1095/#000000/-35withmutationG->T
                     4:1096..1096/#000000/mutationA->C
                     5:1097..1113/#993300
                     6:1114..1119/#000000/-10
                     7:1120..1125/#993300
                     8:1126..1145/#993300/Lutzlinkerafterpromoter"
     protein_bind    1072..1090
                     /label="tet operator"
                     /note="tet operator"
                     /note="pLannotate"
                     /note="color: #31849b; direction: RIGHT"
     promoter        1096..1115
                     /label="tight TRE promoter (fragment)"
                     /note="tight TRE promoter"
                     /note="pLannotate"
                     /note="color: #ffffff; direction: RIGHT"
     protein_bind    1097..1115
                     /gene="tetO"
                     /bound_moiety="tetracycline repressor TetR"
                     /note="tet operator"
                     /note="
                     "
                     /note="color: #31849b"
     misc_feature    1098..1125
                     /note="tetO1-multi-LysKOrv"
                     /note="color: #a6acb3"
     RBS             1152..1170
                     /note="RBS-34"
                     /note="This forward directional feature has 2 segments:
```

```
                    1:1152..1158/#993300/biobrickscar1
                    2:1159..1170/#993300/RBS"
RBS                 1159..1170
                    /note="color: #a6acb3"
RBS                 1159..1170
                    /note="RBS34min"
                    /note="color: #ff6600"
CDS                 1177..1404
                    /codon_start=1
                    /note="MS2 gpL lysis protein"
                    /note="color: #00ccff"
                    /translation="METRFPQQSQQTPASTNRRRPFKHEDYPCRRQQRSSTLYVLIFLA
                    IFLSKFTNQLLLSLLEAVIRTVTTLQQLLT"
RBS                 1408..1419
                    /note="B0064"
                    /note="color: #666699"
RBS                 1408..1419
                    /note="RBS-64"
                    /note="color: #993300; direction: RIGHT"
CDS                 1426..1701
                    /codon_start=1
                    /note="PhiX174 gpE lysis protein"
                    /note="color: #00ccff"
                    /translation="MVRWTLWDTLAFLLLLSLLLPSLLIMFIPSTFKRPVSSWKALNLR
                    KTLLMASSVRLKPLNCSRLPCVYAQETLTFLLTQKKTCVKNYVRKE"
CDS                 1706..2029
                    /codon_start=1
                    /note="Lambda lysS"
                    /note="color: #00ccff"
                    /translation="MKMPEKHDLLAAILAAKEQGIGAILAFAMAYLRGRYNGGAFTKTV
                    IDATMCAIIAWFIRDLLDFAGLSSNLAYITSVFIGYIGTDSIGSLIKRFAAKKAGVEDG
                    RNQ"
CDS                 2013..2489
                    /codon_start=1
                    /note="Lambda lysR"
                    /note="color: #00ccff"
                    /translation="MVEINNQRKAFLDMLAWSEGTDNGRQKTRNHGYDVIVGGELFTDY
                    SDHPRKLVTLNPKLKSTGAGRYQLLSRWWDAYRKQLGLKDFSPKSQDAVALQQIKERGA
                    LPMIDRGDIRQAIDRCSNIWASLPGAGYGQFEHKADSLIAKFKEAGGTVREIDV"
CDS                 2486..2947
                    /codon_start=1
                    /note="Lambda Rz"
                    /note="color: #00ccff"
                    /translation="MSRVTAIISALVICIIVCLSWAVNHYRDNAITYKAQRDKNARELK
                    LANAAITDMQMRQRDVAALDAKYTKELADAKAENDALRDDVAAGRRRLHIKAVCQSVRE
                    ATTASGVDNAASPRLADTAERDYFTLRERLITMQKQLEGTQKYINEQCR"
CDS                 2486..2946
                    /codon_start=1
                    /note="Lambda Rz"
                    /note="color: #00ccff"
                    /translation="MSRVTAIISALVICIIVCLSWAVNHYRDNAITYKAQRDKNARELK
                    LANAAITDMQMRQRDVAALDAKYTKELADAKAENDALRDDVAAGRRRLHIKAVCQSVRE
                    ATTASGVDNAASPRLADTAERDYFTLRERLITMQKQLEGTQKYINEQCR"
CDS                 2486..2944
                    /label="Rz"
                    /note="Rz"
                    /note="pLannotate"
```

```
                     /note="color: #993366; direction: RIGHT"
terminator           2948..2995
                     /label="T7 terminator"
                     /note="T7 terminator pLannotate"
                     /note="pLannotate"
                     /note="color: #ffffff; direction: RIGHT"
CDS                  2996..3025
                     /label="GFP (fragment)"
                     /note="GFP"
                     /note="pLannotate"
                     /note="color: #993366; direction: RIGHT"
terminator           2996..3025
                     /note="T500 Terminator"
                     /note="color: #993300; direction: RIGHT"
misc_feature         3026..3040
                     /note="BioBrick suffix"
                     /note="This feature has 3 segments:
                     1:3026..3035/#a6ccff/ChangedSpeItoNheI(com...
                     2:3036..3036/#a6ccff/obliterationofEagI/NotI
                     3:3037..3040/#a6ccff"
misc_feature         3041..3058
                     /label="MCS (fragment)"
                     /note="MCS"
                     /note="pLannotate"
                     /note="color: #a6acb3; direction: RIGHT"
CDS                  3066..3090
                     /label="lacZ-alpha (fragment)"
                     /note="lacZ-alpha"
                     /note="pLannotate"
                     /note="color: #993366; direction: RIGHT"
primer_bind          3067..3090
                     /note="F24"
                     /note="color: #a020f0; direction: RIGHT"
misc_feature         3097..3199
                     /note="T0"
                     /note="color: #a6acb3"
terminator           complement(3097..3199)
                     /note="T0 terminator"
                     /note="color: #ffffff; direction: LEFT"
terminator           3101..3195
                     /label="lambda t0 terminator"
                     /note="lambda t0 terminator pLannotate"
                     /note="pLannotate"
                     /note="color: #ffffff; direction: RIGHT"
terminator           3151..3185
                     /note="lambda t0r terminator"
                     /note="color: #800080; direction: RIGHT"
promoter             3228..3326
                     /note="KanR-aph(3')-Ia promoter"
                     /note="color: #808000; direction: RIGHT"
CDS                  3327..4142
                     /note="Km"
                     /note="color: #993366; direction: RIGHT"
CDS                  3327..4139
                     /label="aphA1"
                     /note="aphA1"
                     /note="pLannotate"
```

```
                     /note="color: #993366; direction: RIGHT"
     CDS             complement(4175..4510)
                     /codon_start=1
                     /transl_table=11
                     /locus_tag="P4p05"
                     /product="hypothetical protein"
                     /note="P4p05"
                     /note="Predicted by GeneMark"
                     /note="color: #993366"
                     /db_xref="GeneID:1261089"
                     /protein_id="NP_597796.1"
                     /translation="MSHIGRMPEVKNRMFTLHPLFTTYHSEIKGENRKVNSVNSSFEKK
                     FFMWIKDALVWIRATDPNLLIKNDLTFRFIDGLKIDWSEKQWGHKRGHISLLCFIICFL
                     SLTYFDV"
     misc_feature    4370..4669
                     /note="crr"
                     /note="crr (cis required region for replication)"
                     /note="color: #a6acb3"
     repeat_region   4370..4489
                     /note="crr 120bp direct repeat"
                     /note="color: #a6acb3"
     repeat_region   4550..4669
                     /note="crr 120bp direct repeat"
                     /note="color: #a6acb3"
     CDS             complement(4715..7048)
                     /codon_start=1
                     /note="P4p06"
                     /note="color: #993366"
                     /translation="MKMNVTATVSHALGHWPRILPALGIQVLKNRHQPCPVCGGSDRFR
                     FDDREGRGTWYCNQCGAGDGLKLVEKVFGVSPSDAAAKVAAVTGSLPPADPAVTTAAVD
                     ETDAARKNAAALAQTLMAKTRTGTGNAYLTRKGFPGRECRMLTGTHRAGGVSWRAGDLV
                     VPLYDDSGELVNLQLISADGRKRTLKGGQVRGTCHTLEGQNQAGKRLWIAEGYATALTV
                     HHLTGETVMVALSSVNLLSLASLARQKHPACQIVLAADRDLSGDGQKKAAAAADACEGV
                     VALPPVFGDWNDAFTQYGGEATRKAIYDAIRPPAESPFDTMSEAEFSAMSTSEKAMRIY
                     EHYGEALAVDANGQLLSRYENGVWKVLPPQDFARDVAGLFQRLRAPFSSGKVASVVDTL
                     KLIIPQQEAPSRRLIGFRNGVLDTQNGTFHPHSPSHWMRTLCDVDFTPPVDGETLETHA
                     PAFWRWLDRAAGGRAEKRDVILAALFMVLANRYDWQLFLEVTGPGGSGKSIMAEIATLL
                     AGEDNATSATIETLESPRERAALTGFSLIRLPDQEKWSGDGAGLKAITGGDAVSVDPKY
                     RDAYSTHIPAVILAVNNNPMRFTDRSGGVSRRRVIIHFPEQIAPQERDPQLKDKITREL
                     AVIVRHLMQKFSDPMLARSLLQSQQNSDEALNIKRDADPTFDFIGYLETLPQTSGMYMG
                     NASIIPRNYRKYLYHAYLAYMEANGYRNVLSLKMFGLGLPVMLKEYGLNYEKRHTKQGI
                     QTNLTLKEESYGDWLPKCDDPTTA"
     CDS             complement(7063..7383)
                     /codon_start=1
                     /transl_table=11
                     /locus_tag="P4p07"
                     /product="hypothetical protein"
                     /note="P4p07"
                     /note="ORF106 (AA 1-106)"
                     /note="color: #993366"
                     /db_xref="UniProtKB/Swiss-Prot:P10278"
                     /db_xref="GeneID:1261093"
                     /protein_id="NP_042037.1"
                     /translation="MKTPLPPVLRAALYRRAVACAWLTVCERQHRYPHLTLESLEAAIA
                     AELEGFYLRQHGEEKGRQIACALLEDLMESGPLKAAPSLSFLGLVVMDELCARHIKAPV
                     LH"
     CDS             complement(7519..7974)
```

```
                     /codon_start=1
                     /transl_table=11
                     /locus_tag="P4p08"
                     /product="hypothetical protein"
                     /note="P4p08"
                     /note="ORF151 (AA 1-151)"
                     /note="color: #993366"
                     /db_xref="UniProtKB/Swiss-Prot:P05464"
                     /db_xref="GeneID:1261086"
                     /protein_id="NP_042038.1"
                     /translation="MFDFPQPGEIYRSAGFPDVAVVGILEDGIPWEMPYRCPDIVWNPY
                     RRKFSILVRILADGRTTDIPLGRFLREFTCDRPDLFKRSPVNRHAVLKEMAGDPELQKW
                     REKYLDIYPQDTVPVSRAAPVAREWREIPRTEPDPETTPDNSYRNYL"
     misc_feature    7863..7871
                     /note="GSG linker"
                     /note="color: #a6acb3"
     misc_feature    8036..8058
                     /note="DISPENSIBLE"
                     /note="color: #000000"
     CDS             complement(8076..8492)
                     /codon_start=1
                     /transl_table=11
                     /locus_tag="P4p10"
                     /product="putative CI repressor"
                     /note="P4p10"
                     /note="cI gene product (AA 1-137)"
                     /note="color: #993366"
                     /db_xref="UniProtKB/Swiss-Prot:P05462"
                     /db_xref="GeneID:1261091"
                     /protein_id="NP_042040.1"
                     /translation="MMVWCVVSRADGIPCILPASAHYAAESMVAQAGQPPGWPVSCEAG
                     ILTPVWAIAIERENSGDSVICYSQEAAIMATTLTPSHPEFVFVFAAVRRADRHPRICML
                     RTVAGDERSARRSLVRDYVLSLAARLPVVEVSRA"
     CDS             complement(8082..8492)
                     /label="cI"
                     /note="cI"
                     /note="pLannotate"
                     /note="color: #993366; direction: LEFT"
     ncRNA           complement(8562..8614)
                     /label="c4-a1b1"
                     /note="c4-a1b1"
                     /note="pLannotate"
                     /note="color: #8fbc8f; direction: LEFT"
     promoter        complement(8995..9141)
                     /note="P4-pLL promoter"
                     /note="This reverse directional feature has 8 segments:
                     1:8995..8995/#800000/+1
                     2:8996..9001/#800000
                     3:9002..9007/#800000/-10
                     4:9008..9022/#800000
                     5:9023..9028/#800000/-35
                     6:9029..9033/#800000
                     7:9034..9063/#ff6600/Ogr/deltaop
                     8:9064..9141/#800000/Coxbindingoperators"
     repeat_region   9084..9093
                     /note="ori type 2 repeat"
                     /note="color: #a6acb3"
```

```
repeat_region   9094..9103
                /note="ori type 2 repeat"
                /note="color: #a6acb3"
repeat_region   9104..9113
                /note="ori type 2 repeat"
                /note="color: #a6acb3"
repeat_region   9104..9113
                /note="ori type 2 repeat"
                /note="color: #a6acb3"
promoter        9395..9468
                /note="P4-sid promoter"
                /note="This forward directional feature has 11 segments:
                1:9395..9400/#800000
                2:9401..9418/#ff9900/op1
                3:9419..9429/#800000
                4:9430..9435/#800000/-35
                5:9436..9438/#800000
                6:9439..9452/#ff9900/op2
                7:9453..9456/#ff9900/-10
                8:9457..9458/#800000/-10
                9:9459..9464/#800000
                10:9465..9465/#800000/+1
                11:9466..9468/#800000"
CDS             9491..10222
                /label="sid"
                /note="P4-gp12 sid"
                /note="pLannotate"
                /note="color: #993366; direction: RIGHT"
gene            10222..10722
                /locus_tag="P4p13"
                /note="P4p13"
                /note="color: #a6acb3; direction: RIGHT"
                /db_xref="GeneID:1261088"
CDS             10222..10722
                /label="Delta"
                /note="P4-gp13 delta"
                /note="pLannotate"
                /note="color: #993366; direction: RIGHT"
gene            10796..11368
                /locus_tag="P4p14"
                /note="P4p14"
                /note="color: #a6acb3; direction: RIGHT"
                /db_xref="GeneID:1261094"
CDS             10796..11368
                /codon_start=1
                /transl_table=11
                /locus_tag="P4p14"
                /product="amber mutation-suppressing protein"
                /note="P4p14"
                /note="psu gene product (AA 1-190)"
                /note="color: #993366"
                /db_xref="UniProtKB/Swiss-Prot:P05460"
                /db_xref="GeneID:1261094"
                /protein_id="NP_042044.1"
                /translation="MESTALQQAFDTCQNNKAAWLQRKNELAAAEQEYLRLLSGEGRNV
                SRLDELRNIIEVRKWQVNQAAGRYIRSHEAVQHISIRDRLNDFMQQHGTALAAALAPEL
                MGYSELTAIARNCAIQRATDALREALLSWLAKGEKINYSAQDSDILTTIGFRPDVASVD
```

```
                     DSREKFTPAQNMIFSRKSAQLASRQSV"
     CDS             10799..11368
                     /codon_start=1
                     /note="P4-gp14 psu"
                     /note="color: #993366"
                     /translation="ESTALQQAFDTCQNNKAAWLQRKNELAAAEQEYLRLLSGEGRNVS
                     RLDELRNIIEVRKWQVNQAAGRYIRSHEAVQHISIRDRLNDFMQQHGTALAAALAPELM
                     GYSELTAIARNCAIQRATDALREALLSWLAKGEKINYSAQDSDILTTIGFRPDVASVDD
                     SREKFTPAQNMIFSRKSAQLASRQSV"
     terminator      11369..11432
                     /note="tsid terminator"
                     /note="color: #000080; direction: RIGHT"
ORIGIN
        1 gcatgcgttt tcctgcctca ttttctgcaa accgcgccat tcccggcgcg gtctgagcgt
       61 gtcagtgcaa ctgcattaaa accgccccgc aaagcgggcg ggcgaggcgg ggaaagcacc
      121 gcgcgcaaac cgacaagtta gttaattatt tgtgtagtca aagtgccttc agtacatacc
      181 tcgttaatac attggagcat aatgaagaaa atctatggcc tatggtccaa aactgtcttt
      241 tttgatggca ctatcctgaa aaatatgcaa aaaatagatt gatgtaaggt ggttcttgtc
      301 agtgtcgcaa gatccttaag aattcgtggc atgagagagt taaaggcttg gactcctgtt
      361 gatagatcca ataatgacct cagaactcca tctggatttg ttcagaacgc tcggttgccg
      421 ccgggcgttt tttattggtg agaatccagt taaacgtggg tgcgatcgcc gcggtctttc
      481 tgctgcgcca ggaaatcacg gcacaggcct ttcgctttca gctcattcag cacgaagtcg
      541 atgtcttcgt tcaggtagga gaagatgtgc gggatataca gctggtgcag accgctcagt
      601 tcacggtgga tcaggtcgcc gccaacgaag cggctgacgg cgctgacacg gttcagtttg
      661 tgatacgctt cgatggacag gtgttcgcgc gcgtcgtgac gcgcggaaac ggtctgcggt
      721 gctttttgt tacgcatcta gtatttcccc tctttctcta gagatctcca cgaccagtct
      781 aaaaagcgcc tgaattcgcg accttctcgt tactgacagg aaaatgggcc attggcaacc
      841 agggaaagat gaacgtgatg atgttcacaa tttgctgaat tgtggccggg cccccgtaat
      901 gacctttata cgactgaccc aaataaaaaa agccaccgtt gcaacttaag agtcactaac
      961 ggcagcttat gcgaatagtg ttgccacttg ctcaagggag accgttagcg gccttcaata
     1021 attggctata aaaataggcg tatcacgagg ccctttcgtc ttcacctcga gtccctatca
     1081 gtgatagaga ttgacctccc tatcagtgat agagatactg agcacatcag caggacgcac
     1141 tgaccccatg gtctagagaa agaggagaaa ggagatatgg agacccgatt ccctcagcaa
     1201 tcgcagcaaa ctccggcatc taccaacaga cgccggccat tcaagcatga ggattaccca
     1261 tgtcgaagac aacaaagaag ttcaactctt tatgtattga tcttcctcgc gatctttctc
     1321 tcgaaattta ccaatcaatt gcttctgtcg ctactggaag cggtgatccg cacagtgacg
     1381 actttacagc aattgcttac ttaaggtaaa gaggggaaag gagatatggt acgctggact
     1441 ttgtgggata ccctcgcttt cctcctgttg ctcagtttat tgctgccgtc attgctgatc
     1501 atgttcatcc cgtcaacatt caaacggcct gtctcatcat ggaaggcgct gaatttacgg
     1561 aaaacactgt taatggcgtc gagcgtccgg ctgaagccgc tgaattgttc gcgtttacct
     1621 tgcgtgtacg cgcaggaaac actgacgttc ttactgacgc agaagaaaac gtgcgtcaaa
     1681 aattacgtgc ggaaggagtg atgtaatgaa gatgccagaa aaacatgacc tgttggccgc
     1741 cattctcgcg gcaaaggaac aaggcatcgg ggcaatcctt gcgtttgcaa tggcgtacct
     1801 tcgcggcaga tataatggcg gtgcgtttac aaaaacagta atcgacgcaa cgatgtgcgc
     1861 cattatcgcc tggttcattc gtgaccttct cgacttcgcc ggactaagta gcaatctcgc
     1921 ttatataacg agcgtgttta tcggctacat cggtactgac tcgattggtt cgcttatcaa
     1981 acgcttcgct gctaaaaaag ccggagtaga agatggtaga aatcaataat caacgtaagg
     2041 cgttcctcga tatgctggcg tggtcggagg gaactgataa cggacgtcag aaaaccagaa
     2101 atcatggtta tgacgtcatt gtaggcggag agctatttac tgattactcc gatcaccctc
     2161 gcaaacttgt cacgctaaac ccaaaactca aatcaacagg cgccggacgc taccagcttc
     2221 tttcccgttg gtgggatgcc taccgcaagc agcttggcct gaaagacttc tctccgaaaa
     2281 gtcaggacgc tgtggcattg cagcagatta aggagcgtgg cgctttacct atgattgatc
     2341 gtggtgatat ccgtcaggca atcgaccgtt gcagcaatat ctgggcttca ctgccgggcg
     2401 ctggttatgg tcagttcgag cataaggctg acagcctgat tgcaaaattc aaagaagcgg
     2461 gcggaacggt cagagagatt gatgtatgag cagagtcacc gcgattatct ccgctctggt
     2521 tatctgcatc atcgtctgcc tgtcatgggc tgttaatcat taccgtgata acgccattac
     2581 ctacaaagcc cagcgcgaca aaaatgccag agaactgaag ctggcgaacg cggcaattac
```

```
2641 tgacatgcag atgcgtcagc gtgatgttgc tgcgctcgat gcaaaataca cgaaggagtt
2701 agctgatgct aaagctgaaa atgatgctct gcgtgatgat gttgccgctg gtcgtcgtcg
2761 gttgcacatc aaagcagtct gtcagtcagt gcgtgaagcc accaccgcct ccggcgtgga
2821 taatgcagcc tccccccgac tggcagacac cgctgaacgg gattatttca ccctcagaga
2881 gaggctgatc actatgcaaa aacaactgga aggaacccag aagtatatta atgagcagtg
2941 cagataacta gcataacccc ttggggcctc taaacgggtc ttgaggggtt ttttgagaca
3001 aacaaaagaa tggaatcaaa gttaatgcta gcagccgccg ctgcaggcat gcaagcttgc
3061 ggccgcgtcg tgactgggaa aaccctggcg actagtcttg gactcctgtt gatagatcca
3121 gtaatgacct cagaactcca tctggatttg ttcagaacgc tcggttgccg ccgggcgttt
3181 tttattggtg agaatccagg ggtccccaat aattacgatt taaatttgtg tctcaaaatc
3241 tctgatgtta cattgcacaa gataaaaata tatcatcatg aacaataaaa ctgtctgctt
3301 acataaacag taatacaagg ggtgttatga gccatattca gcgtgaaacg agctgtagcc
3361 gtccgcgtct gaacagcaac atggatgcgg atctgtatgg ctataaatgg gcgcgtgata
3421 acgtgggtca gagcggcgcg accatttatc gtctgtatgg caaaccggat gcgccggaac
3481 tgtttctgaa acatggcaaa ggcagcgtgg cgaacgatgt gaccgatgaa atggtgcgtc
3541 tgaactggct gaccgaattt atgccgctgc cgaccattaa acattttatt cgcacccogg
3601 atgatgcgtg gctgctgacc accgcgattc cgggcaaaac cgcgtttcag gtgctggaag
3661 aatatccgga tagcggcgaa aacattgtgg atgcgctggc cgtgtttctg cgtcgtctgc
3721 atagcattcc ggtgtgcaac tgcccgttta acagcgatcg tgtgtttcgt ctggcccagg
3781 cgcagagccg tatgaacaac ggcctggtgg atgcgagcga ttttgatgat gaacgtaacg
3841 gctggccggt ggaacaggtg tggaaagaaa tgcataaact gctgccgttt agcccggata
3901 gcgtggtgac ccacggcgat tttagcctgg ataacctgat tttcgatgaa ggcaaactga
3961 ttggctgcat tgatgtgggc cgtgtgggca ttgcggatcg ttatcaggat ctggccattc
4021 tgtggaactg cctgggcgaa tttagcccga gcctgcaaaa acgtctgttt cagaaatatg
4081 gcattgataa tccggatatg aacaaactgc aatttcatct gatgctggat gaatttttct
4141 aacttggaag taagaatggt gccgaaggcc ggactcaaac atcaaaataa gttaatgata
4201 aaaaacaaat aataaaacac aacaatgaaa tatgccccct tttgtgcccc cactgttttt
4261 ctgaccaatc tattttcagc ccatcaataa atcggaaagt taaatcattt ttaatcagta
4321 agtttggatc cgtagctcgg atccaaacca gtgcatcttt tatccacata aaaaattttt
4381 tttcgaaaga actgttcaca ctgttcacct ttctgttttc tccttttatt tcagagtgat
4441 aggtggtgaa taatgggtga agggtgaaca ttcgattctt cacctccggc attctgccga
4501 tgtgactcat accggtgatt aatcctccgc actgaaatca ctcaggaaga aaaaagtttt
4561 ttttgatttg attgttcaca ctgttcacct ttcgtttttc tcttttaatt tcagtgtgat
4621 aacgggtgaa tatacggtga agggtgaaca gtggattgtt caccttcggg ggatatcggg
4681 ataaaaaaag accggcagat gccggtcagg tgggtcaggc tgttgtaggg tcgtcacatt
4741 ttggcagcca gtcgccgtag ctttcctctt tcagcgtcag gttggtctgt atcccctgtt
4801 tggtatggcg tttctcgtaa ttcagtccgt attccttcag catcaccggc agccccagcc
4861 cgaacatttt cagactgagt acattccggt agccgtttgc ctccatgtag gccagatagg
4921 cgtgatagag gtatttacgg taattgcgcg ggatgatact ggcgttcccc atatacatgc
4981 cgctggtctg cggcagggtt tccagatagc cgataaaatc aaacgtcggg tcggcatccc
5041 gtttgatgtt cagtgcctcg tctgagttct gctgggactg aagcagtgac cgggcgagca
5101 tcgggtcgct gaacttctgc atcaggtgac gcacgatgac cgccagctcg cgggtgattt
5161 tgtccttaag ctgcgggtcg cgctcctgcg gggctatctg ttccgggaag tgaataatca
5221 cccgtcggcg tgacacgccg ccgctgcggt cggtgaagcg catcgggtta ttgttcacgg
5281 ccagaatcac cgccgggatg tgcgtggagt acgcatcccg gtatttcggg tcaacggaca
5341 ccgcatcgcc gccggtgatg gccttgagtc cggcaccgtc gccgctccat ttttcctggt
5401 ccggcaggcg tatcagtgag aagccagtta acgcggcacg ttcacgcggg gattccagcg
5461 tctcgatggt ggccgacgtg gcgttatcct ccccggccag cagggtggct atttcggcca
5521 tgatactttt gccgctgccg ccgggaccgg tcacctccag aaagagctgc cagtcgtagc
5581 ggtttgccag caccataaac agtgcagcca gaatcacgtc gcgtttttcc gcacggccac
5641 ctgcggcacg gtcaagccag cgccagaagg cgggggcgtg ggtttccagc gtttcaccgt
5701 ccaccggcgg ggtgaaatcc acatcgcaca gggtgcgcat ccagtgtgac ggactgtgcg
5761 ggtggaacgt gccgttctgc gtgtcgagca cgccgttacg aaagccaatc aggcggcggg
5821 agggggcttc ctgctgcgga ataatcagct tcagggtgtc caccacggag gccaccttcc
5881 cggaggagaa cggcgcacgc agacgctgaa acagcccggc cacatcccgg gcaaagtcct
5941 gtggcggcag caccttccag acaccatttt catagcggga cagaagctgg ccgttggcat
6001 cgaccgcgag cgcctcgccg taatgctcat agatacgcat ggccttttcg ctggtactca
```

```
6061 tggcggaaaa ctccgcttcg ctcatggtgt cgaacgggct ttcagccggt ggccggatgg
6121 catcgtaaat ggccttacgg gtggcctccc cgccgtactg cgtgaaggca tcattccagt
6181 caccgaagac cggcggcagg gcaacaacac cttcacacgc atctgcggct gcggcggctt
6241 ttttctggcc gtcaccactg aggtcacggt cagcggcaag gacaatctga caggcggggt
6301 gcttctgccg ggcaaggctg gccagagaaa ggaggttcac ggaagaaagc gccaccatca
6361 ccgtttcacc ggtcaggtga tgtacggtaa gtgcggtcgc gtatccctcc gctatccaca
6421 gacgttttcc ggcctgattc tgtccttcaa gggtgtgaca ggtgcccctg acctgtccgc
6481 ctttcagggt gcgcttacgg ccgtcagcac tgattaactg aaggttaacc agttcgccgc
6541 tgtcgtcata cagtggcacc acaaggtcac cggcgcgcca gctcacgcca ccggctctgt
6601 gtgtgccggt cagcatccgg cattcccggc cgggaaagcc cttgcgggtc aggtaggcgt
6661 taccggttcc ggtacgggtt ttcgccatca gggtttgtgc cagtgcggcg gcgttcttcc
6721 gggcagcgtc tgtttcatca acggcggcgg tcgtcactgc cgggtcagcc ggtggcaggc
6781 tgccggtcac ggcagccacc tttgcggccg cgtcggacgg ggaaacacca aaaacctttt
6841 caaccagttt caggccgtca ccggcaccac actgattgca gtaccaggtg ccgcgcccct
6901 ccctgtcatc aaaacggaag cggtcactcc cgccacagac cggacagggc tgatgacggt
6961 tcttcagcac ctgaatcccc agcgccggga gaatacgcgg ccagtggccg agcgcatggc
7021 tgacggtggc ggttacgttc attttcatgg tgttgttctc cttcagtgca gtaccggcgc
7081 ttttatgtga cgggcacaga gttcatccat cacaaccagc ccgagaaagg acagcgacgg
7141 cgcggccttc agggggccgg attccattaa atcttccagc agggcacagg ctatctgacg
7201 ccctttttcc tcaccgtgct ggcgcagata aaagccttcc agctcagcgg cgatggccgc
7261 ctccagtgac tcaagggtga gatgcgggta gcggtgctga cgttcgcaca cggtcagcca
7321 ggcacaggcg acagcgcgac ggtaaagggc agcgcgtaag acgggcggta agggtgtttt
7381 catttgcttt tctccctgtg acagatgact gcattccgtg ccggttgcat taactgataa
7441 ggcatatctg cgtctcctga agacgtgcgt atccctgcgc gaatacgcac atttaatttt
7501 tcgggggtcg ttttttaatt acagataatt gcggtaactg ttatccgggg tggtttccgg
7561 gtcaggctcc gtgcggggaa tttcccgcca ttcccgcgcc accggtgctg cccggctgac
7621 cggaacagtg tcctgcgggt aaatatccag atatttttcc cgccatttct gtaattccgg
7681 gtctccggcc atttctttca gtaccgcatg ccggtttacg gggctgcgtt taaacaggtc
7741 aggacggtca caggtaaatt cccgcagaaa acgccccagc gggatgtctg tggtgcgtcc
7801 gtcagcgagg atacgcacaa ggatactgaa tttacggcgg tacgggttcc agacaatgtc
7861 cgggcagcgg tacggcattt cccacggaat accgtcttcc agaatgccga ccacggccac
7921 atcgggaaaa ccggcagaac ggtaaatctc accgggctgg ggaaaatcaa acatgcgtcc
7981 tgtctccccg gtctttctgc tgggcgagaa aatcgcggca caggcctttg gctttcagct
8041 cattcagcac aaaatcaatc tgaggggctt ttttattacg cacgggacac ctccaccacc
8101 ggcagacggg cagcaaggga gagcacatag tcacggacaa gggaacggcg ggcactgcgt
8161 tcatcaccgg cgacggtgcg aagcatacag atacggggat gacggtctgc gcgacggaca
8221 gccgcaaaca caaagacaaa ttcagggtgt gaggggggtaa gggttgtagc catgatggca
8281 gcctcctgtg aatagcaaat aacgctatcg ccggagttct cacgctcgat ggcgatagcc
8341 cagacggggg tgagaatacc ggcttcacag gataccggcc agcccggagg ctgccccgcc
8401 tgagctacca ttgactctgc ggcataatga gcggacgcgg gcaggatgca cggaatgcca
8461 tctgcacgac tgaccacaca ccacaccata atctggcgct ctgtggcatt gattgcgaca
8521 caaaaaaaga cgcgtggcgc gtcatatgtc gcctgtgaat tgctcgggtt ctcacgcccg
8581 gctgccgatt ttgcggcagc ggaaaaacta tatccgcaaa tgccggaaaa aggcaagcca
8641 gaaaaaggga gtttttgcag agcgggcatc atcatgcgtc gtacccccgt ttgcgtccgg
8701 caatgcgtcc ggccatccat gcggtgactt cagagtgcag ccaggccaca tttttaccgc
8761 caagactcac ctgcggcgga aattcccccct tacggatgag ttcgtagatg gtcgagcgtg
8821 acaggccgca caggtgcatc acttccggca gacgtaaaaa acgctcctgc gtgatgtccg
8881 gcagcggcat cagtggcgtc actggggcgg gagacgggga agaaaaaaca gcttgcatcg
8941 ggctacctcg ttaatgtcca tacagcaccg gataagtccg tccggcttcg ggtagcgctt
9001 tattttatga atattttcag cagacgcaac agggggggatt tgttcaggct gtcttacaat
9061 ggctgtgtgt tttttgttca tctccactta aagtcattta aagccactta aagcaatttg
9121 taatttttat agtgaaatac aaatcgtttc ttcttattca ttcccggcga attaataaaa
9181 acaaacagta gtaaacagca caaaaagccc atcaacgggt gaacagtggt gaacagacgg
9241 tgaacagtca ttactgcgat tgttcaccct ttaacttact gtattactta tctttttttat
9301 taaggtgaac agaggtgaac agtaaaatat aaaaaaacaa acagtaagcc ggtttttcct
9361 gcgacctttt cctggcttgc cggtctgagg atgagtctcc tgtgtcaggg ctggcacatc
9421 tgcaatgcgt cgtgttgttg tccggtgtac gtcacaattt tcttaacctg aagtgacgag
```

```
     9481 gagccggaaa atgtctgacc acactatccc tgaatatctg caacccgcac tggcacaact
     9541 ggaaaaggcc agagccgccc atcttgagaa cgcccgcctg atggatgaga ccgtcacggc
     9601 cattgaacgg gcagagcagg aaaaaaatgc gctggcgcag gccgacggaa acgacgctga
     9661 cgactggcgc acggcctttc gtgcagccgg tggtgtcatg agcgacgagc tgaaacagcg
     9721 ccacattgag cgcgtggcac gccgggagct ggtacaggaa tatgacaatc tggccgtggt
     9781 gctgaatttc gaacgtgaac gcctgaaagg ggcgtgtgac agcacggcca ccgcctaccg
     9841 gaaggcacat catcaccttc tgagtctgta tgcagagcat gagctggaac acgccctgaa
     9901 tgaaacctgt gaggcgcttg tccgggcaat gcatctgagc attctggtac aggaaaatcc
     9961 gctcgccaac accaccggcc atcagggcta cgtcgcaccg gaaaaggctg tcatgcagca
    10021 ggtgaaatca tcgctggaac agaaaattaa acagatgcaa atcagcctca ccggcgagcc
    10081 ggttctccgg ctgaccggac tgtcagcggc aacactcccg cacatggatt atgaggtggc
    10141 aggcacaccg gcacagcgca aggtgtggca ggacaaaata gaccagcagg gagcagagct
    10201 taaggccaga gggctgctgt catgatttac tgtccgtcgt gtggacatgt tgctcacacc
    10261 cgtcgcgcac atttcatgga cgatggcacc aagataatga ttgcacagtg ccggaatatt
    10321 tattgctctg cgacatttga agcgagtgaa agctttttct ctgacagtaa agattcagga
    10381 atggaataca tttcaggcaa acagagatac cgcgattcac tgacgtcagc ctcctgcggt
    10441 atgaaacgcc cgaaaagaat gcttgttacc ggatattgtt gtcggagatg taaaggcctt
    10501 gcactgtcaa gaacatcgcg gcgtctgtct caggaagtca ccgagcgttt ttatgtgtgc
    10561 acggatccgg gctgtggtct ggtgtttaaa acgcttcaga ccatcaaccg cttcattgtc
    10621 cgcccggtca cgccggacga actggcagaa cgcctgcatg aaaaacagga actgccgcca
    10681 gtacggttaa aaacacaatc atattcgctg cgtctggaat gagggctgcc ggttaacacc
    10741 ggccgtcgcc gcacaccgta tttttattct tcagcatgat gagaaagaga taacgatgga
    10801 aagcacagcc ttacagcagg cctttgacac ctgtcagaat aacaaagcag catggctgca
    10861 acgcaaaaat gagctggcag cggccgaaca ggaatatctg cggcttctgt caggagaagg
    10921 cagaaacgtc agtcgcctgg acgaattacg caatattatc gaagtcagaa aatggcaggt
    10981 gaatcaggcc gccggtcgtt atattcgttc gcatgaagcc gttcagcaca tcagcatccg
    11041 cgaccggctg aatgatttta tgcagcagca cggcacagca ctggcggccg cactggcacc
    11101 ggagctgatg ggctacagtg agctgacggc cattgcccga aactgtgcca tacagcgtgc
    11161 cacagatgcc ctgcgtgaag cccttctgtc ctggcttgcg aagggtgaaa aaattaatta
    11221 ttccgcacag gatagcgaca ttttaacgac catcggattc aggcctgacg tggcttcggt
    11281 ggatgacagc cgtgaaaaat tcacccctgc gcagaacatg atttttcgc gtaaaagtgc
    11341 gcaactggca tcacgtcagt cagtgtaaaa ttccccgaaa atccgcccgt ttttactgaa
    11401 aaaagccatg catcgataag gtgcatggct tt
//
```
